# Complementary stable and dynamic prelimbic ensembles encode learned threat value underlying generalization and discrimination

**DOI:** 10.64898/2026.03.08.710406

**Authors:** M.E. Normandin, P.M. Ogallar, M.R. Lopez, M.M. Ramos-Alvarez, I.A. Muzzio

## Abstract

Adaptive behavior requires assigning emotional value to sensory cues and generalizing to similar stimuli. The prelimbic cortex (PL) is critical for threat expression and discrimination, yet its neuronal ensembles undergo substantial turnover over time. How stable population codes emerge despite this reorganization remains unknown. Using longitudinal calcium imaging in freely moving mice, we tracked PL activity for 30 days during auditory discriminative fear learning and retrieval. Despite substantial ensemble turnover, stable graded representations persisted in PL populations. A generalized linear model showed that the tone threat value explained graded activity better than freezing, indicating that PL primarily encodes stimulus significance. Population similarity across stimulus presentations was highest for shock-associated and highly generalized tones, suggesting a stable population code underlying threat generalization. Stable neurons preserved graded gradients across sessions, whereas learning modulated the rate of dynamic frequency-selective neurons. Together, these populations enabled stable threat-value coding despite ongoing ensemble reorganization.

## Introduction

Generalization allows learned responses to extend from a conditioned stimulus to similar cues, whereas discrimination enables animals to distinguish threat from safety. Adaptive behavior requires a balance between these processes, permitting defensive responses to potential danger while minimizing these responses to safe stimuli. Despite the importance of this balance for survival, how neural populations simultaneously support generalization and discrimination remains poorly understood. Two major theoretical frameworks have been proposed to explain these processes. Sensory similarity models posit that generalization arises primarily from overlap in sensory representations (Shepard, 1987). More recent work, however, suggests that sensory similarity alone cannot fully account for generalization. Instead, the brain is proposed to construct mnemonic representations of learned threat value, such that neural activity reflects the estimated probability or magnitude of danger associated with different stimuli (Dunsmoor & LaBar, 2013; Laufer et al., 2016; Verra et al., 2026). Under this framework, sensory similarity allows novel stimuli to be compared with the conditioned stimulus, whereas mnemonic representations acquired through associative learning determine the threat value assigned on the basis of that comparison. These frameworks therefore make distinct predictions about neural activity. Sensory similarity models predict that neural activity is organized according to perceptual similarity, with generalized responses arising from overlapping neural representations and successful discrimination emerging from increasingly distinct representations, sometimes involving separate neuronal populations encoding threat and safety (Grosso et al., 2018). In contrast, threat-value models propose that neural activity reflects the expected danger associated with each stimulus, organizing generalization and discrimination along a continuum of learned threat rather than solely according to perceptual similarity (Levy & Schiller, 2021). In this study, we investigated how the prelimbic cortex (PL), a region implicated in fear expression and threat generalization (Burgos-Robles et al., 2009; Rosas-Vidal et al., 2025; Sierra-Mercado et al., 2011; Sotres-Bayon & Quirk, 2010; Stujenske et al., 2022), supports generalization and discrimination.

Previous studies have established that the prelimbic cortex is required for discriminating between threat and safety cues. During discriminative fear conditioning, transient inhibition of the PL causes animals to freeze similarly to conditioned threat and safe stimuli, resulting in overgeneralization (Likhtik et al., 2014). Likewise, disruption of endocannabinoid signaling in the PL abolishes the neuronal distinction between threat and neutral cues (Rosas-Vidal et al., 2025). Together, these findings indicate that the PL plays a critical role in organizing neural representations that support appropriate discrimination while limiting threat generalization. However, how individual PL neurons encode these computations remains unclear. Human neuroimaging studies have shown that medial prefrontal activity scales with gradients of learned threat, consistent with value-based representations (Lissek et al., 2014). In contrast, other studies suggest that activity in this region primarily reflects the representational similarity of affective or valence information (Chavez & Heatherton, 2015). Adding further complexity, PL neuronal activity is also strongly correlated with freezing behavior—a hallmark defensive response during fear retrieval—regardless of the specific cue predicting an aversive outcome (Casanova et al., 2024; Kyriazi et al., 2020). Consequently, it remains unclear whether PL population activity primarily reflects sensory similarity, learned threat value, or defensive behavior, and how these factors together shape threat generalization and discrimination.

An additional challenge in identifying the cortical neural codes underlying threat generalization and discrimination is that both episodic memory ensembles (DeNardo et al., 2019; Kitamura et al., 2017) and procedural memory ensembles (Do-Monte et al., 2015; Iqbal et al., 2026) undergo substantial reorganization across days—a hallmark of systems consolidation—even while learned behaviors remain remarkably stable. This dissociation between neuronal instability and behavioral persistence suggests that stable behavior can be supported by population-level representations despite continuous changes in their neuronal composition (Deitch et al., 2021; Gallego et al., 2020). Because threat generalization requires evaluating novel stimuli in relation to previously learned representations, often long after the original threat association has been acquired, it may be particularly sensitive to ongoing ensemble reorganization. These observations raise several important questions. Can stable population codes supporting threat generalization emerge despite continuous changes in their neural composition? Do distinct neuronal subpopulations differ in their long-term stability? If so, do stable and dynamic neuronal subpopulations differentially represent learned threat value, sensory similarity, or the expression of defensive behavior?

We hypothesized that the PL contains population- and single-cell representations of learned threat value—defined here as mnemonic representations of the threat or safety significance acquired by conditioned stimuli—that support threat generalization and discrimination. We further hypothesized that the persistence of these representations across retrieval sessions depends on complementary contributions from stable and dynamic neuronal ensembles. Finally, we hypothesized that sensory similarity increases the consistency of population responses across similar sounds, thereby facilitating generalization. To test these hypotheses, we combined longitudinal calcium imaging, computational clustering analyses, and a generalized linear model (GLM) to characterize PL ensemble dynamics during auditory discriminative fear learning and memory retrieval in freely moving mice. Our results indicate that the PL represents learned threat value at both the population and single-neuron levels despite extensive ensemble turnover across retrieval sessions. Population similarity across sound presentations closely tracked behavioral generalization, suggesting that sensory similarity shapes the overlap among neuronal population responses, whereas mnemonic representations of associative learning determine which conditioned stimulus anchors the generalization gradient and defines its direction. Moreover, neuronal activity was more strongly associated with learned threat value than with measured freezing behavior. Together, these findings provide a neural framework for understanding how the PL supports adaptive threat generalization and discrimination.

## Results

### Logarithmic tone separation predicts freezing level

To assess memory retrieval and generalization, GRIN-lens–implanted and non-implanted mice were trained in a differential auditory fear-conditioning paradigm (Fig. 1a). One tone (CS⁺; 80 dB) was paired with a mild foot shock (0.5 mA, 0.5 s), whereas a second tone (CS⁻; 80 dB) was never paired with shock. In one group, the CS⁺ was 15 kHz and the CS⁻ was 3 kHz (CS⁺15; *N* = 27; 7 implanted, 20 non-implanted); in a second group, contingencies were reversed (CS⁺3; *N* = 22; 5 implanted, 17 non-implanted). A no-shock control group was exposed to the same tones without shock (*N* = 13; 6 implanted, 7 non-implanted).

**Figure 1.**
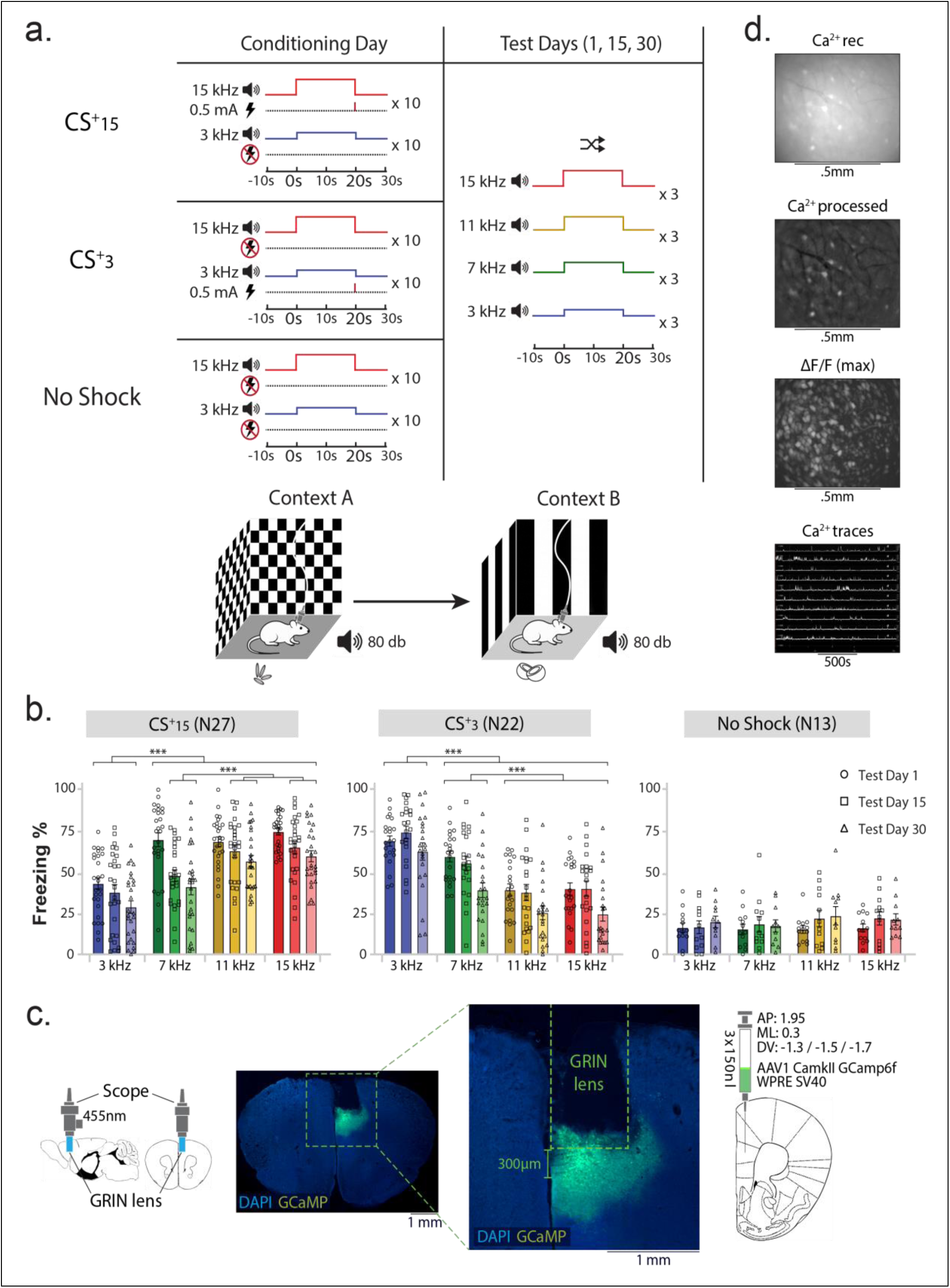
Experimental design, behavioral responses, viral strategy, and calcium imaging. **a.** Experimental design illustrating discriminative fear conditioning with either a 15-kHz CS+ or a 3-kHz CS+ (conditioned groups) and no-shock controls, which were tested at the same time points but never received footshocks. All groups were tested with 3- and 15-kHz tones and two intermediate frequencies (7 and 11 kHz) on days 1, 15, and 30 after conditioning. **b.** Behavioral responses across retrieval sessions (CS+15 group: *n* = 27 mice; CS+3 group: *n* = 22 mice; no-shock controls: *n* = 13 on days 1 and 15 and *n* = 11 on day 30). Two control mice lost their lenses after day 15 and therefore did not contribute data to the day 30 session. The repeated-measures analysis included the 11 control mice with complete data across all sessions. Two-way repeated-measures ANOVA. CS+15 group: main effect of day, *F*(2,52) = 13.28, *p* < 0.001; main effect of frequency, *F*(3,78) = 85.09, *p* < 0.001; and day × frequency interaction, *F*(6,156) = 3.79, *p* = 0.002. CS+3 group: main effect of day, *F*(2,42) = 14.42, *p* < 0.001; main effect of frequency, *F*(3,63) = 58.81, *p* < 0.001; and day × frequency interaction, *F*(6,126) = 1.41, *p* = 0.217. No-shock controls: main effect of day, *F*(2,20) = 1.38, *p* = 0.276; main effect of frequency, *F*(3,30) = 1.43, *p* = 0.253; and day × frequency interaction, *F*(6,60) = 0.335, *p* = 0.916. Significant pairwise comparisons following Tukey’s multiple-comparisons test are denoted by asterisks: \*\*\**p* < 0.001. **c.** Viral strategy and representative histological section showing GCaMP6f expression. Imaging depth did not exceed 300 μm, restricting recordings to the PL. **d.** Representative calcium-imaging data showing a raw fluorescence frame, processed frame, maximum-intensity projection across frames, and corresponding activity traces.

Retrieval was assessed 1, 15, and 30 days after conditioning to probe recent, long-term, and remote memory (Bontempi et al., 1996). During each retrieval session, mice were tested in a novel context with the CS⁺, the CS⁻, and two intermediate frequencies (7 and 11 kHz), presented in a semi-random order with each tone repeated three times (Fig. 1a). The intermediate frequencies were selected to evaluate whether perceptual similarity predicts threat generalization. Although the separation between the CS⁺ and CS⁻ was identical in both groups (2.32 octaves), the spacing between the CS⁺ and the nearest intermediate frequency differed (0.45 octaves for 11–15 kHz versus 1.22 octaves for 3–7 kHz). Thus, if threat generalization is determined primarily by sensory similarity, freezing should generalize more strongly to 11 kHz in CS⁺15 mice than to 7 kHz in CS⁺3 mice.

In the CS⁺15 group, mice consistently discriminated between the CS⁺ and CS⁻. Although generalization to intermediate frequencies was broad on day 1, it became progressively restricted to 11 kHz—the frequency adjacent to the CS⁺—during long-term (day 15) and remote (day 30) retrieval (p < 0.05; Fig. 1b, left). CS⁺3 mice also discriminated between the CS⁺ and CS⁻; however, these animals exhibited greater freezing to the CS⁺ than to all intermediate frequencies across testing days (p < 0.05; Fig. 1b, center), indicating reduced generalization to the adjacent tone when the logarithmic separation was larger. No-shock control mice showed no significant differences in freezing across frequencies on any testing day (p > 0.05; Fig. 1b, right), confirming that the graded freezing patterns resulted from associative learning rather than the acoustic properties of the tones. Together, these results indicate that logarithmic spacing (i.e., frequency similarity), rather than simple proximity to the CS⁺, was the primary determinant of generalization.

Because our experimental design involved repeated retrieval testing over the course of a month, we examined whether repeated tone presentations produced extinction. We quantified stimulus discrimination using discrimination ratios (DRs), which measure discrimination between each tone and the CS+. As shown in Fig. S1, DRs increased over successive retrieval sessions in both conditioned groups. In CS+15 mice, significant increases were observed for the 3 and 7 kHz tones (*p* < 0.05), whereas persistent generalization remained only for the 11 kHz tone. Similarly, CS+3 mice exhibited significant increases in DRs for the CS− and all intermediate frequencies (*p* < 0.05, Fig. S1). Thus, repeated testing progressively sharpened discrimination rather than reducing conditioned responding. This pattern is inconsistent with extinction, which would be expected to flatten discrimination gradients, and instead agrees with previous studies showing that discrimination training narrows behavioral generalization gradients (Dunsmoor & LaBar, 2013; Herzog et al., 2021; Jenkins & Harrison, 1960; Lommen et al., 2017).

No sex differences were observed in CS⁺15 mice (11 females, 16 males) or (6 females, 7 males; *p* > 0.05). In CS⁺3 mice (11 females, 11 males), minor and inconsistent sex-related effects were detected on days 1 and 15, but these effects were absent by day 30 (Fig. S2). Accordingly, sex was not included as a factor in subsequent neural analyses. Behavioral performance did not differ between implanted and non-implanted mice on days 1 and 15 (*p* > 0.05). On day 30, a modest main effect of group was observed in the CS⁺3: group, reflecting higher overall freezing levels in implanted animals compared with non-implanted mice (*p* < 0.03). Importantly, both experimental groups exhibited robust frequency effects across days (*p* < 0.001), with no group × frequency interactions (*p* > 0.05), indicating preserved threat discrimination and generalization profiles.

### PL sound-responsive ensembles exhibit dynamic properties across retrieval sessions

To examine the temporal evolution of PL neuronal responses, we performed longitudinal calcium imaging in CS⁺15 (N = 7), CS⁺3 (N = 5), and no-shock control mice (N = 6). Across groups, neurons were tracked during conditioning and all retrieval sessions. To assess registration quality, we used the conditioning session as a reference and measured the centroid shifts between matched neurons across retrieval sessions. For each animal, we generated histograms of these shifts (µm) and compared them with the median equivalent circular footprint diameter (Fig. S3). Across animals, 99–100% of registered neurons exhibited centroid shifts smaller than the median footprint diameter, indicating highly reliable cell registration. The proportion of registered neurons was equivalent across groups (Fig. S3). Fig. 1c illustrates the GCaMP6f viral injection strategy and representative histological sections showing the pattern of viral expression. Fig. 1d shows a representative raw calcium video frame, processed frames, and activity traces.

Sound-evoked responses were visualized using activity plots with calcium activity ranked across simultaneously recorded neurons (Fig. 2a). Across animals, we visualized distinct neuronal subpopulations showing positive modulation, negative modulation, mixed responses, or no consistent response to sound in each session (Fig. 2b). Sound-responsive neurons were identified using a sound responder statistical test that detected activity modulation relative to baseline, allowing reliable identification of sound responses while remaining robust to noise. This approach was effective for neurons exhibiting either brief calcium transients that rose and decayed rapidly or sustained activity throughout the tone presentation. Fig. 2c summarizes the proportions of response types pooled across animals and sessions.

**Figure 2.**
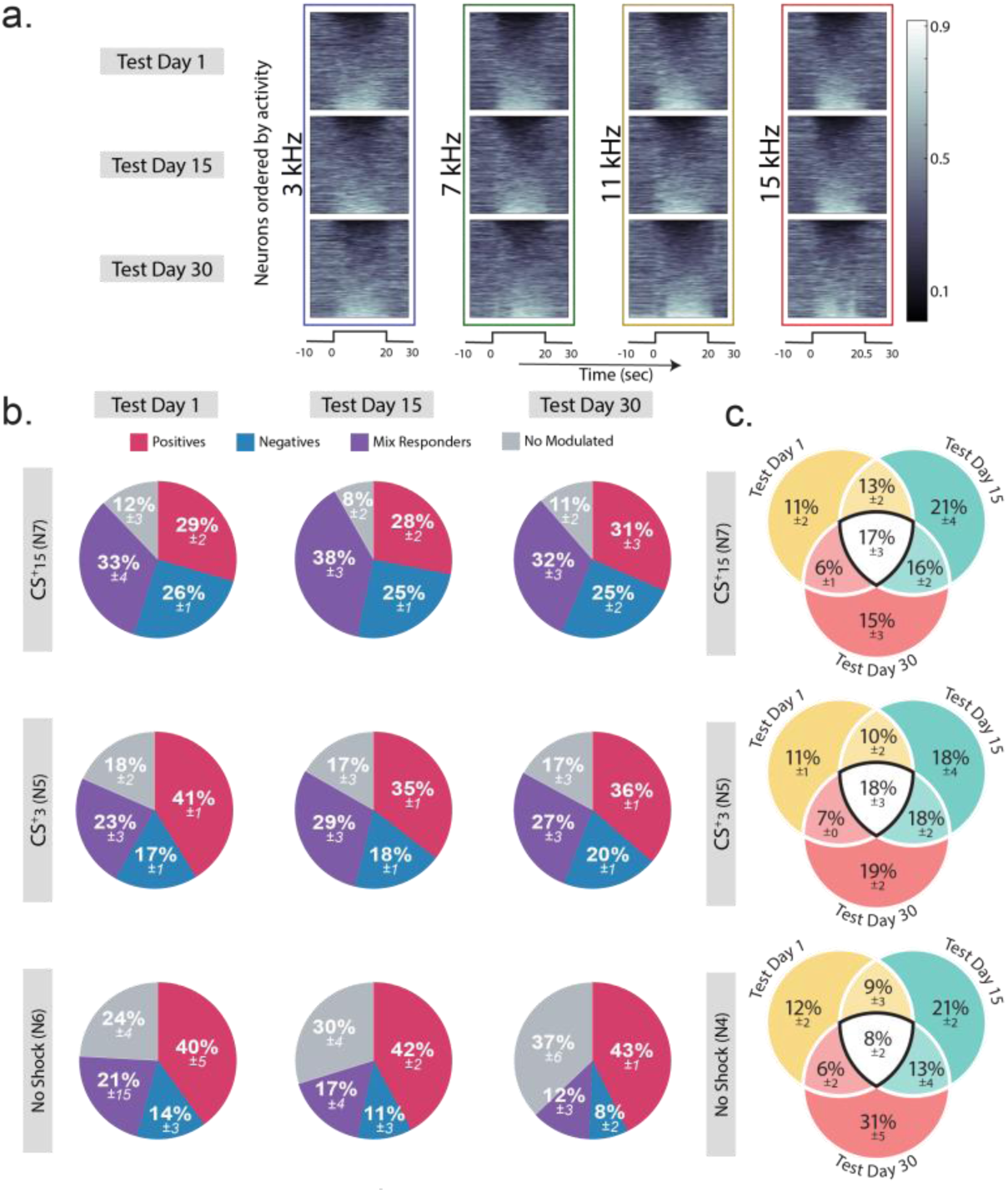
Distribution of stable and dynamic cell types across sessions. **a.** Activity plots from a representative animal conditioned with a 15-kHz CS+, with neurons ordered by activity level. The plots show positive tone-responsive neurons (bottom), negative tone-responsive neurons (top), and mixed or nonresponsive neurons (middle) before and after tone presentation. **b.** Proportions of neuronal response types across experimental groups and sessions. Numbers of cells—CS+15 group: conditioning, 3,834; day 1, 3,177; day 15, 4,186; and day 30, 3,693. CS+3 group: conditioning, 1,628; day 1, 1,381; day 15, 1,881; and day 30, 1,833. No-shock controls: conditioning, 1,118; day 1, 986; day 15, 1,386; and day 30, 563. Two control mice yielded data only through day 15. **c.** Venn diagrams showing the proportions of consistently active neurons (active during all three retrieval sessions), partially active neurons (active during two sessions), and transiently active neurons (active during only one session). The proportions of neurons in these categories were similar across groups, except that transient neurons on day 30 were more abundant in the control group. This difference reached significance only between the control and CS+15 groups (*p* < 0.05; Table S1). Diagrams and analyses for the control group included only mice recorded through day 30.

Given the ongoing debate over whether neocortical memory ensembles stabilize or remain dynamic over time (DeNardo et al., 2019; Kitamura et al., 2017; Kupke & Oliveira, 2025; Lopez et al., 2024; Mau et al., 2020; Rao-Ruiz et al., 2021; Refaeli et al., 2023; Terranova et al., 2023; Zaki & Cai, 2024), we next assessed the temporal stability of PL ensembles by tracking the cellular footprints of sound-responsive neurons across retrieval sessions. Ensemble stability was visualized using Venn diagrams (Fig. 2c). A moderate proportion of neurons was active across all retrieval sessions, with no differences between groups (p > 0.05). Likewise, there were no group differences in the proportion of neurons overlapping across only two sessions (p > 0.05). For neurons active in only a single session, control mice exhibited a higher proportion on day 30 compared with CS15+ mice (p < 0.05, Table S1), with no other differences observed. Overall, these data indicate that the majority of neurons overlapped across only a limited number of sessions or were transiently active, reflecting pronounced population-level dynamism across retrieval sessions.

### Sound-modulated PL population responses encode generalization gradients

To assess population-level representations in response to conditioned and novel tones, calcium activity was averaged across three presentations for each frequency during a 10 -s baseline, 20-s tone presentation, and 10-s post-stimulus interval. In CS⁺15 mice, positively modulated sound-responsive neurons exhibited graded tone activity reflecting learned contingency value across testing days (Fig. 3a, top). Area-under-the-curve (AUC) analyses obtained by averaging recorded cells per animal revealed the highest responses at 15 and 11 kHz, intermediate responses at 7 kHz, and the lowest at 3 kHz; with no difference between 11 and 15 kHz (*p* > 0.05) across testing days. This population level gradient mirrored behavioral generalization (Fig. 1b, left). By contrast, negatively modulated neurons showed overlapping responses across frequencies with no significant differences at any test session (*p* > 0.05; Fig. 3a, bottom). A complementary pattern was observed in CS⁺3 mice, with graded responses that peaked at 3 kHz (Fig. 3b). As in CS⁺15 mice, negatively modulated neurons exhibited overlapping responses across frequencies and days (*p* > 0.05). In no-shock controls, although both positive and negative responses were present, population activity was not modulated by tone frequency (*p* > 0.05; Fig. 3c, bottom panel), indicating that neuronal graded responses require associative learning.

**Figure 3.**
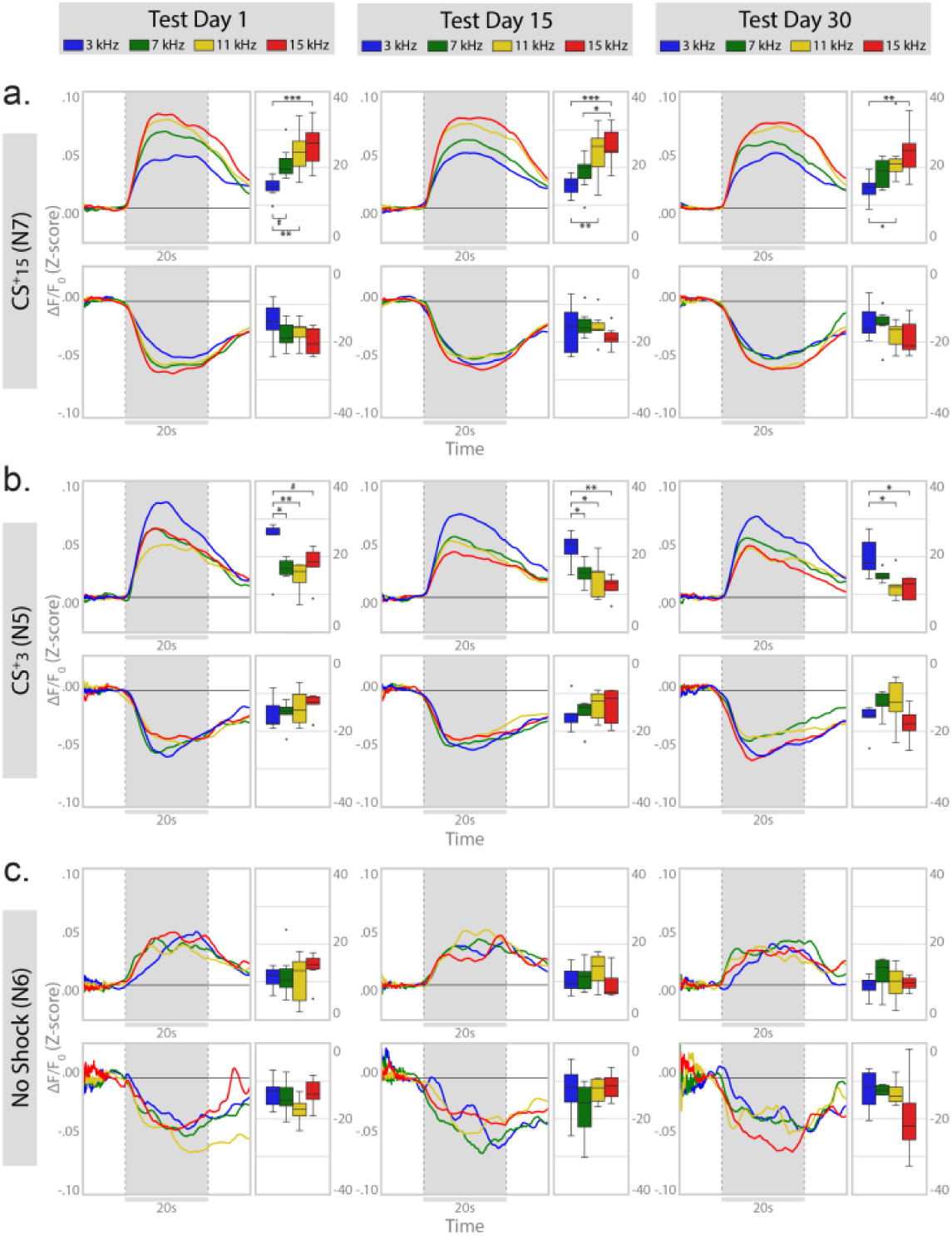
Population activity of positive tone-responsive neurons exhibits graded patterns reflecting learned threat value. **a–c.** Population responses in animals conditioned with a 15-kHz CS+ (**a**), animals conditioned with a 3-kHz CS+ (**b**), and no-shock controls (**c**). In each group, the upper panels show positive tone-responsive neurons, and the lower panels show negative tone-responsive neurons. Adjacent boxplots show the areas under the population response curves (AUCs). Boxplots indicate the median (center line) and interquartile range (box); whiskers extend to the most extreme values within 1.5 times the interquartile range, and points beyond the whiskers represent individual outliers. CS+15 group—positive responders: day 1, *F*(3,18) = 9.963, *p* < 0.001; day 15, *F*(3,18) = 9.973, *p* < 0.001; and day 30, *F*(3,18) = 6.627, *p* = 0.003. Negative responders: day 1, *F*(3,18) = 2.483, *p* = 0.094; day 15, *F*(3,18) = 1.877, *p* = 0.178; and day 30, *F*(3,18) = 2.753, *p* = 0.073. CS+3 group—positive responders: day 1, *F*(3,12) = 6.899, *p* = 0.006; day 15, *F*(3,12) = 9.247, *p* = 0.002; and day 30, *F*(3,12) = 6.123, *p* = 0.009. Negative responders: day 1, *F*(3,12) = 0.870, *p* = 0.484; day 15, *F*(3,12) = 0.512, *p* = 0.641; and day 30, *F*(3,12) = 1.448, *p* = 0.278. No-shock controls—positive responders: day 1, *F*(3,15) = 0.527, *p* = 0.670; day 15, *F*(3,15) = 1.852, *p* = 0.181; and day 30, *F*(3,9) = 1.046, *p* = 0.418. Negative responders: day 1, *F*(3,13) = 1.205, *p* = 0.347; day 15, *F*(3,13) = 1.375, *p* = 0.294; and day 30, *F*(3,9) = 0.950, *p* = 0.457. Asterisks denote significant Tukey post hoc comparisons: \**p* < 0.05, \*\**p* < 0.01, and \*\*\**p* < 0.001.

To determine how AUC varied across groups over retrieval sessions, we performed a three-way mixed-effects ANOVA with group (CS+15, CS+3, and no shock), frequency (3, 7, 11, and 15 kHz), and test days (1, 15, and 30) as factors, with repeated measures on frequency and test days. For positive responder neurons, the analysis revealed significant main effects of group (p < 0.001) and test days (p < 0.05), as well as a significant group × frequency interaction (p < 0.001), whereas the test days × frequency and group × test days × frequency interactions were not significant (p > 0.05; Table S2a). Tukey-corrected post hoc comparisons showed that, in the CS+15 group, AUC differed between all frequency pairs except 11 and 15 kHz (p < 0.05). In the CS+3 group, the AUC at 3 kHz differed from those at 7, 11, and 15 kHz (p < 0.05), whereas no significant frequency differences were observed in the no-shock controls (p > 0.05). For negative responder neurons, the only significant effect was a test days × frequency interaction (p < 0.01). Tukey-corrected simple-effects analyses revealed that, on day 30, the AUC at 15 kHz differed from those at 3, 7, and 11 kHz (p < 0.05; Table S2b). Because this pattern was observed across all experimental groups, including the no-shock controls, it is unlikely to reflect associative learning. These results indicate that although the AUC exhibited modest changes over retrieval sessions, these changes were not group-specific and therefore do not support learning-dependent alterations in neuronal responses. Together, these results show that despite substantial neuronal turnover, PL population responses encode generalization gradients, closely matching behavioral expression.

### Consistently active neurons preserve stable generalization gradients, whereas newly recruited neurons refine them over retrieval sessions

Longitudinal tracking of individual neurons revealed how subpopulations with distinct stability profiles shape the evolution of representations that parallel behavioral expression. We compared population responses across three stability types: consistently active neurons (active during conditioning and all retrieval sessions), emerging–retained neurons (recruited after conditioning and persisting through day 30), and transiently active neurons (only active during a single retrieval session). Consistently active, positively modulated neurons exhibited graded population responses reflecting a generalization gradient from day 1 onward in both experimental groups, which was quantified by calculating AUC per animal (*p* < 0.05; Fig. S4a–b, Table S3). In contrast, negatively responding neurons showed no graded tuning across days (*p* > 0.05), except for a day 1 difference between 15 and 3 kHz in CS⁺15 mice (*p* < 0.05). In control mice, positively and negatively modulated neurons displayed variable and inconsistent activity across frequencies in all cell categories (consistently active, emerging–retained, and transiently active). As a result, these neurons were not included in further cell-type analyses.

Emerging-retained neurons recruited after conditioning exhibited graded population responses when recruitment occurred after day 1 (Fig. S5a–b, Table S3). This category encompassed neurons that emerged on day 1 and persisted through day 30, as well as neurons that emerged on day 15 and remained active through day 30. In both experimental groups, graded tuning was absent on day 1 (p > 0.05) but emerged on days 15 and 30, when population responses closely mirrored behavioral generalization, with responses to 11 and 15 kHz becoming increasingly similar regardless of CS⁺ identity (*p* > 0.05). By contrast, emerging–retained negatively responding neurons showed no significant emotional tuning across days.

Finally, we examined transiently active neurons, defined as cells active only on a single retrieval session. Transiently positive responders showed no valence tuning when active on day 1 but exhibited clear graded population responses when recruited on days 15 or 30 in both experimental groups (Fig. S6a–b, Table S3). In contrast, transiently negative responders showed no significant frequency tuning at any test day (*p* > 0.05, Table S3). Together, these results indicate that consistently active neurons maintain stable generalization gradients across retrieval, whereas neurons recruited after conditioning progressively sharpen their selectivity over test days.

### Tone identity is a stronger predictor of PL activity than freezing

Previous studies have shown that PL activity is associated with fear expression ((Burgos-Robles et al., 2009; Sierra-Mercado et al., 2011; Sotres-Bayon & Quirk, 2010), and more recent work has demonstrated that freezing can be decoded from PL neuronal activity (Casanova et al., 2024; Kyriazi et al., 2020). During fear retrieval and generalization, tones with greater learned threat value also tend to evoke more freezing. This covariation makes it difficult to determine whether PL activity primarily reflects the learned significance of the tones or simply tracks freezing behavior. Our experimental design provided leverage to partially dissociate these possibilities. The same auditory stimuli were presented across sessions and experimental groups, but their assignments as the CS+ and CS− were reversed between groups, and freezing varied substantially across tones and retrieval sessions. Moreover, continuous acceleration measurements obtained from the miniendoscope enabled probabilistic estimates of freezing without imposing an arbitrary threshold. To quantify the independent contributions of tone identity and freezing, we fitted a generalized linear model (GLM) to each neuron’s calcium activity, using tone identity and freezing probability as predictors.

We analyzed positively and negatively responsive neurons separately in the CS+15, CS+3, and control groups. For positively responsive neurons, freezing-corrected tone regression coefficients (β) exhibited opposite monotonic gradients that peaked at the CS+ in the two conditioned groups, whereas controls showed relatively flat coefficients (Fig. 4a; effect of groups: p < 0.01, group x frequency interaction: p < 0.001, Table S4a). To compare the relative contributions of tone identity and freezing, we calculated the freezing-to-tone β ratio. Across all groups, this ratio remained below 1 (Fig. 4b, Table S4b), indicating that tone identity contributed more strongly than freezing to neuronal activity. Within-cell comparisons showed that the median absolute β coefficient for tone identity was significantly greater than that for freezing for all groups (p < 0.001, Fig. 4c, Table S4c). Consistent with this observation, only a modest fraction of neurons was classified as freezing-dominant (Fig. S7a), and this proportion remained stable across retrieval sessions, with no significant differences between groups (p > 0.05; Fig. 4d; Table S4d).

**Figure 4.**
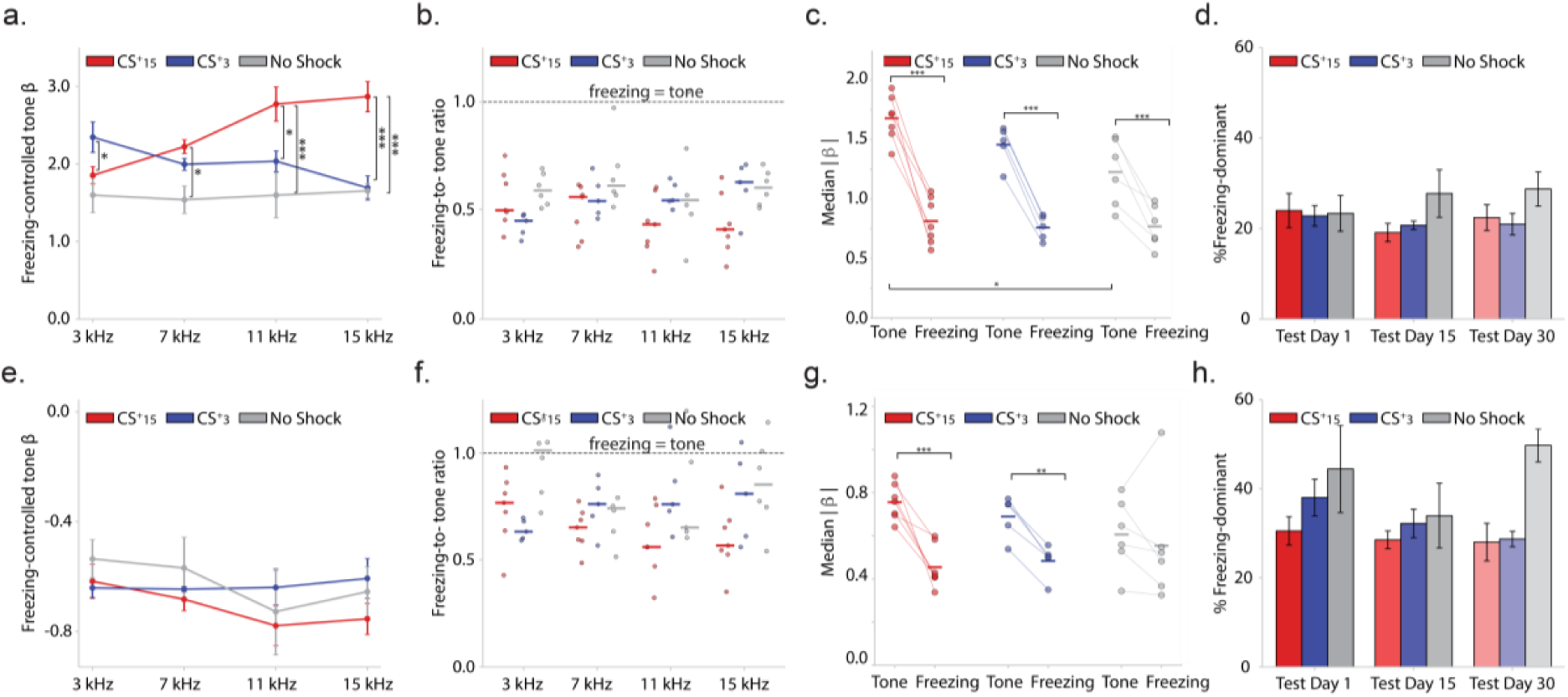
Generalized linear model (GLM) analysis of the relative contributions of tone identity and freezing behavior to neuronal activity for positive (a-d) and negative (e-h) responder cells. **a.** Regression coefficients (β) for positive tone-selective neurons in conditioned mice exhibit opposite monotonic gradients across tones after controlling for freezing. This graded pattern is absent in no-shock controls. **b.** The freezing-to-tone coefficient ratio remains below 1 for every tone in all groups, indicating a greater contribution of tone identity than freezing to the activity of positive tone-selective neurons. **c.** Within-cell comparisons confirm that the absolute median β coefficients for tone exceed the absolute freezing coefficients in all groups. **d.** When pooled across tones, the proportion of freezing-dominant neurons remains stable across retrieval sessions. **e.** Negative tone-selective neurons exhibit suppressive tone coefficients across all frequencies, with only weak frequency dependence, indicating the absence of pronounced graded responses. **f.** Freezing-to-tone coefficient ratios are higher than those observed in positive responders but remain below 1 across tones in conditioned mice. In no-shock controls, the ratio approaches 1, indicating similar contributions of freezing and tone identity. **g.** Within-cell comparisons confirm that the absolute tone coefficients exceed the absolute freezing coefficients, but only in the conditioned groups. **h.** The proportion of freezing-dominant neurons remains relatively stable across retrieval sessions in each group. Controls displayed a slightly higher proportion of freezing-dominant neurons, but this difference was not significant. Significant Tukey multiple comparisons are denoted by asterisks: \**p* < 0.05, \*\**p* < 0.01, and \*\*\**p* < 0.001.

Negatively responsive neurons displayed a different pattern. Tone β coefficients showed little evidence of graded tuning (p > 0.05; Table S4e; Fig. 4e), consistent with the population responses shown in Fig. 3. Although tone identity remained the stronger predictor overall, freezing contributed proportionally more than in positively responsive neurons, resulting in higher freezing-to-tone β ratios (Fig. 4f, Table S4f). However, the absolute β coefficient for tone identity was higher than that for freezing in the conditioned groups (p > 0.01, Fig. 4g, Table S4g). There were no differences in the proportion of freezing dominant negative responders between groups across tones (p > 0.05, Fig. S7b) and these proportions remain stable across retrieval days (p > 0.05, Table S4h). Together, these findings demonstrate that PL population activity is better explained by tone identity than by freezing behavior, particularly among positively responsive neurons.

### Population vector similarity at stimulus onset determines degree of generalization

The preceding results indicate that PL population activity is organized along generalization gradients, such that perceptually similar tones (i.e., adjacent frequencies) elicit similar population responses. However, the neural mechanisms that calibrate the similarity relationships remain unclear. Previous studies have shown that consistent neuronal responses across repeated stimulus presentations support stable population representations that facilitate generalization to similar cues (Hoshi et al., 2023). To assess population similarity within each retrieval session, we compared population activity across the three repeated presentations of each tone. Population activity was estimated using CASCADE, a validated spike-inference neural network (Rupprecht et al., 2021), and activity vectors were constructed from simultaneously recorded neurons. Response similarity between tone pairs was then quantified during stimulus presentation to assess the consistency of population activity across repeated trials.

Rate-based population similarity analyses revealed reliable, temporally structured responses for 3/3 kHz tone pairs in CS⁺3 mice and for 15/15 and 15/11 kHz tone pairs in CS⁺15 mice. In control mice high population similarity was absent for any frequency pair (Fig. 5a-d). Because population similarity peaked shortly after stimulus onset, we quantified similarity during the first 5 s after tone onset relative to the CS⁺. In CS⁺15 mice, early population similarity was highest for 15/15 and 15/11 tone pairs (*p* < 0.001), with no difference between them (*p* > 0.05), and was significantly greater than for 15/3 comparisons (*p* < 0.05, Fig. 5e). In CS⁺3 mice, early similarity was highest for 3/3 tone pairs (*p* < 0.004) and significantly lower for 11/3 and 15/3 comparisons (Fig. 5f; *p* < 0.05). The 3/3 and 3/7 comparisons showed a non-significant trend (*p* = 0.08). No differences were observed among 7/15, 11/15, and 15/15 tone pairs in either group (*p* > 0.05). These findings indicate that population-level similarity at stimulus onset scales with behavioral threat generalization.

**Figure 5.**
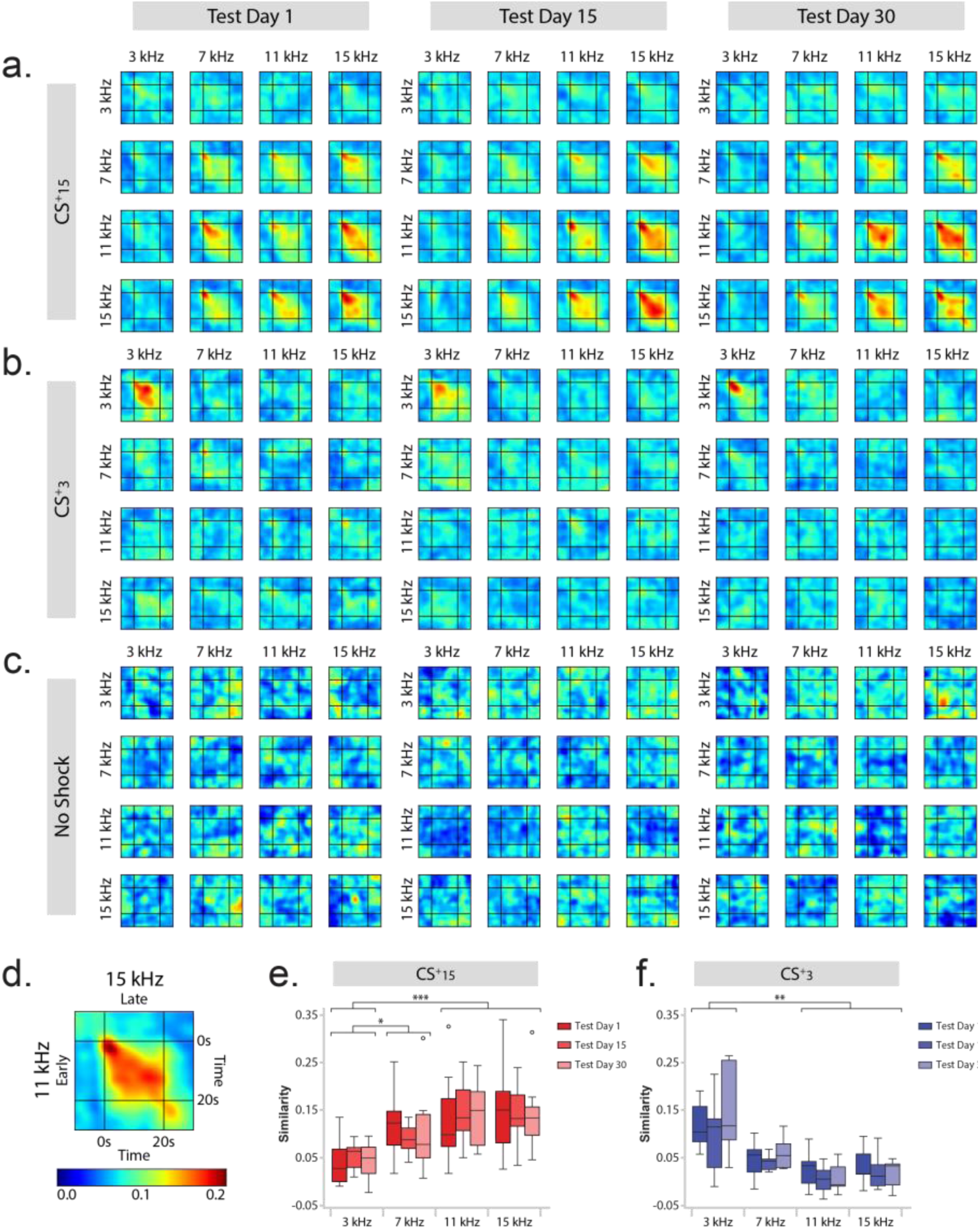
Population-vector similarity across tones and time. **a–c.** Population-similarity maps for all tone pairs across the time course of tone presentation in the CS+15 (**a**), CS+3 (**b**), and no-shock control (**c**) groups. **d.** Schematic illustrating the color scale and orientation of the similarity maps. The y-axis represents the earlier tone in each comparison, and the x-axis represents the later tone. **e, f.** Boxplots showing population similarity during the first 5 s after tone onset, quantified relative to the CS+, for the CS+15 (**e**; *F*(3,36) = 12.025, *p* < 0.001) and CS+3 (**f**; *F*(3,24) = 7.435, *p* = 0.004) groups. Asterisks denote significant Tukey post hoc comparisons: \**p* < 0.05, \*\**p* < 0.01, and \*\*\**p* < 0.001.

In summary, PL population responses formed generalization gradients that paralleled behavioral generalization and were primarily organized according to tone learned threat value. The overlap among responses to conditioned and novel tones was consistent with their perceptual similarity, whereas repeated nonreinforced testing was accompanied by progressive sharpening of both behavioral discrimination and the response gradients expressed by neurons recruited after conditioning. Together, these findings provide a potential neural framework through which stored threat associations may guide the evaluation of perceptually similar cues.

### Different subpopulations encode acoustic versus learned properties of sound association

Our previous analyses demonstrated that generalization gradients are represented at the population level. However, these findings do not reveal how these representations arise. Specifically, the observed population gradients could emerge either from the pooled activity of frequency-selective neurons that respond to individual tones or from neuronal subpopulations that integrate information across tones to encode their learned threat-value. To investigate these possibilities, we developed a clustering approach based on mutual information (MI), which captures both linear and nonlinear relationships between neuronal response profiles (Quian Quiroga & Panzeri, 2009). MI was used to group neurons with related activity patterns into functional subpopulations. Because MI does not preserve response polarity, neurons were subsequently classified by the sign of their pairwise correlations and re-clustered, consistently revealing two major response classes: positively and negatively modulated sound responders (Fig. 6a). Clustering was performed across all animals to derive global cluster identities and within each experimental and control group to identify learning-specific subpopulations (Figs. S7–S9).

**Figure 6.**
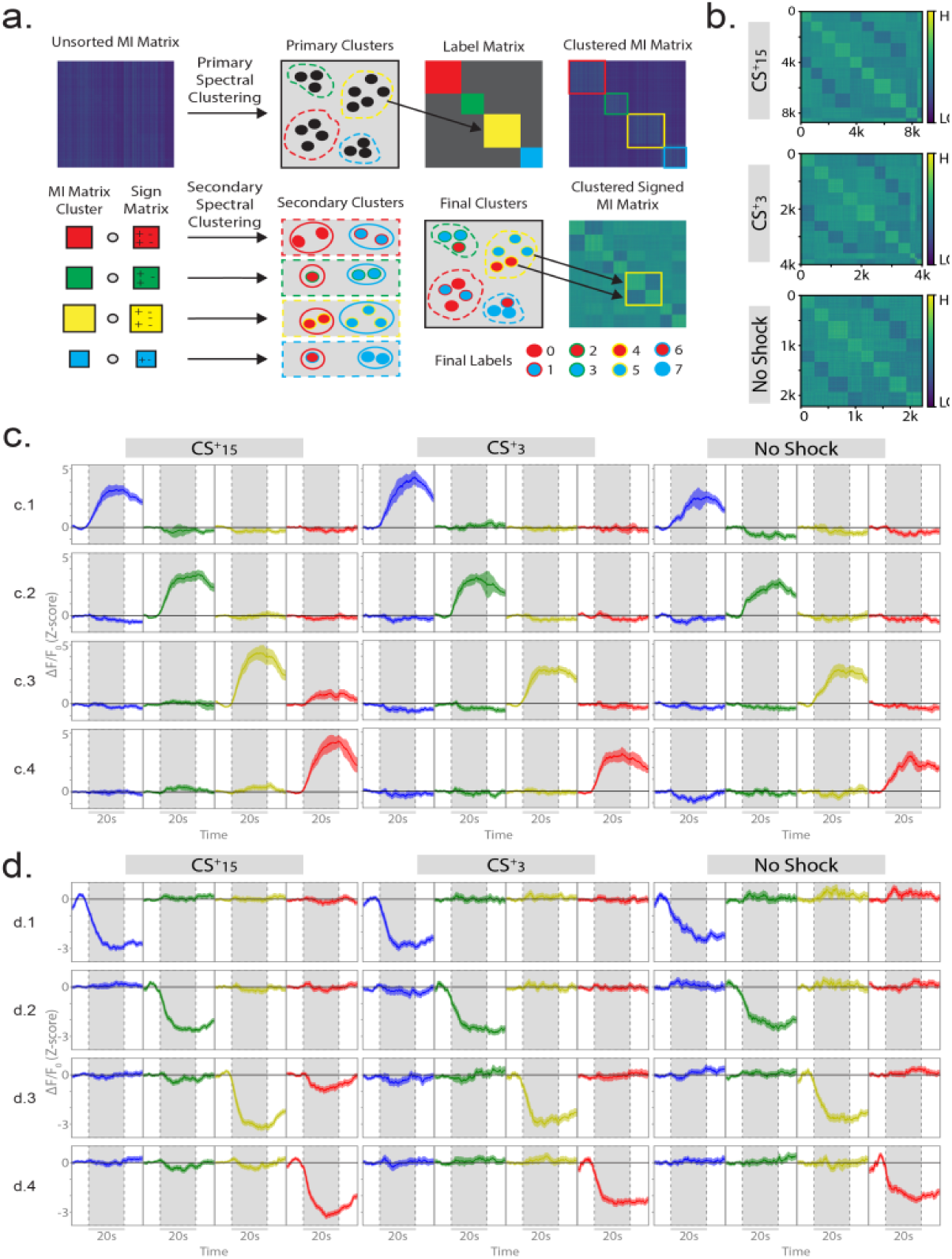
Signed mutual information (MI) clustering identifies PL subpopulations with distinct tone-response profiles. **a.** MI-based clustering pipeline. An MI matrix generated from simultaneously recorded PL neurons was spectrally clustered according to shared information, independently of response sign. Within each primary cluster, MI values were combined with the signs of pairwise correlations and reclustered to identify subclusters with distinct positive or negative activity patterns. Subcluster labels were then combined to generate the final signed MI matrix. **b.** Clustered signed MI matrices for the CS+15 (top), CS+3 (middle), and no-shock control (bottom) groups, with neurons sorted by cluster label. Color indicates the sign and magnitude of MI. **c.** Mean stimulus-aligned responses of clusters positively modulated by individual tones. **c.1–c.4** correspond to 3, 7, 11, and 15 kHz, respectively. **d.** Mean stimulus-aligned responses of clusters negatively modulated by individual tones. **d.1–d.4** correspond to 3, 7, 11, and 15 kHz, respectively.

Clustering quality was assessed by sorting signed MI values by global cluster identity and comparing within-versus across-cluster correlations (Fig. 6b). The resulting box-diagonal structure indicated robust clustering, with comparable quality indices across groups (CS⁺15: 0.28; CS⁺3: 0.27; controls: 0.30). In all cases, within-cluster correlations exceeded across-cluster correlations, supporting the reliability of the approach (CS⁺15: 0.30 vs. −0.03; CS⁺3: 0.27 vs. −0.02; controls: 0.28 vs. −0.04).

Across all groups, we identified positively and negatively modulated clusters selective for individual frequencies, indicating that frequency-specific representations are present in PL independent of learning (Fig. 6c–d; Fig. S8–S10), consistent with a role in auditory processing (Hockley & Malmierca, 2024; Zikopoulos & Barbas, 2006). In contrast, clusters encoding graded patterns— characterized by neurons responding to all frequencies in a graded manner—were observed exclusively in trained animals (Fig. 7). Graded neurons peaking at 15 kHz were specific to the CS⁺15 group (Fig. 7a–b; Fig. S8), whereas neurons peaking at 3 kHz were unique to the CS⁺3 group (Fig. 7c–e; Fig. S9).

**Figure 7.**
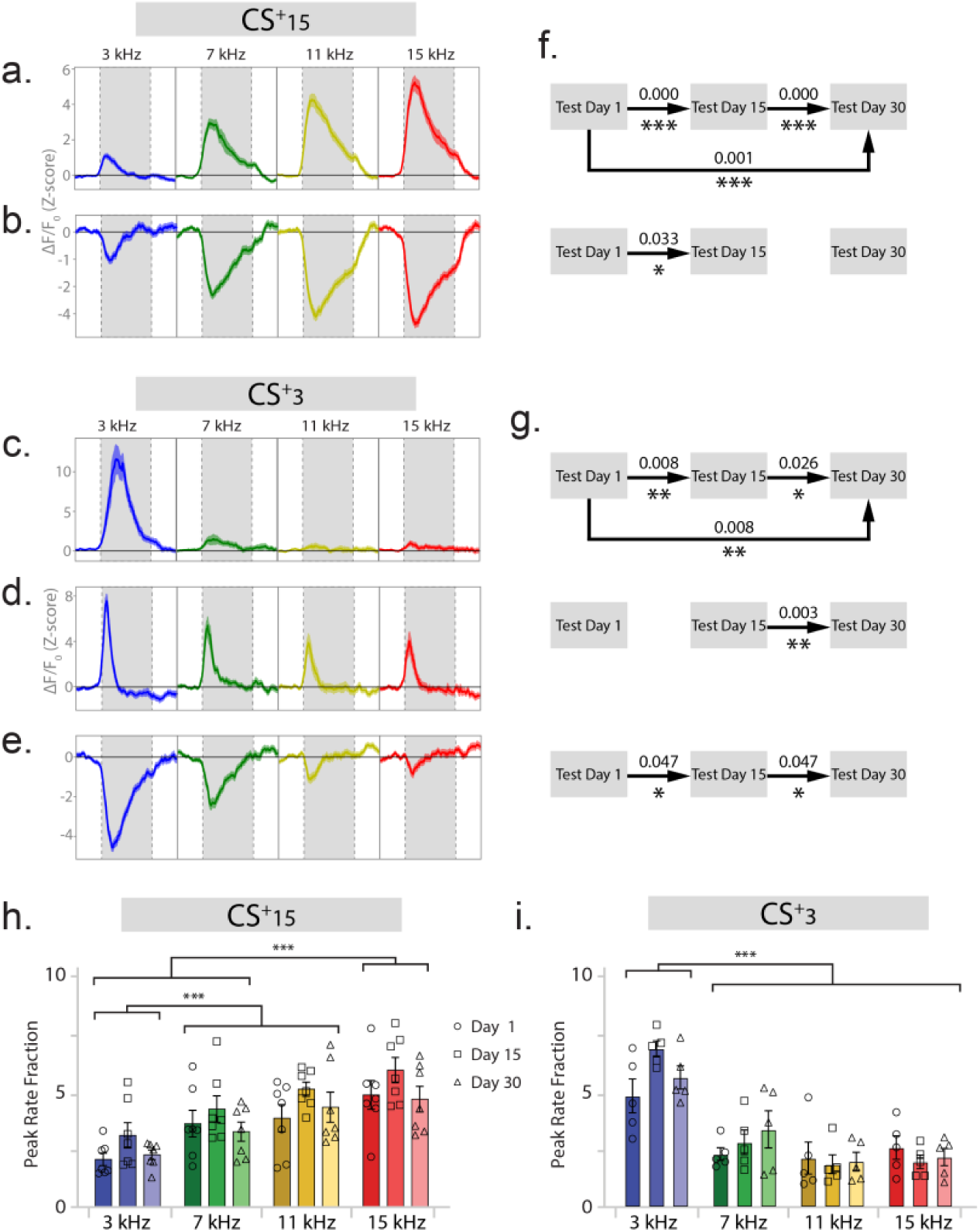
Graded tone-responsive neurons are present only in conditioned groups. **a, b.** Mean stimulus-aligned population responses of clusters exhibiting graded tone responses in animals conditioned with a 15-kHz CS+, showing positive (**a**) and negative (**b**) response patterns. **c–e.** Mean stimulus-aligned population responses of clusters exhibiting graded tone responses in animals conditioned with a 3-kHz CS+, showing positive (**c, d**) and negative (**e**) response patterns. **f, g.** Stability of neuronal identity within graded tone-responsive clusters across retrieval sessions. Asterisks denote significance after Benjamini–Hochberg correction. **h, i.** Baseline-to-stimulus firing-rate ratios (BSRs) showing changes in the activity of graded clusters across retrieval sessions in the CS+15 (**h**) and CS+3 (**i**) groups. Graded clusters were not identified in no-shock controls. Asterisks denote significant Tukey post hoc comparisons: \**p* < 0.05, \*\**p* < 0.01, and \*\*\**p* < 0.001.

If neurons encoding graded responses represent the learned contingencies, they should exhibit stability across retrieval sessions. To test this hypothesis, we quantified the proportion of registered neurons that retained their cluster identity across at least two retrieval sessions and compared these values to a shuffled null distribution (10,000 iterations), with multiple comparisons controlled using the Benjamini– Hochberg procedure. Only the positively modulated graded clusters showed significant identity stability across all retrieval intervals (days 1–15, 15–30, and 1–30; Fig. 7a-c). In contrast, negatively modulated graded neurons and a smaller positively modulated cluster in the CS⁺3 group showed stability only between adjacent sessions (Fig. 7b, d, e, Table S5). These data demonstrate that graded clusters remain consistently active, retaining their neuronal profiles, at levels exceeding chance, providing a stable representation of learned contingencies and generalization gradients.

### Activity changes in frequency selective neurons encode learned value

Graded clusters encode learned contingencies but represent only a subset of the active neuronal population. Nevertheless, population-level representations, which incorporate all active neurons, robustly preserve these gradients. This led us to hypothesize that dynamically recruited, frequency-selective neurons also contribute to threat-value representations through changes in response magnitude. Consistent with hippocampal studies showing that spatially selective neurons encode task contingencies via firing-rate modulation (Gagliardi et al., 2024; Huxter et al., 2003; Sanders et al., 2019), we tested whether dynamic, frequency-selective clusters exhibited firing-rate changes proportional to learned significance of the stimuli. To do so, we quantified the normalized baseline-to-stimulus response ratio (BSR) of positively responding neurons within each cluster using CASCADE-inferred spikes (Rupprecht et al., 2021).

We found that although tone-selective clusters emerged independently of learning (i.e., they were present in both conditioned and control mice), their firing rates were shaped by associative learning. Specifically, 3-kHz-selective neurons exhibited higher BSR in both conditioned groups than in controls (p < 0.05), regardless of whether 3 kHz served as the CS⁺ or CS⁻. In contrast, 11- and 15-kHz-selective neurons displayed significantly higher BSR in the CS⁺15 group than in the CS⁺3 or control groups on days 15 and 30 (p < 0.05; Fig. S11, Table S6). Notably, in the CS⁺15 group, where 11 kHz elicited robust threat generalization, the 11- and 15-kHz-selective clusters exhibited similarly high BSRs to both tones. The higher activity rates observed for tones associated with threat or inducing generalization suggests that these neurons are shaped by the animal’s associative experience, including both conditioning and retrieval-dependent updating.

As expected, positively modulated graded neurons closely mirrored population responses, exhibiting the highest BSRs to the CS⁺ and strongly generalized tones and lower BSRs to discriminated tones (p < 0.05; Fig. 7h–i, Table S6). Together, these findings indicate that firing-rate modulation enables both graded and frequency-selective PL neurons to represent the learned contingencies.

### Stable and dynamic neuronal subpopulations encode learned threat value beyond freezing

Our earlier GLM analysis established that tone identity explains neuronal activity more strongly than freezing in the overall PL population, we next asked whether this relationship differed among the neuronal subpopulations identified by our clustering analysis. We first examined frequency-selective (single-tone) neurons, assigning each neuron’s preferred tone according to its cluster identity. Summary statistics were calculated using animals as the experimental unit (CS+15: N = 7; CS+3: N = 5; controls: N = 6).

Among positively modulated single-tone neurons, freezing-corrected tone β coefficients increased monotonically toward the CS+ in opposite directions for the CS+15 and CS+3 groups, whereas coefficients remained relatively flat in controls (Fig. 8a; effect of groups: p < 0.001; group x frequency interaction: p < 0.002, Table S7a). Thus, although these neurons responded preferentially to a single frequency, their β coefficient magnitude scaled with the learned threat value of that tone rather than its absolute frequency.

**Figure 8.**
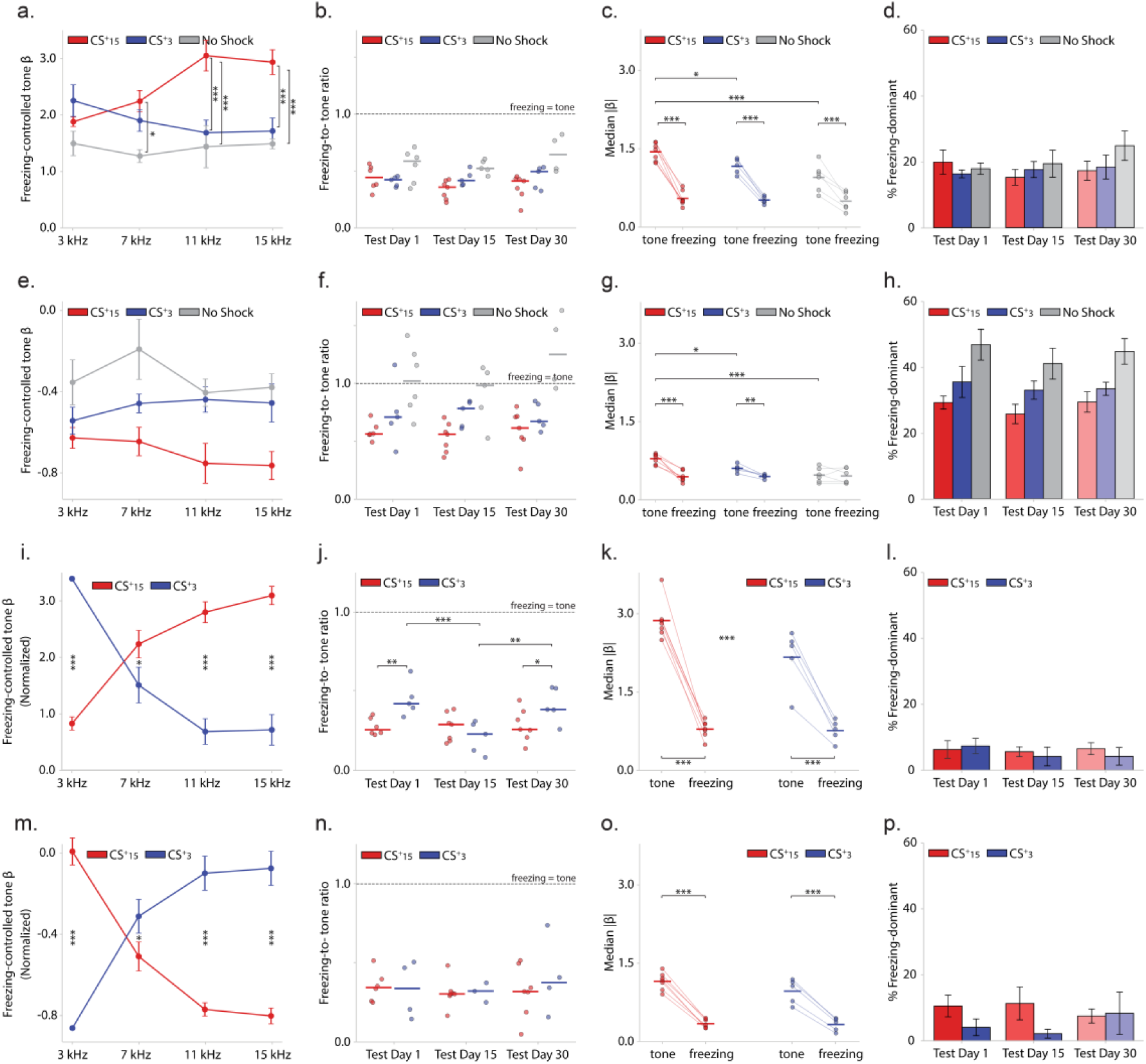
GLM analysis of the contributions of tone identity and freezing to the activity of clustered PL neurons. **a–h.** Positive (**a–d**) and negative (**e–h**) single-tone-selective neurons. **a.** Tone β coefficients in conditioned mice form opposing gradients that peak at the CS+; these gradients are absent in no-shock controls. **b.** Freezing-to-tone ratios remain below 1 across preferred tones and retrieval sessions. **c.** Within-cell comparisons confirm that absolute tone coefficients exceed absolute freezing coefficients in all groups. **d.** The proportion of freezing-dominant neurons is low and stable across retrieval sessions. **e.** Negative tone-selective neurons exhibit suppressive tone β coefficients with weak frequency dependence and greater variability in no-shock controls. **f.** Freezing-to-tone ratios are higher than those of positive responders but remain below 1 in conditioned mice. **g.** Within-cell comparisons show that absolute tone coefficients exceed absolute freezing coefficients in negative responders from conditioned mice, although by a smaller margin than in positive responders. **h.** The proportion of freezing-dominant neurons is higher than among positive responders but remains stable across retrieval sessions. **i–p.** Positive (**i–l**) and negative (**m–p**) graded neurons. **i.** Tone β coefficients obtained after controlling for freezing exhibit strong CS+-centered gradients. **j.** Freezing-to-tone ratios are well below 1, indicating that tone identity contributes substantially more strongly to neuronal activity than freezing. **k.** Within-cell comparisons confirm that absolute tone coefficients greatly exceed absolute freezing coefficients. **l.** Only a small and stable proportion of positive graded neurons is freezing-dominant. **m.** Negative graded neurons exhibit strong suppressive gradients. **n.** Freezing-to-tone ratios remain below 1. **o.** Absolute tone coefficients are approximately threefold greater than absolute freezing coefficients. **p.** The proportion of freezing-dominant neurons remains low across retrieval sessions. Asterisks denote significant Tukey post hoc comparisons: \**p* < 0.05, \*\**p* < 0.01, and \*\*\**p* < 0.001.

Consistent with the population GLM, freezing-to-tone β ratios remained well below 1 (median ≈ 0.4; Fig. 8b), indicating that tone β coefficients were approximately two- to threefold larger than freezing β coefficients. This effect, however, was larger in experimental groups (effect of groups: p < 0.001, Tukey comparisons: CS+15 vs control: p < 0.008, CS+3 vs control: p = 0.08, Table S7b). Likewise, within-cell comparisons showed larger median tone than freezing β coefficients across all groups for each animal (Fig. 8c; effect of group: p < 0.02; effect of state condition (tone vs freezing): p < 0.001; interaction: p < 0.002, Tukey multiple comparisons: Tone vs. freezing p < 0.001 for all groups; β coefficients for tone: CS+15 vs controls: p < 0.00, CS+15 vs CS+3: p < 0.03, Table S7c), and only a small, stable fraction of neurons was classified as freezing-dominant throughout retrieval, with no significant differences between groups across test days (p > 0.005; Table S7d; Fig. 8d). Together, these findings indicate that positively modulated frequency-selective neurons primarily encode learned threat value despite their narrow frequency selectivity.

Negatively modulated single-tone neurons exhibited suppression at their preferred frequency but little evidence of systematic scaling with learned threat value (Fig. 8e, effect of groups: p < 0.001, interaction frequency x group: p > 0.05; Tukey multiple comparisons: CS+15 vs control: p < 0.001; Table S7e). Tone identity remained the stronger predictor of neuronal activity across groups, with lower freezing-to-tone β ratios in the conditioned groups than in no-shock controls (Fig. 8f, effect of group: p < 0.001, Tukey multiple comparisons: CS+15 vs control: p < 0.001, CS+3 vs control: p < 0.006; Table S7f). Within-cell comparisons of tone and freezing β coefficients showed a difference in the experimental groups but not the controls (Fig. 8g, effect of groups: p < 0.05, condition (tone vs freezing): p < 0.001, interaction: p < 0.001, Tukey multiple comparisons: tone vs. freezing: CS+15: p < 0.001; CS+3: p < 0.01; control: p=0.977, Table S7g). Finally, the proportion of freezing-dominant neurons was highest in controls, although this difference reached significance only relative to the CS+15 group (Fig. 8h, effect of group: p < 0.005, Tukey tests: CS+15 vs control: p < 0.005, Table S7h). These findings suggest that negatively responsive frequency-selective neurons multiplex stimulus identity and behavioral state to a greater extent than positively responsive neurons.

We next applied the same analysis to neurons classified as graded (Fig. 7). Because graded clusters were absent in controls, this analysis included only the CS+15 and CS+3 groups. For each graded cell we obtained normalized freezing-corrected β coefficients by dividing each tone beta by the largest beta out of the 4. The normalized freezing-corrected β coefficients formed steep opposing monotonic gradients that tracked learned threat value in both conditioning groups (Fig. 8i; effect of groups: p < 0.001, interaction group x frequency: p < 0.001; Tukey multiple comparisons: CS+15 vs CS+3 for 3, 11, and 15 kHz: p < 0.001; CS+15 vs CS+3 for 7 kHz: p < 0.02, Table S7i). All freezing-to-tone β ratios remained well below 1 across retrieval sessions (Fig. 8j), with the ratios for the CS+15 being even lower than for CS+3 on days 2 and 30 (p < 0.05, Table S7j). The within-cell median β coefficients for freezing and tone were different in both experimental groups (p < 0.001; Fig. 8k), with the tone coefficients being higher in the CS+15 group (p <0.003, Table S7k). Only 4–7% of neurons were classified as freezing-dominant across retrieval sessions (Fig. 8l, Table S7l), a lower percentage than that observed in single-frequency neurons. Similar patterns were observed among negatively modulated graded neurons, although the difference between tone and freezing coefficients was smaller (Fig. 8m–p, Table 7Sm-Sp).

These analyses demonstrate that both dynamically recruited frequency-selective neurons and stable graded neurons encode learned threat value beyond freezing behavior. Graded neurons represented threat value with minimal influence of behavioral state, whereas frequency-selective neurons also encoded threat value, although their responses were more strongly modulated by freezing. Together, these complementary coding strategies likely enable PL populations to maintain stable representations of learned threat value despite ongoing changes in ensemble composition and behavioral state.

## Discussion

Our findings indicate that sensory similarity and learned threat value are not competing explanations of threat generalization but complementary components of a single neural code in the prelimbic cortex. Conditioning organized PL population and single-neuron responses into gradients anchored to the CS+, with novel tones represented according to their similarity to the conditioned stimuli. This graded response structure persisted despite substantial reorganization of ensemble membership across retrieval sessions. Tone identity—which carried opposite learned significance in the two reversed conditioning groups—accounted for more variance in neuronal activity than freezing, indicating that the gradients reflected the learned value of each tone rather than the animal’s ongoing behavioral state. Two populations contributed to this stability in different ways: longitudinally stable graded neurons, whose response profiles spanned the full stimulus continuum and changed little across sessions, and narrowly tuned frequency-selective neurons, which were variably recruited across sessions and whose response magnitudes were modified by learning. Together, these findings suggest that a stable graded code in PL is maintained by a persistent subpopulation while remaining open to modification through ongoing ensemble turnover, supporting adaptive generalization and discrimination.

Generalization has traditionally been explained in terms of perceptual similarity (Shepard, 1987): stimuli resembling a conditioned cue recruit overlapping sensory representations and therefore evoke similar behavioral responses (Corches et al., 2019; Grosso et al., 2018). More recent frameworks propose that responses to novel stimuli depend not only on sensory similarity but also on mnemonic representations of learned threat value, allowing behavior to scale with the predicted significance of each stimulus (Verra et al., 2026; Zaman et al., 2023). This mnemonic contribution should be especially important when generalization is assessed long after conditioning because responding to a novel cue requires retrieval of the associations formed during learning. Our design did not manipulate sensory similarity and learned threat value independently. However, the reversed conditioning procedure dissociated them in one respect: depending on whether 3 or 15 kHz served as the CS+, the neural gradients ran in opposite directions along the same frequency continuum, with each anchored to the threat-associated tone. A gradient determined by spectral features alone would have run in the same direction in both groups. Thus, the learned associations and their mnemonic representations must have determined the anchor and direction of the neural gradient. By contrast, the contribution of sensory similarity is inferred rather than directly demonstrated: intermediate tones were represented in an order that followed their spectral distance from the CS+. Because discriminative conditioning tends to narrow generalization gradients (Dunsmoor & LaBar, 2013; Herzog et al., 2021; Jenkins & Harrison, 1960; Lommen et al., 2017), tracking neural representations over 30 days allowed us to examine how PL populations evolved as behavioral discrimination became progressively more precise during repeated retrieval. Whereas stable neuronal populations retained consistent response profiles across retrieval sessions, neurons recruited after conditioning exhibited changes that paralleled the behavioral sharpening. These results suggest that memory, new learning, and sensory similarity interact to shape generalization gradients.

The ensemble turnover observed here is consistent with previous studies demonstrating dynamic reorganization of cortical activity patterns across temporally separated retrieval sessions (DeNardo et al., 2019; Gallego et al., 2020; Kitamura et al., 2017; Tome et al., 2024). Such reorganization has been proposed to provide flexibility by allowing new information to be incorporated into existing cortical representations while preserving stable behavioral performance (Mau et al., 2020; Zaki & Cai, 2024). Several processes could contribute to this turnover, including additional learning during repeated nonreinforced testing, memory updating, and reconsolidation (Lacagnina et al., 2019; Mau et al., 2020; Sangha, 2015; Zaki & Cai, 2024). In our experiments, turnover was already evident at the first retrieval session, before repeated testing could have exerted a cumulative effect, indicating that this reorganization emerges early after learning. Nevertheless, repeated testing provides a plausible explanation for changes during subsequent sessions. Each retrieval session reactivated the original CS+/CS− associations while presenting the conditioned and intermediate tones without reinforcement, thereby creating opportunities for additional discrimination learning. These experiences may have progressively sharpened differentiation between the CS+ and CS-, modifying the retrieved memory representations and changing which neurons participated in representing the refined associations. Although our data do not establish a mechanistic link between neuronal turnover and additional learning, they raise the possibility that dynamic ensemble membership provides the flexibility needed to incorporate new information.

Turnover was not uniform across the population. Graded neurons retained remarkably consistent response profiles across retrieval sessions, showing that a stable component of the population code can coexist with extensive reorganization of the surrounding ensembles. These neurons scaled their response magnitude with the learned threat value of the conditioned stimuli, and their responses to intermediate tones varied along a continuum that followed each tone’s similarity to the CS+. In this respect PL appears to differ from what has been reported in the amygdala, where distinct populations have been described for stimuli of opposing valence (Grosso et al., 2018) and neurons respond preferentially to the CS+ over the CS− (Grundemann et al., 2019). Rather than segregating threat value across separate populations, individual PL neurons can represent it continuously, encoding the generalization gradient within their own response profiles. The coexistence of these stable neurons with dynamic ensembles may allow repeated retrieval to refine a memory while preserving core features of its original representation (Mau et al., 2020; Tome et al., 2024; Zaki & Cai, 2024). Determining whether these stable graded neurons are causally required for memory storage or retrieval—and whether they constitute a persistent cortical memory trace—will require longitudinal imaging combined with selective manipulation of this population.

Because freezing covaries with threat value, it was important to disentangle their respective contributions to neuronal activity. After accounting for freezing, our GLM analysis revealed opposite signs for the tone-identity beta coefficients in the two conditioning groups, as expected if responses reflect the learned significance of each tone. This relationship was most pronounced among graded neurons, in which tone identity accounted for more variance than freezing in approximately 80–85% of cells. Freezing, however, does not exhaust the conditioned defensive state, and we cannot rule out contributions from unmeasured behavioral or physiological correlates of fear, including vigilance, scanning, orienting, startle, and changes in autonomic state. Our findings are therefore consistent with PL neurons representing the learned significance of the stimuli, but they do not establish that this representation is independent of all fear-related behavioral and physiological states.

Frequency-selective neurons were present in both conditioned and control animals, indicating that their tuning reflects pre-existing sensory properties of PL rather than response profiles created by learning (Hockley & Malmierca, 2024; Zikopoulos & Barbas, 2006). What conditioning changed was their gain: these neurons responded progressively more strongly to threat-associated than to safety-associated tones while retaining their underlying frequency selectivity. Their contribution to the population code was therefore dynamic in two distinct senses—their inferred firing rates were shaped by associative experience, and the particular cells active at any given session varied across retrieval. Gain modulation of an existing sensory-tuned population offers a mechanism by which novel stimuli can be evaluated against both their spectral similarity to conditioned cues and the learned significance of those cues, without requiring a new representation to be constructed for each stimulus.

A comparable organization was apparent among neurons that were suppressed by sound. In the auditory cortex, sound-evoked suppressive (“negative”) responses contribute to lateral inhibition, sharpening frequency tuning and improving sensory discrimination (Kato et al., 2017; Wehr & Zador, 2003). Negative responders in PL did not show robust graded responses when averaged across the population, but clustering revealed subsets whose profiles mirrored those of the positively graded clusters across the stimulus continuum. Whether these suppressed populations provide complementary inhibitory contrast that enhances discrimination while preserving graded representations of learned threat value remains an open question that causal experiments will be required to address.

Together, our findings support a model in which sensory similarity and associative learning make complementary contributions to threat generalization in PL. Associative learning anchors the organization and direction of the generalization gradient, while the ordering of novel stimuli along that gradient follows their sensory similarity to the conditioned cues. This code is sustained by a division of labor between subpopulations: stable graded neurons preserve its core structure across time, whereas dynamically recruited, frequency-selective neurons provide a substrate through which new experience can modify it. More broadly, our results provide a framework for understanding how the PL supports adaptive threat generalization and discrimination.

## Methods

### Subjects

Female and male C57BL/6J mice (IMSR_JAX:000664; Jackson Laboratory, Bar Harbor, ME), aged 8–10 weeks, were housed on a 12 h light/dark cycle. All experiments were conducted during the light phase of the cycle. Animal housing and care were consistent with standards set by the Association for Assessment and Accreditation of Laboratory Animal Care (AAALAC). All experimental procedures were approved by the University of Iowa Institutional Animal Care and Use Committee.

### Discriminatory fear conditioning task

The training day, mice were placed in a square conditioning chamber (20 × 20 cm; Context A, Fig. 1a). The first 3 minutes served as a habituation period. Following habituation, experimental mice received presentations of a conditioned stimulus (CS^+^) tone (15 or 3 kHz, 80 dB, 20.5 s) that co-terminated with a mild foot shock (unconditioned stimulus, US; 0.5 s, 0.5 mA). Between CS^+^ presentations, animals were presented with a non-conditioned stimulus (CS^−^) tone (3 or 15 kHz, 80 dB, 20.5 s) that was never paired with shock. Mice received 10 CS+ tone–shock pairings and 10 CS− tone presentations. Intertrial intervals between CS^+^ and CS^−^ presentations alternated pseudo-randomly between 2, 4, and 6 minutes. Following the final CS^−^ presentation, mice remained in the chamber for an additional 60 s before being returned to their home cages. Control mice were exposed to the same auditory stimuli but did not receive foot shocks. On day 1, mice were placed in a novel context (Context B; 20 × 20 cm; see Fig. 1a) that differed from the training context in wall color, floor texture, and background scent. Animals were allowed to freely explore the chamber for 3 minutes, after which they were presented with auditory stimuli consisting of the conditioned tone (CS^+^) the non-conditioned tone (CS^−^), and two intermediate frequencies (7 and 11 kHz) spanning the same spectral range. These intermediate frequencies were spaced linearly rather than logarithmically. Tones were presented in a semi-random order, with three presentations per frequency and 3-min intertrial intervals. Following the final tone presentation, mice remained in the chamber for an additional 60 s period before being returned to their home cages.

### Behavioral analysis

All behavioral sessions were video recorded using a mounted camera. Freezing behavior was defined as complete immobility except for respiratory movements and was quantified as a percentage of total trial time. Freezing was measured using FreezeFrame 5 software (Actimetrics, RRID:SCR_014429) for non-implanted mice and inertial measurement unit (IMU) data from the Inscopix miniscope system for implanted mice. This approach quantifies movement across three axes and circumvents visual occlusion introduced by the miniendoscope tether. To validate automated scoring, a subset of sessions from implanted and non-implanted mice were randomly selected and manually scored by an experimenter blind to experimental condition, following established criteria (Phillips & LeDoux, 1992; Wang et al., 2012). Automated and manual measurements were highly correlated (Pearson’s r = 0.98 and 0.96), confirming strong inter-method reliability. Freezing responses were analyzed across three retrieval sessions (days 1, 15, and 30). Two animals in the non-shock control group (JAM061 and JAM062) were excluded from repeated-measures analyses due to missing data on day 30 but were retained for day-specific comparisons.

### Viral construct stereotaxic surgery

Mice were anesthetized with isoflurane (3% in oxygen at 1 L/min) and placed in a stereotaxic apparatus (David Kopf Instruments, Tujunga, CA). During surgery, anesthesia was maintained at 1.5%. The head was positioned to ensure a flat skull in both the anterior–posterior (AP) and medial–lateral (ML) planes. Rimadyl (5 mg/kg, s.c.) was administered preoperatively and for two consecutive days postoperatively for analgesia. For calcium imaging experiments, three injections (150 nL each) of AAV1-CaMKIIα-GCaMP6f-WPRE-SV40 (Addgene cat. # 100834) were delivered into the PL at the following stereotaxic coordinates relative to bregma: AP +1.95 mm, ML ±0.3 mm, and DV −1.7, −1.5, and −1.3 mm. Injections were performed at a rate of 50 nL/min. After each injection, the syringe was left in place for 5 minutes to allow for viral diffusion and to minimize backflow. Mice were allowed to recover for four weeks prior to implantation of the GRIN lens and baseplate assembly for in vivo imaging.

### Miniendoscope baseplate placement

Following AAV-GCaMP6f injections, a gradient-index GRIN lens (Inscopix, diameter: 1.0 mm, length: 4.0 mm, numerical aperture: 0.5, model: 1050-004637) and baseplate assembly (Inscopix, Mountain View, CA) were implanted above the PL. Stereotaxic coordinates relative to bregma were AP +1.94 mm, ML ±0.3 mm, and DV −1.5 mm. The cortical surface was identified and set as the zero reference point. To create a tract for lens placement, a 26-gauge syringe needle was lowered to approximately two-thirds of the final depth (DV −1.0 mm) and then slowly retracted. The GRIN lens was subsequently lowered to the final DV coordinate (−1.5 mm) and secured in place with a thin layer of cyanoacrylate adhesive applied around the lens perimeter and over the exposed skull. Dental cement (Lang Dental) was used to further stabilize the baseplate and anchor it to the skull. Mice were allowed to recover for at least two weeks before the onset of behavioral training.

## Calcium imaging analysis

Calcium imaging data acquired using the Inscopix nVoke 2.0 miniscope system were processed using Inscopix Data Processing Software (IDPS; Inscopix, Mountain View, CA). Raw recordings were first preprocessed using the standard IDPS pipeline, which included spatial downsampling, background subtraction, spatial filtering, and rigid motion correction to compensate for movement artifacts. Fluorescence signals were then normalized to ΔF/F, defined as the change in fluorescence relative to a baseline fluorescence level computed for each pixel, to quantify activity-dependent calcium dynamics. Following preprocessing, neuronal signals were extracted using a constrained non-negative matrix factorization approach optimized for one-photon calcium imaging (CNMF-E implemented by IDPS). This method decomposes the imaging data into spatial components corresponding to putative neurons and their associated temporal activity traces while accounting for background and neuropil contamination. Extracted components were manually curated to exclude non-neuronal signals and artifacts based on established morphological and activity criteria (Fig. 1d).

Neuronal activity was aligned to tone onset for all analyses, whereas signal representation, normalization, baseline correction, and temporal binning were adapted to the analytical objective. Sound responder classification, average stimulus-aligned traces, population trace analyses, stimulus response vectors, and the GLM were performed on ΔF/F fluorescence signals, whereas population similarity and baseline-to-stimulus ratio (BSR) analyses used CASCADE-deconvolved activity. Depending on the analysis, normalization consisted of either trial-wise or session-wise z-scoring, with baseline correction subsequently applied when quantifying stimulus-evoked responses. Analysis-specific processing steps are summarized in the table below and details provided in the relevant method subsections.

| **Analysis** | **Input** | **Normalization** | **Temporal processing** |
| --- | --- | --- | --- |
| Sound responder classification | CNMF-e<br>Traces | Trial-wise z-score | Sliding windows |
| Population traces | CNMF-e<br>Traces | Session z-score | Averaged SAT<br>(average-stimulus aligned procedure) |
| Stimulus response vectors | CNMF-e<br>Traces | Session z-score | Concatenated responses |
| GLM | CNMF-e<br>Traces | Session z-score | Frame-by-frame |
| Population similarity | CASCADE | Session z-score | 1-s windows |
| BSR | CASCADE | Session z-score | Session average |

### Sound responder classification

To quantify stimulus-evoked modulation of neuronal activity, we implemented a window-based sound response test (SRT) that detects changes in calcium activity relative to the pre-stimulus baseline while preserving the temporal structure of the response.

#### Data alignment and normalization

For each neuron, ΔF/F traces were aligned to tone onset and resampled onto a common relative time axis spanning a pre-stimulus baseline (10s before tone onset) and the 20-s stimulus period. All time points within that trial were normalized by subtracting the baseline mean and dividing by the standard deviation of that trial’s baseline across time. A small constant was added to the denominator to prevent division by zero. This within-trial normalization places all trials on a common scale prior to being combined and is distinct from the between-trial term described below, which standardizes the reliability of the baseline across trials.

#### Sliding window construction

To capture temporally localized stimulus responses, normalized traces were segmented into overlapping sliding windows spanning the stimulus period. Windows were defined by a fixed duration (1 s) and stride (0.1 s), with window boundaries determined directly from the sampling interval of the data. Only windows fully contained within the stimulus epoch were included. This approach enabled detection of responses with variable onset latencies and durations without assuming a fixed response time.

#### Window-wise response quantification

For each window, the median normalized activity was computed and compared with the corresponding trial baseline median. Baseline-subtracted responses were then averaged across trials to obtain a mean response magnitude for each window. Separately, to down-weight windows in cells whose baseline was unreliable across repetitions, a between-trial standardization was computed for each window as the mean baseline-subtracted response divided by the standard deviation of the per-trial baseline medians across trials. This computation indexes trial-to-trial baseline reliability rather than within-trial fluctuation, and was used only to gate which windows entered the significance test (see below); the response magnitude used to classify sound responders is the trial-averaged, baseline-relative response itself, already expressed in units of each trial’s baseline standard deviation. For single-trial cases, where a between-trial standard deviation is undefined, the trial-averaged baseline-relative response was used directly.

#### Statistical comparison to baseline

For each window, normalized activity values were compared with the pooled baseline distribution across trials using a two-sample Kolmogorov–Smirnov test. This nonparametric test was selected because it detects differences in both central tendency and distribution without assuming normality.

#### Identification of significant response epochs

A window was classified as significant if it met two criteria: (i) a p-value below a predefined threshold (typically p < 0.05) and (ii) a between-trial standardized response of at least one baseline standard deviation. Significant windows were classified as positive or negative according to the sign of the response, and contiguous windows of the same sign were merged into response epochs, allowing gaps of one window or less to account for noise. For each neuron and response direction, the longest response epoch was retained.

#### Summary response metrics

For each response epoch, we extracted the trial-averaged baseline-relative response and associated duration. Based on these metrics, responses were classified as positively modulated (trial-averaged baseline-relative response ≥ 3 and significant for at least 1 s) or negatively modulated (baseline-relative response ≤ −1.5 and significant for at least 1 s), where the response is expressed in units of each trial’s baseline standard deviation.

### Mapping active ensembles across time

For longitudinal analyses, neuronal identity across imaging sessions was tracked using spatial footprint registration to identify putatively identical neurons across days. Spatial footprints of cells detected during conditioning were registered longitudinally using a probabilistic model implemented in CellReg [Inscopix, (Evangelidis & Psarakis, 2008)]. This algorithm aligns neurons based on the similarity of their spatial footprints, enabling consistent tracking of neuronal identity across imaging sessions. To evaluate registration quality, we used the conditioning session as the reference and measured the centroid distances between each neuron and its corresponding match in each retrieval session. For each animal, we generated histograms of these centroid shifts (expressed in micrometers) and compared their distributions with the median equivalent footprint diameter of the neurons. To estimate this value, we thresholded each neuronal footprint at half of its maximum intensity and calculated the diameter of an equivalent circle having the same area. Cells classified as *consistently active* were those reliably detected across all phases of conditioning and retrieval, including sessions on days 1, 15, and 30.

### Average stimulus-aligned trace procedure

For each neuron and tone frequency, **session z-scored** ΔF/F traces were aligned to tone onset, and 40-s epochs (10 s pre-stimulus, 20 s stimulus, 10 s post-stimulus) were extracted. Epochs were linearly interpolated to a common time base (to allow for later averaging), after which the mean activity during the pre-stimulus period was subtracted to baseline-correct each epoch. Baseline-corrected epochs were then averaged across repetitions to generate a single average stimulus-aligned trace (ASAT) for each neuron and tone.

### Generation of Population Sound-Response Curves

Population sound-response curves were generated by first averaging ASATs across neurons within each animal and condition, and then averaging these animal-level traces across animals within each experimental group, representing the population response to each auditory stimulus.

#### Population similarity across repeated tone presentations

CASCADE-inferred firing rates were Gaussian-smoothed (2-s kernel), z-scored within each session, and averaged within overlapping 1-s windows (0.5-s overlap) to generate population activity vectors. Population similarity matrices were computed by correlating these vectors across independent presentations of the same tone (within-tone comparisons) and different tones (cross-tone comparisons). Similarity matrices were not symmetrized, preserving asymmetries arising from stimulus order (e.g., 11 vs. 3 kHz or 3 vs. 11 kHz), tone identity, and stimulus-evoked population dynamics. Matrix axes represent time relative to tone onset (−10 to 30 s), with tone presentation occurring from 0 to 20 s. The earlier tone presentation is plotted on the y-axis and the later presentation on the x-axis. To quantify similarity during stimulus onset, correlation values from the first 5 s of tone presentation were averaged for each animal and compared across tone pairs relative to the CS+ in each experimental group.al, focusing on tone pairs relative to the CS+ in each experimental group.

### Classification of subpopulation tone-response patterns

#### Stimulus response vectors

For each neuron, baseline-corrected ΔF/F responses were extracted from 40-s stimulus-aligned epochs (10 s pre-stimulus, 20 s stimulus, 10 s post-stimulus), interpolated to a common time base, and averaged across repetitions for each tone frequency. Responses to the four frequencies were then concatenated in ascending order to generate a composite response vector for each neuron.

#### Mutual information matrix

For each pair of cells, we computed the mutual information (MI) between their stimulus response vectors to quantify the statistical dependence between their responses. MI measures the reduction in uncertainty about one variable given knowledge of another. In this context, low MI values indicate that the stimulus response vectors of two cells are largely independent, whereas high MI values indicate shared information or coordinated response structure. The MI values for all cell pairs were assembled into a symmetric matrix (n_cells × n_cells), with each entry representing the information shared between a given pair of cells. Separate MI matrices were computed for each experimental group (non-shock, CS+3, CS+15), pooling cells across recording days within each group. In what follows, “MI matrix” refers to one of these matrices.

#### Adjacency matrix construction

To transform the MI matrix into a network graph representation, we applied a summed top-k adjacency procedure. For each cell (matrix row), connections to other cells were ranked by MI strength, and only the top-k values were retained. Each retained connection was weighted by the reciprocal of its rank (1 / rank), favoring the strongest associations. The resulting matrix was symmetrized to represent an undirected network of functional connectivity (if cell A is functionally connected to cell B then this enforces that B is also functionally connected to A).

#### Spectral clustering

The adjacency matrix was analyzed using spectral clustering to identify functional subpopulations. Spectral clustering converts the adjacency matrix into a graph Laplacian, computes its eigenvalues and eigenvectors, and projects the data into a low-dimensional space in which clusters are more separable. K-means clustering was then applied to the eigenvector representations to delineate groups of highly connected neurons. This approach is well suited for detecting non-linear or irregular structures in neuronal co-activity subpopulations.

#### Hyperparameter selection

Two hyperparameters were determined: the top-k value and the number of clusters. We evaluated a range of top-k values (2 to 100) and, for each, computed the first 30 eigenvalues of the normalized graph Laplacian obtained during spectral clustering. Eigenvalues were averaged across top-k values to yield a single representative spectrum. The optimal cluster number was then determined using the eigengap method, defined as the largest gap between consecutive eigenvalues. The top-k values associated with this solution were averaged to obtain the final k.

#### Positively and negatively modulated subpopulations

Because MI is non-directional, it cannot distinguish between positively and negatively correlated responses. To separate these responses, Spearman’s correlation was computed for all pairs of cells within the cluster, and only the sign of each correlation was retained, resulting in a sign matrix of the same dimensions as the MI matrix. A signed MI matrix was then created by element-wise multiplication of the sign matrix by the MI matrix. The signed MI matrix was then used as input for another round of clustering, as previously described. Briefly, this involved generating a top-k adjacency matrix, computing eigenvalues from the normalized Laplacian, determining the optimal number of clusters using the eigengap method, and assigning cluster labels. Based on this procedure, one cluster was separated into three sub-clusters, while all other clusters were separated into two sub-clusters.

#### Global labels

Since our clustering method begins with a MI matrix, and each group has its own MI matrix and associated subclusters and subcluster labels, we performed a final cluster labeling step that allowed for comparison across groups. The results are such that, if a set of cells in group A have been assigned cluster label L, then the set of cells in group B having similar response patterns will be assigned the same cluster label L, allowing for comparison across groups.

#### Cluster Stability

To assess the stability and reorganization of functional clusters across days, we tracked registered cells that were detected on at least two of the three imaging sessions (day 1, day 15, and day 30). For each pair of days (A, B) and each cluster label L on day A, we quantified label stability (“percent_same”) — the percentage of cells that had label *L* on day A and retained the same label on day B:

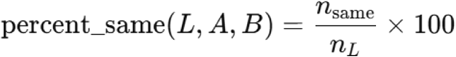

where *n_L_* is the number of cells assigned to label *L* on day A and *n_same_* is the number of those cells that were assigned the same label on day B. Higher percent_same values indicate that a larger fraction of cells maintained the same cluster identity across days, suggesting that the corresponding functional cell assemblies were stable over time. Conversely, lower percent_same values reflect cluster reorganization, where individual cells changed their cluster membership between sessions.

### Baseline-to-stimulus firing rate ratio (BSR)

For each neuron, the mean CASCADE-inferred firing rate during the 20-s stimulus period was divided by the mean firing rate during the 10-s pre-stimulus baseline. Ratios were then averaged across cells within each animal, cell type, and recording day.

### Generalized linear model (GLM)

#### Freezing estimate

Movement was quantified using the inertial measurement unit (IMU) integrated into the head-mounted miniscope. Because the IMU provides band-limited body acceleration rather than locomotion speed, we fitted a two-component Gaussian mixture model to the distribution of log-transformed body-acceleration magnitude. The posterior probability that each imaging frame belonged to the low-movement (freezing) component, *p*_freeze(*t*) ∈ [0,1], was used as a continuous freezing regressor. Samples containing invalid or missing IMU values were identified using a validity mask and excluded from the analysis rather than being classified as freezing. We used a continuous estimate of freezing instead of applying a binary threshold because it captures graded variations in immobility and uncertainty in behavioral state. Consequently, the GLM removes more behavior-related variance from the neural signal than would be expected with a binary freezing classifier. This provides a conservative test of tone encoding: if neuronal activity were primarily explained by freezing, the continuous regressor would account for more of that variance, making it more difficult—not easier—to detect an independent contribution of tone identity.

#### Freezing-controlled time-series GLM

We fitted a generalized linear model (GLM) to the z-scored ΔF/F time series of each neuron to estimate the independent contributions of tone identity and freezing behavior. The design matrix included four tone regressors (3, 7, 11, and 15 kHz), a continuous freezing regressor derived from IMU measurements (see previous subsection), and a trial-number regressor to account for within-session drift. The model was fitted without an intercept so that each tone coefficient represented activity relative to the inter-tone baseline. Standard errors were estimated using heteroskedasticity- and autocorrelation-consistent (HAC; Newey–West (Newly, 1987)) estimators because calcium activity exhibits substantial temporal autocorrelation, making conventional ordinary least squares standard errors anti-conservative. Autocorrelation was estimated within, but not across, trials by resetting the Bartlett kernel at trial boundaries, and the truncation lag was selected using the Newey–West plug-in rule, capped at the duration of a single trial. Statistical significance was assessed using two-sided *t*-tests based on the HAC standard errors.

To ensure that the effects of tone identity and freezing could be estimated independently, we assessed collinearity using the variance inflation factor (VIF). The freezing regressor exhibited minimal collinearity with the tone regressors (median VIF ≈ 1.1), confirming that the independent contributions of tone identity and freezing could be reliably estimated within the same model.

#### Derived per-cell metrics

From each fitted model we extracted the preferred-tone coefficient (β_pref), the freezing coefficient (β_freeze), the freezing-to-tone ratio (|β_freeze|/|β_pref|), and a binary freezing-dominant classification (|β_freeze|>|β_pref|, HAC p<0.05). A neuron was classified as freezing-dominant when the freezing coefficient was significant (HAC *p* < 0.05) and its absolute magnitude exceeded that of the corresponding tone coefficient.

For the general population GLM, in which neurons were not assigned to functional clusters, we used the same cells as used for the population curves and their associated beta tone values. i.e. If a cell contributed to a population response curve of a given tone, then we also used the same cell and its associated beta tone. For frequency-selective neurons, the preferred tone corresponded to the neuron’s preferred tone as determined by the mutual-information (MI) clustering analysis. For the graded neurons, each neurons generated four β tone values that were normalized by dividing by the maximum of the four.

### Statistics

Behavioral and neural data were analyzed using one- or two-way analyses of variance (ANOVAs) with repeated measures. For one-way ANOVAs, tone frequency was treated as the repeated factor; for two-way ANOVAs, a between-subjects group factor (e.g., experimental vs. control) was additionally included. To evaluate the effects of frequency over time across all groups, we performed a three-way mixed-effects ANOVA with group (CS+15, CS+3, and control) as the between-subjects factor and frequency and time as within-subjects (repeated-measures) factors. When ANOVAs revealed significant main effects or interactions, post hoc comparisons were performed using Tukey multiple-comparisons test. Data normality was assessed using the Shapiro–Wilk test and homogeneity of variances with the Brown–Forsythe test prior to statistical analysis. When assumptions were violated, Friedman tests were used for repeated-measures analyses, and Kruskal–Wallis tests were used for comparisons without repeated measures. Rank-based tests were followed by Dunn’s post hoc multiple-comparison tests. Statistical significance was evaluated using a two-sided alpha level of 0.05. For analyses involving multiple cell-identity comparisons, p values were adjusted using the Benjamini–Hochberg false discovery rate procedure.

### Histology

At the conclusion of experiments, mice were deeply anesthetized with isoflurane and transcardially perfused with phosphate-buffered saline (PBS), followed by 4% paraformaldehyde (PFA). Extracted brains were post-fixed in 4% PFA for 24 hours and subsequently transferred to a 20% sucrose solution containing sodium azide at 4 °C for an additional 24 hours. Frozen brains were coronally sectioned at 50 µm using a cryostat, and sections were mounted on Superfrost Plus microscope slides (Fisher Scientific). Histological verification was performed to confirm GCaMP expression and accurate lens placement within the PL.

## Acknowledgements

This works has been funded by NSF (NSF/IOS 2303305 to IAM), NIH (R01 MH123260-01 to IAM; RISE GMO60655 to MRL).

## Author Contributions

MEN developed method to identify subpopulations, wrote code for analysis, and contributed to writing the manuscript, PMO collected data, conducted behavioral and in vivo recording experiments, and contributed to writing the manuscript, MRL contributed to experimental design, collected data, conducted behavioral and in vivo recording experiments. MMRA contributed to the statistical analysis. IAM supervised design, experiments, analysis, and writing of the manuscript.

## References

1. Bontempi, B., Jaffard, R., & Destrade, C. (1996). Differential temporal evolution of post-training changes in regional brain glucose metabolism induced by repeated spatial discrimination training in mice: visualization of the memory consolidation process? Eur J Neurosci, 8(11), 2348–2360. 10.1111/j.1460-9568.1996.tb01198.x

2. Burgos-Robles, A., Vidal-Gonzalez, I., & Quirk, G. J. (2009). Sustained conditioned responses in prelimbic prefrontal neurons are correlated with fear expression and extinction failure. J Neurosci, 29(26), 8474–8482. 10.1523/JNEUROSCI.0378-09.2009

3. Casanova, J. P., Pouget, C., Treiber, N., Agarwal, I., Brimble, M. A., & Vetere, G. (2024). Threat-dependent scaling of prelimbic dynamics to enhance fear representation. Neuron, 112(14), 2304–2314 e2306. 10.1016/j.neuron.2024.04.029

4. Chavez, R. S., & Heatherton, T. F. (2015). Representational similarity of social and valence information in the medial pFC. J Cogn Neurosci, 27(1), 73–82. 10.1162/jocn_a_00697

5. Corches, A., Hiroto, A., Bailey, T. W., Speigel, J. H., 3rd, Pastore, J., Mayford, M., & Korzus, E. (2019). Differential fear conditioning generates prefrontal neural ensembles of safety signals. Behav Brain Res, 360, 169–184. 10.1016/j.bbr.2018.11.042

6. Deitch, D., Rubin, A., & Ziv, Y. (2021). Representational drift in the mouse visual cortex. Curr Biol, 31(19), 4327–4339 e4326. 10.1016/j.cub.2021.07.062

7. DeNardo, L. A., Liu, C. D., Allen, W. E., Adams, E. L., Friedmann, D., Fu, L., Guenthner, C. J., Tessier-Lavigne, M., & Luo, L. (2019). Temporal evolution of cortical ensembles promoting remote memory retrieval. Nat Neurosci, 22(3), 460–469. 10.1038/s41593-018-0318-7

8. Do-Monte, F. H., Quinones-Laracuente, K., & Quirk, G. J. (2015). A temporal shift in the circuits mediating retrieval of fear memory. Nature, 519(7544), 460–463. 10.1038/nature14030

9. Dunsmoor, J. E., & LaBar, K. S. (2013). Effects of discrimination training on fear generalization gradients and perceptual classification in humans. Behav Neurosci, 127(3), 350–356. 10.1037/a0031933

10. Evangelidis, G. D., & Psarakis, E. Z. (2008). Parametric image alignment using enhanced correlation coefficient maximization. IEEE Trans Pattern Anal Mach Intell, 30(10), 1858–1865. 10.1109/TPAMI.2008.113

11. Gagliardi, C. M., Normandin, M. E., Keinath, A. T., Julian, J. B., Lopez, M. R., Ramos-Alvarez, M. M., Epstein, R. A., & Muzzio, I. A. (2024). Distinct neural mechanisms for heading retrieval and context recognition in the hippocampus during spatial reorientation. Nat Commun, 15(1), 5968. 10.1038/s41467-024-50112-7

12. Gallego, J. A., Perich, M. G., Chowdhury, R. H., Solla, S. A., & Miller, L. E. (2020). Long-term stability of cortical population dynamics underlying consistent behavior. Nat Neurosci, 23(2), 260–270. 10.1038/s41593-019-0555-4

13. Grosso, A., Santoni, G., Manassero, E., Renna, A., & Sacchetti, B. (2018). A neuronal basis for fear discrimination in the lateral amygdala. Nat Commun, 9(1), 1214. 10.1038/s41467-018-03682-2

14. Grundemann, J., Bitterman, Y., Lu, T., Krabbe, S., Grewe, B. F., Schnitzer, M. J., & Luthi, A. (2019). Amygdala ensembles encode behavioral states. Science, 364(6437). 10.1126/science.aav8736

15. Herzog, K., Andreatta, M., Schneider, K., Schiele, M. A., Domschke, K., Romanos, M., Deckert, J., & Pauli, P. (2021). Reducing Generalization of Conditioned Fear: Beneficial Impact of Fear Relevance and Feedback in Discrimination Training. Front Psychol, 12, 665711. 10.3389/fpsyg.2021.665711

16. Hockley, A., & Malmierca, M. S. (2024). Auditory processing control by the medial prefrontal cortex: A review of the rodent functional organisation. Hear Res, 443, 108954. 10.1016/j.heares.2024.108954

17. Hoshi, A., Hirayama, Y., Saito, F., Ishiguro, T., Suetani, H., & Kitajo, K. (2023). Spatiotemporal consistency of neural responses to repeatedly presented video stimuli accounts for population preferences. Sci Rep, 13(1), 5532. 10.1038/s41598-023-31751-0

18. Huxter, J., Burgess, N., & O’Keefe, J. (2003). Independent rate and temporal coding in hippocampal pyramidal cells. Nature, 425(6960), 828–832. 10.1038/nature02058

19. Iqbal, J., Kim, S., Lawal, S., Shah, A., Punepalle, L., Sanghvi, H., Gallagher, A., Wilson, A., Xu, B., & Shrestha, P. (2026). A prelimbic molecular clock of protein synthesis for memory persistence. bioRxiv. 10.64898/2026.01.02.697403

20. Jenkins, H. M., & Harrison, R. H. (1960). Effect of discrimination training on auditory generalization. J Exp Psychol, 59, 246–253. 10.1037/h0041661

21. Kato, H. K., Asinof, S. K., & Isaacson, J. S. (2017). Network-Level Control of Frequency Tuning in Auditory Cortex. Neuron, 95(2), 412–423 e414. 10.1016/j.neuron.2017.06.019

22. Kitamura, T., Ogawa, S. K., Roy, D. S., Okuyama, T., Morrissey, M. D., Smith, L. M., Redondo, R. L., & Tonegawa, S. (2017). Engrams and circuits crucial for systems consolidation of a memory. Science, 356(6333), 73–78. 10.1126/science.aam6808

23. Kupke, J., & Oliveira, A. M. M. (2025). The molecular and cellular basis of memory engrams: Mechanisms of synaptic and systems consolidation. Neurobiol Learn Mem, 219, 108057. 10.1016/j.nlm.2025.108057

24. Kyriazi, P., Headley, D. B., & Pare, D. (2020). Different Multidimensional Representations across the Amygdalo-Prefrontal Network during an Approach-Avoidance Task. Neuron, 107(4), 717–730 e715. 10.1016/j.neuron.2020.05.039

25. Lacagnina, A. F., Brockway, E. T., Crovetti, C. R., Shue, F., McCarty, M. J., Sattler, K. P., Lim, S. C., Santos, S. L., Denny, C. A., & Drew, M. R. (2019). Distinct hippocampal engrams control extinction and relapse of fear memory. Nat Neurosci, 22(5), 753–761. 10.1038/s41593-019-0361-z

26. Laufer, O., Israeli, D., & Paz, R. (2016). Behavioral and Neural Mechanisms of Overgeneralization in Anxiety. Curr Biol, 26(6), 713–722. 10.1016/j.cub.2016.01.023

27. Levy, I., & Schiller, D. (2021). Neural Computations of Threat. Trends Cogn Sci, 25(2), 151–171. 10.1016/j.tics.2020.11.007

28. Likhtik, E., Stujenske, J. M., Topiwala, M. A., Harris, A. Z., & Gordon, J. A. (2014). Prefrontal entrainment of amygdala activity signals safety in learned fear and innate anxiety. Nat Neurosci, 17(1), 106–113. 10.1038/nn.3582

29. Lissek, S., Bradford, D. E., Alvarez, R. P., Burton, P., Espensen-Sturges, T., Reynolds, R. C., & Grillon, C. (2014). Neural substrates of classically conditioned fear-generalization in humans: a parametric fMRI study. Soc Cogn Affect Neurosci, 9(8), 1134–1142. 10.1093/scan/nst096

30. Lommen, M. J. J., Duta, M., Vanbrabant, K., de Jong, R., Juechems, K., & Ehlers, A. (2017). Training discrimination diminishes maladaptive avoidance of innocuous stimuli in a fear conditioning paradigm. PLoS One, 12(10), e0184485. 10.1371/journal.pone.0184485

31. Lopez, M. R., Wasberg, S. M. H., Gagliardi, C. M., Normandin, M. E., & Muzzio, I. A. (2024). Mystery of the memory engram: History, current knowledge, and unanswered questions. Neurosci Biobehav Rev, 159, 105574. 10.1016/j.neubiorev.2024.105574

32. Mau, W., Hasselmo, M. E., & Cai, D. J. (2020). The brain in motion: How ensemble fluidity drives memory-updating and flexibility. Elife, 9. 10.7554/eLife.63550

33. Newly, W. K., West, K.D. (1987). A simple, positive semi-definite, heteroskedasticity and autocorrelation consistent covariance matrix. Econometrica, 55(3), 703–708.

34. Phillips, R. G., & LeDoux, J. E. (1992). Differential contribution of amygdala and hippocampus to cued and contextual fear conditioning. Behav Neurosci, 106(2), 274–285. 10.1037//0735-7044.106.2.274

35. Quian Quiroga, R., & Panzeri, S. (2009). Extracting information from neuronal populations: information theory and decoding approaches. Nat Rev Neurosci, 10(3), 173–185. 10.1038/nrn2578

36. Rao-Ruiz, P., Visser, E., Mitric, M., Smit, A. B., & van den Oever, M. C. (2021). A Synaptic Framework for the Persistence of Memory Engrams. Front Synaptic Neurosci, 13, 661476. 10.3389/fnsyn.2021.661476

37. Refaeli, R., Kreisel, T., Groysman, M., Adamsky, A., & Goshen, I. (2023). Engram stability and maturation during systems consolidation. Curr Biol, 33(18), 3942–3950 e3943. 10.1016/j.cub.2023.07.042

38. Rosas-Vidal, L. E., Naskar, S., Mayo, L. M., Perini, I., Masroor, R., Altemus, M., Ramos-Medina, L., Zaidi, S. D., Engelbrektsson, H., Jagasia, P., Heilig, M., & Patel, S. (2025). Prefrontal correlates of fear generalization during endocannabinoid depletion. J Clin Invest, 135(11). 10.1172/JCI179881

39. Rupprecht, P., Carta, S., Hoffmann, A., Echizen, M., Blot, A., Kwan, A. C., Dan, Y., Hofer, S. B., Kitamura, K., Helmchen, F., & Friedrich, R. W. (2021). A database and deep learning toolbox for noise-optimized, generalized spike inference from calcium imaging. Nat Neurosci, 24(9), 1324–1337. 10.1038/s41593-021-00895-5

40. Sanders, H., Ji, D., Sasaki, T., Leutgeb, J. K., Wilson, M. A., & Lisman, J. E. (2019). Temporal coding and rate remapping: Representation of nonspatial information in the hippocampus. Hippocampus, 29(2), 111–127. 10.1002/hipo.23020

41. Sangha, S. (2015). Plasticity of Fear and Safety Neurons of the Amygdala in Response to Fear Extinction. Front Behav Neurosci, 9, 354. 10.3389/fnbeh.2015.00354

42. Shepard, R. N. (1987). Toward a universal law of generalization for psychological science. Science, 237(4820), 1317–1323. 10.1126/science.3629243

43. Sierra-Mercado, D., Padilla-Coreano, N., & Quirk, G. J. (2011). Dissociable roles of prelimbic and infralimbic cortices, ventral hippocampus, and basolateral amygdala in the expression and extinction of conditioned fear. Neuropsychopharmacology, 36(2), 529–538. 10.1038/npp.2010.184

44. Sotres-Bayon, F., & Quirk, G. J. (2010). Prefrontal control of fear: more than just extinction. Curr Opin Neurobiol, 20(2), 231–235. 10.1016/j.conb.2010.02.005

45. Stujenske, J. M., O’Neill, P. K., Fernandes-Henriques, C., Nahmoud, I., Goldburg, S. R., Singh, A., Diaz, L., Labkovich, M., Hardin, W., Bolkan, S. S., Reardon, T. R., Spellman, T. J., Salzman, C. D., Gordon, J. A., Liston, C., & Likhtik, E. (2022). Prelimbic cortex drives discrimination of non-aversion via amygdala somatostatin interneurons. Neuron, 110(14), 2258–2267 e2211. 10.1016/j.neuron.2022.03.020

46. Terranova, J. I., Yokose, J., Osanai, H., Ogawa, S. K., & Kitamura, T. (2023). Systems consolidation induces multiple memory engrams for a flexible recall strategy in observational fear memory in male mice. Nat Commun, 14(1), 3976. 10.1038/s41467-023-39718-5

47. Tome, D. F., Zhang, Y., Aida, T., Mosto, O., Lu, Y., Chen, M., Sadeh, S., Roy, D. S., & Clopath, C. (2024). Dynamic and selective engrams emerge with memory consolidation. Nat Neurosci, 27(3), 561–572. 10.1038/s41593-023-01551-w

48. Verra, L., Spitzer, B., Schuck, N. W., & Zika, O. (2026). Increased generalisation in trait anxiety is driven by aversive value transfer. Commun Psychol, 4(1). 10.1038/s44271-026-00415-w

49. Wang, M. E., Wann, E. G., Yuan, R. K., Ramos Alvarez, M. M., Stead, S. M., & Muzzio, I. A. (2012). Long-term stabilization of place cell remapping produced by a fearful experience. J Neurosci, 32(45), 15802–15814. 10.1523/JNEUROSCI.0480-12.2012

50. Wehr, M., & Zador, A. M. (2003). Balanced inhibition underlies tuning and sharpens spike timing in auditory cortex. Nature, 426(6965), 442–446. 10.1038/nature02116

51. Zaki, Y., & Cai, D. J. (2024). Memory engram stability and flexibility. Neuropsychopharmacology, 50(1), 285–293. 10.1038/s41386-024-01979-z

52. Zaman, J., Yu, K., & Verheyen, S. (2023). The idiosyncratic nature of how individuals perceive, represent, and remember their surroundings and its impact on learning-based generalization. J Exp Psychol Gen, 152(8), 2345–2358. 10.1037/xge0001403

53. Zikopoulos, B., & Barbas, H. (2006). Prefrontal projections to the thalamic reticular nucleus form a unique circuit for attentional mechanisms. J Neurosci, 26(28), 7348–7361. 10.1523/JNEUROSCI.5511-05.2006

