## Supplemental Figures S1-S11 for "Complementary stable and dynamic prelimbic ensembles encode learned threat value underlying generalization and discrimination"

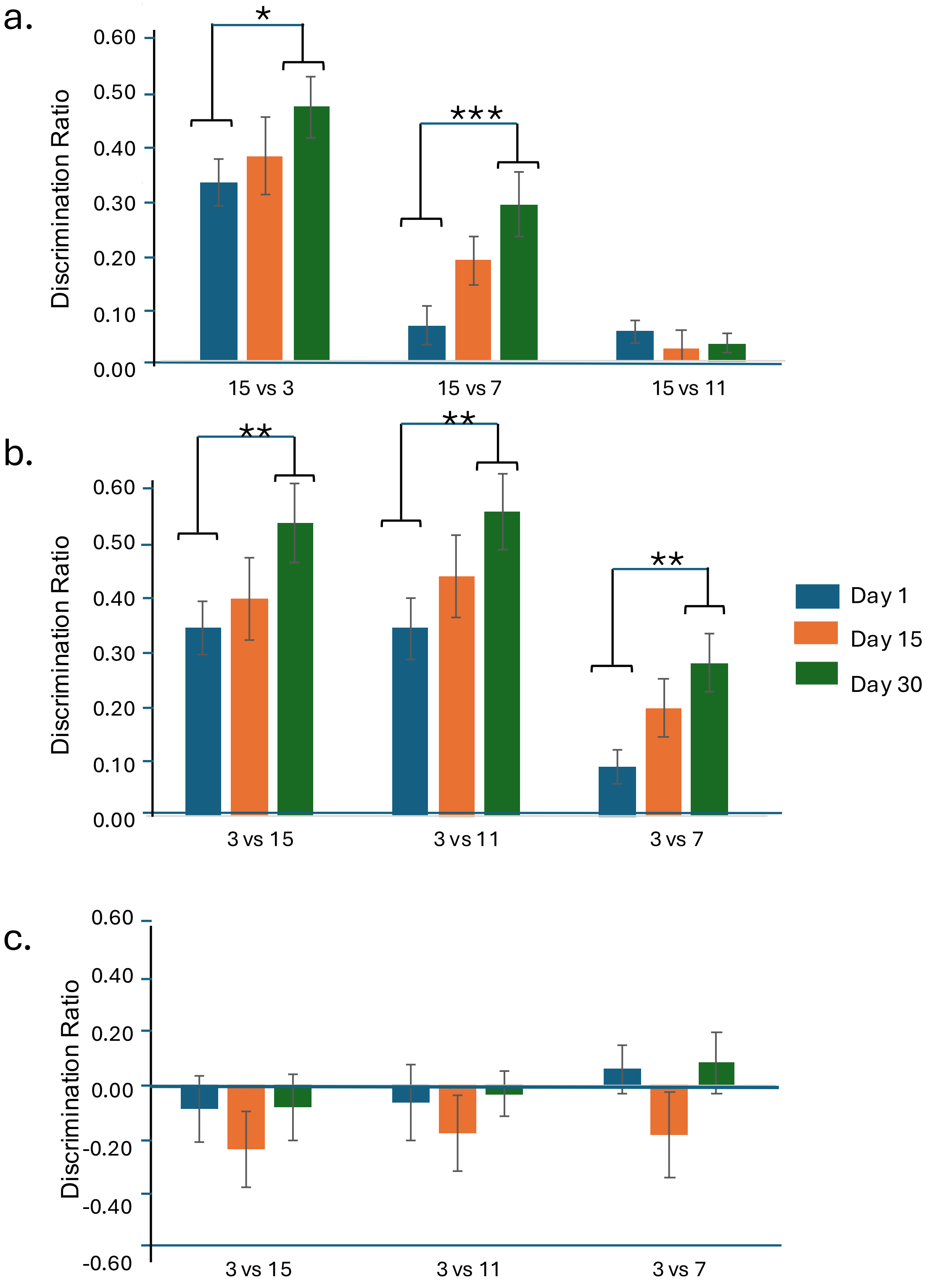


**Figure S1. (a–c)** Discrimination ratios (DRs) were used to quantify discrimination between each frequency and the CS+ across testing days in the CS+15 group (a), CS+3 group (b), and control animals (c). In both experimental groups, discrimination increased over time, particularly between the CS+ and the CS− and between the CS+ and the frequency adjacent to the CS−. Discrimination between the CS+ and its adjacent frequency also increased across testing days in the CS+3 group but not in the CS+15 group (Tukey's multiple comparisons). In contrast, control animals showed no consistent changes in discrimination over time. Repeated-measures ANOVAs with frequency comparison and testing day as factors revealed significant effects of frequency comparison, testing day, and their interaction in the CS+15 group (frequency: $F(2,52) = 44.362$, $p < 0.001$; day: $F(2,52) = 4.822$, $p = 0.012$; interaction: $F(4,104) = 3.421$, $p = 0.011$). In the CS+3 group, significant effects of frequency comparison and testing day were observed, but not their interaction (frequency: $F(2,42) = 24.415$, $p < 0.001$; day: $F(2,42) = 5.499$, $p = 0.008$; interaction: $F(4,84) = 0.23$, $p = 0.921$). No significant effects were detected in control animals (frequency: $F(2,20) = 1.21$, $p = 0.319$; day: $F(2,20) = 0.767$, $p = 0.477$; interaction: $F(4,40) = 0.154$, $p = 0.960$). Asterisks denote Tukey's multiple comparisons: $p < 0.05$; ** $p < 0.01$; *** $p < 0.001$.

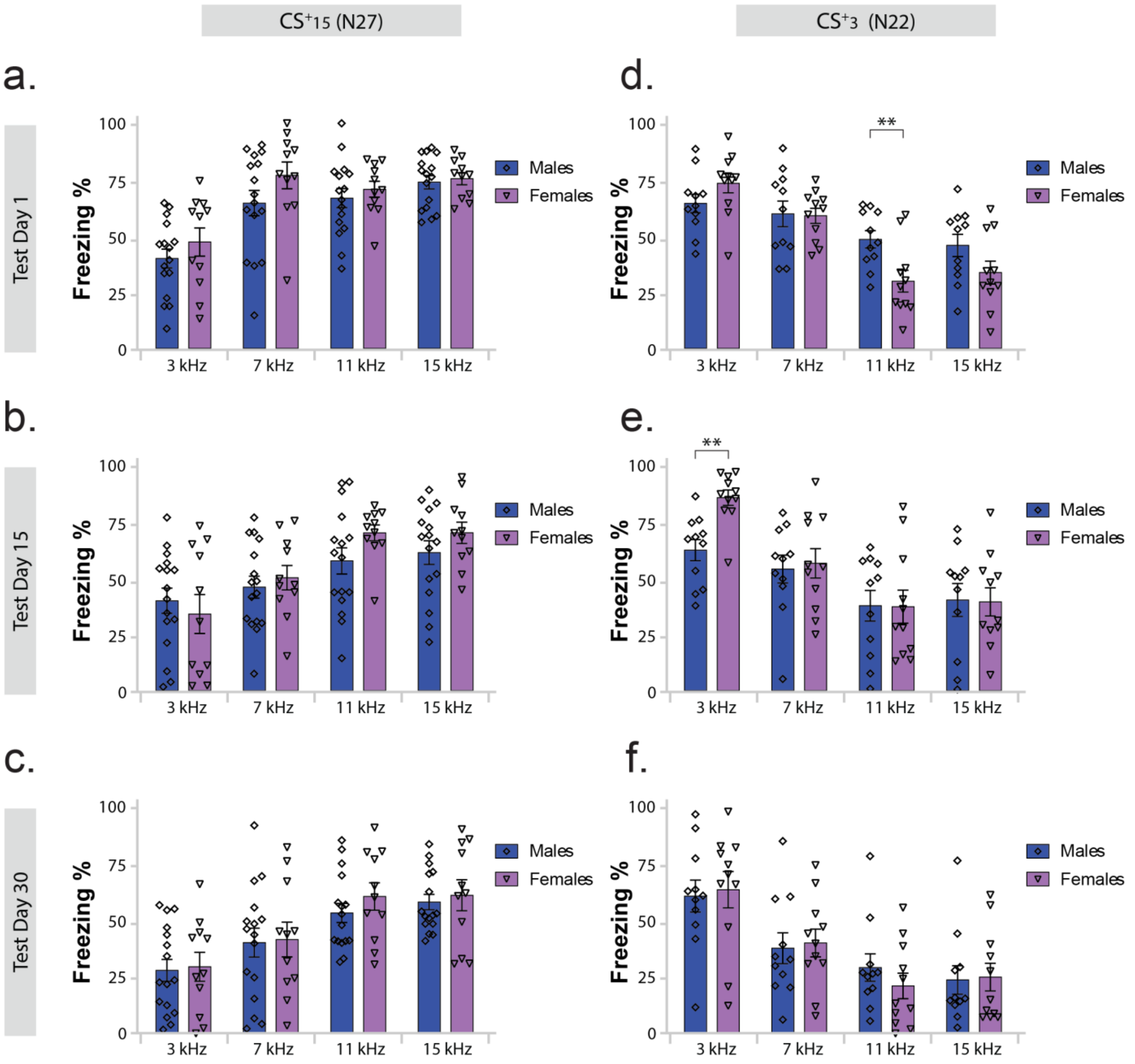


**Figure S2. Sex differences in behavior.**

**(a-c) Sex differences in CS$^+$15-trained mice.** No sex differences were observed on any testing day (effect of sex: day 1, F(1,25) = 1.52, *p* = 0.23; day 15, F(1,25) = 0.55, *p* = 0.467; day 30, F(1,25) = 0.17, *p* = 0.681). On all days, there was a significant main effect of frequency, indicating successful learning in both males and females (day 1, F(3,75) = 33.46, *p* < 0.001; day 15, F(3,75) = 26.32, *p* < 0.001; day 30, F(3,75) = 51.47, *p* < 0.001). No sex × frequency interactions were detected on any testing day (*p* > 0.05). Post hoc comparisons showed that on day 1, mice discriminated 3 kHz from 7, 11 and 15 kHz (*p* < 0.05), but generalized among the higher frequencies (*p* > 0.05). On days 15 and 30, mice discriminated 3 kHz from 7, 11 and 15 kHz (*p* < 0.05) and 7 kHz from 11 and 15 kHz (*p* < 0.05), while generalization persisted between 11 and 15 kHz (*p* > 0.05).

**(d-f) Sex differences in CS$^+$3-trained mice.** No main effect of sex was observed on any testing day (day 1, F(1,20) = 1.24, *p* = 0.279; day 15, F(1,20) = 0.78, *p* = 0.388; day 30, F(1,20) = 0.003, *p* = 0.956). A significant main effect of frequency was present across all days, indicating learning in both sexes (day 1, F(3,60) = 40.65, *p* < 0.001; day 15, F(3,60) = 30.17, *p* < 0.001; day 30, F(3,60) = 35.19, *p* < 0.001). Post hoc comparisons indicated discrimination among all frequencies across days (*p* < 0.05), except between 11 and 15 kHz (*p* > 0.05). Significant sex × frequency interactions were observed on day 1 (F(3,60) = 6.91, *p* < 0.001) and day 15 (F(3,60) = 3.55, *p* < 0.02), reflecting sex-specific differences at 11 kHz on day 1 and at 3 kHz on day 15. These interaction effects were not present on day 30 (F(3,60) = 0.76, *p* = 0.522) and likely reflect increased behavioral variability rather than stable sex differences.

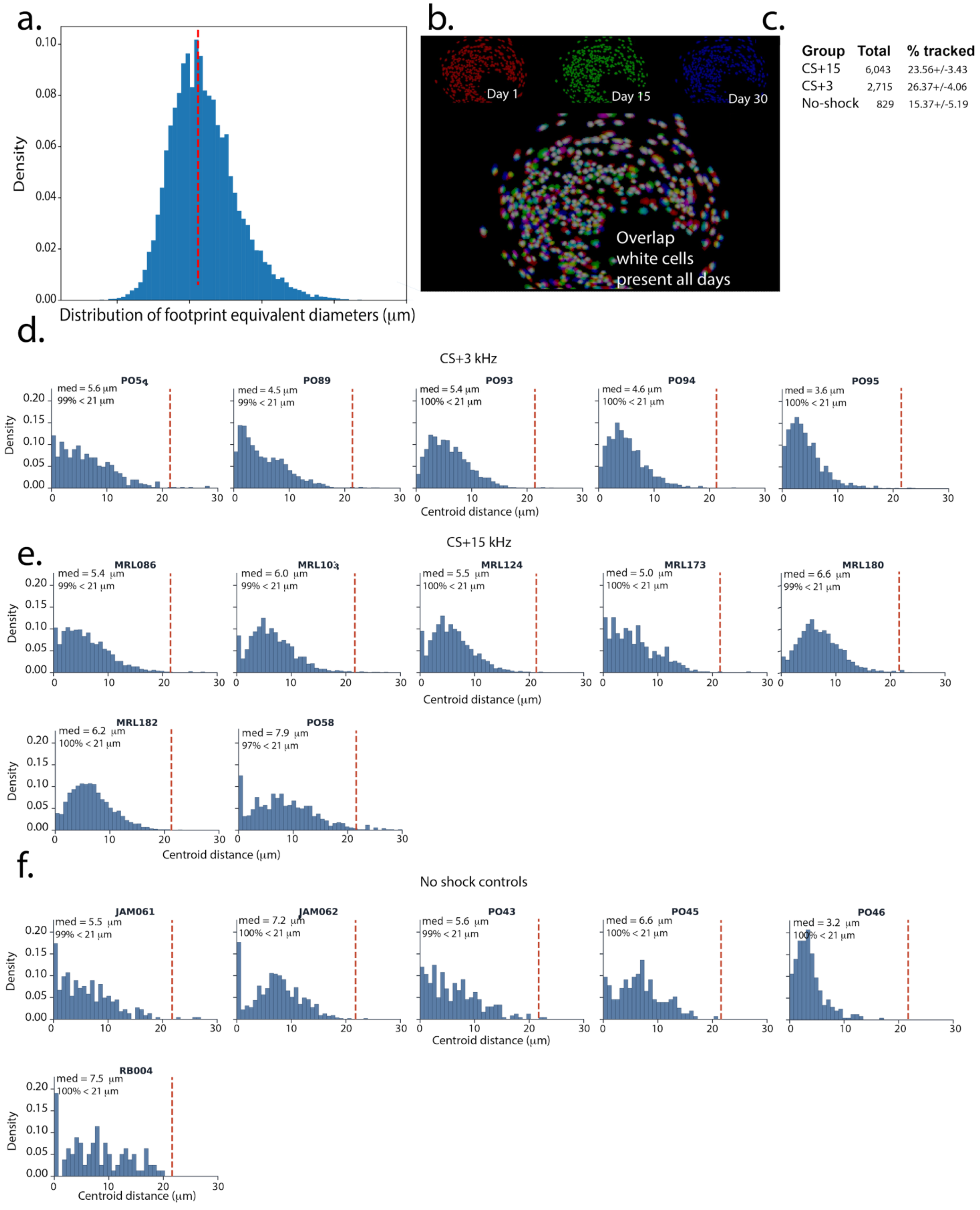


**Figure S3. Evaluation of cell registration quality.**

**(a)** Distribution of equivalent neuronal footprint diameters (µm) for all registered neurons. **(b)** Representative spatial footprints recorded on Day 1 (red), Day 15 (green), and Day 30 (blue), along with the merged image showing neurons tracked throughout the experiment (white). **(c)** Total number of neurons recorded in each experimental group and the percentage of neurons successfully registered across all sessions. Note that two control animals were excluded because they lost the GRIN lens after Day 15. **(d–f)** Distributions of centroid shifts between matched neurons in the retrieval sessions relative to the conditioning session for the CS+3 **(d)**, CS+15 **(e)**, and control **(f)** groups. The red line indicates the median equivalent neuronal footprint diameter (21.4 µm). The upper left corner of each panel shows the median centroid shift and the percentage of registered neurons with centroid shifts smaller than the median footprint diameter.

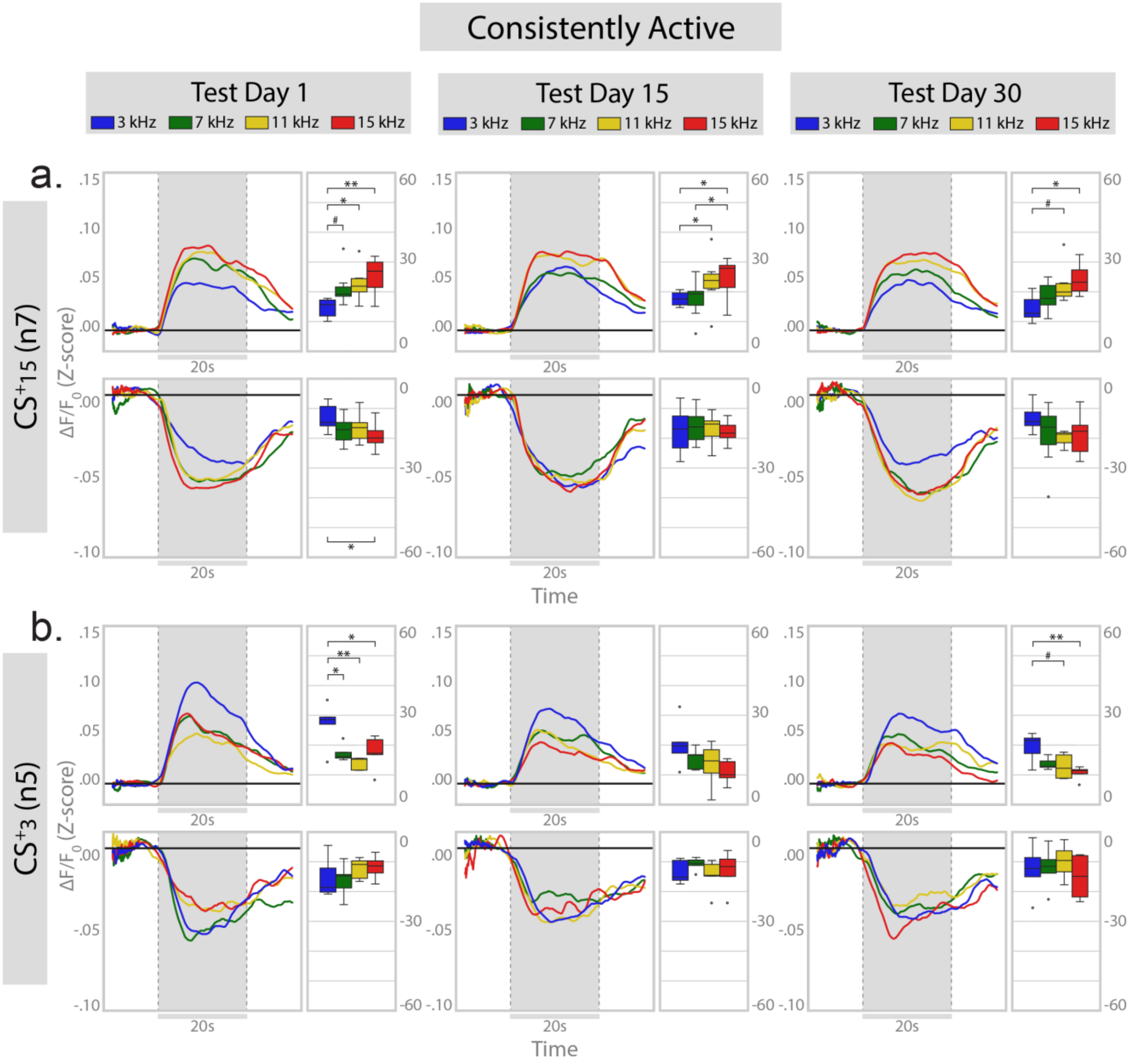


**Figure S4. Population activity of consistently active neurons.**
**(a-b)** Population activity of consistently active neurons from animals trained with a 15 kHz CS+ **(a)** or a 3 kHz CS+ **(b).** Upper panels show positively tone-responsive neurons and lower panels show negatively tone-responsive neurons. Adjacent boxplots display areas under the population response curves (AUC). Boxplots show the median (center line), interquartile range (box), and whiskers extending to ±1.5× the interquartile range; points outside the whiskers represent individual observations beyond this range. Consistently active positive responders exhibit graded responses across test days (day 1 to day 30). *$p < 0.05$; **$p < 0.01$.

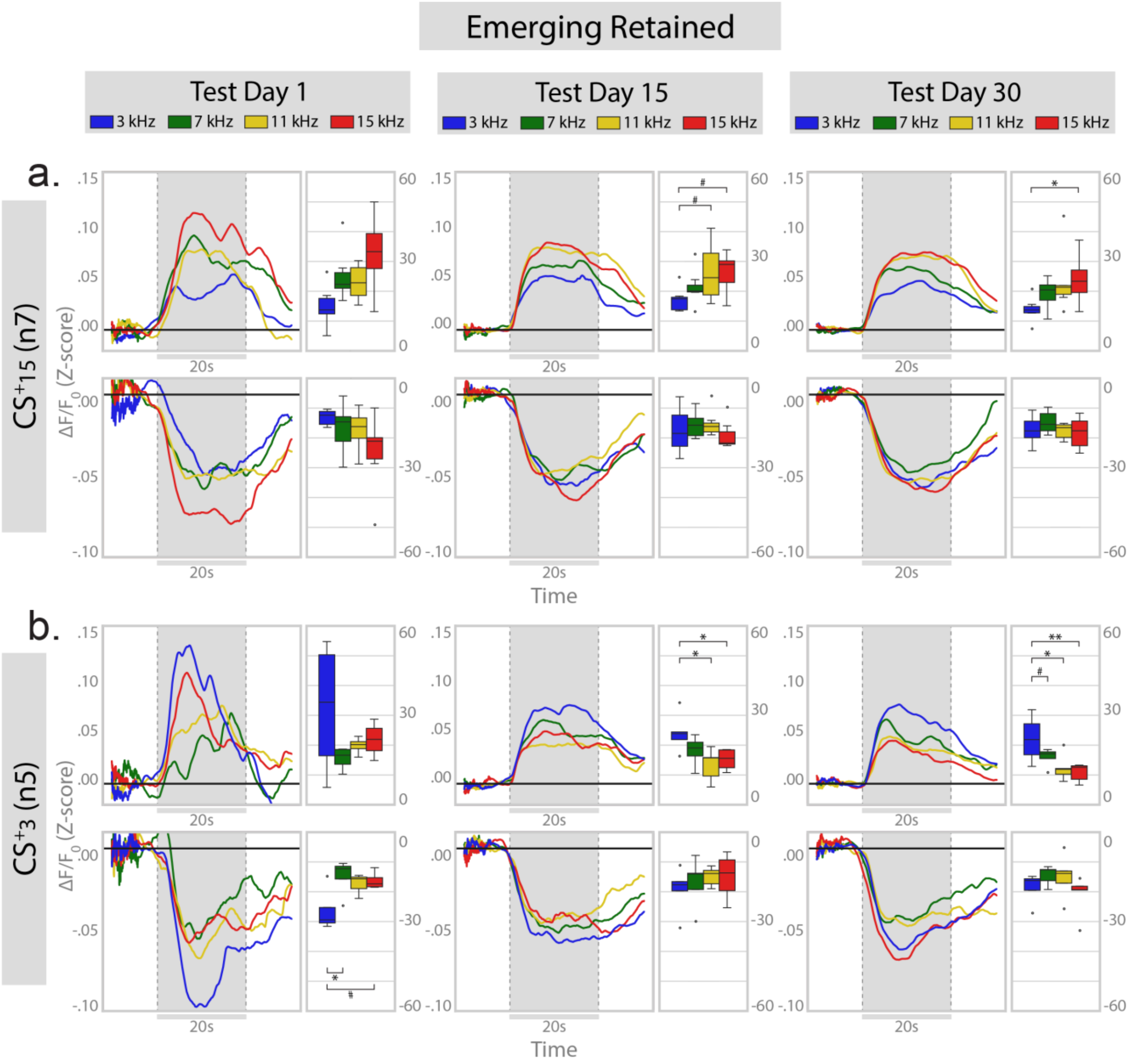


**Figure S5. Population activity of emerging-retained neurons.**
Emerging-retained neurons become part of the active ensemble after conditioning and remaining active after that. **(a-b)** Population activity of emerging-retained neurons from animals trained with a 15 kHz CS+ **(a)** or a 3 kHz CS+ **(b).** Upper panels show positively tone-responsive neurons and lower panels show negatively tone-responsive neurons. Adjacent boxplots display areas under the population response curves. Emerging-retained neurons show greater variability on day 1 compared with later test days. Only positive responders show significant graded valence. $^*p < 0.05$; $^{**}p < 0.01$.

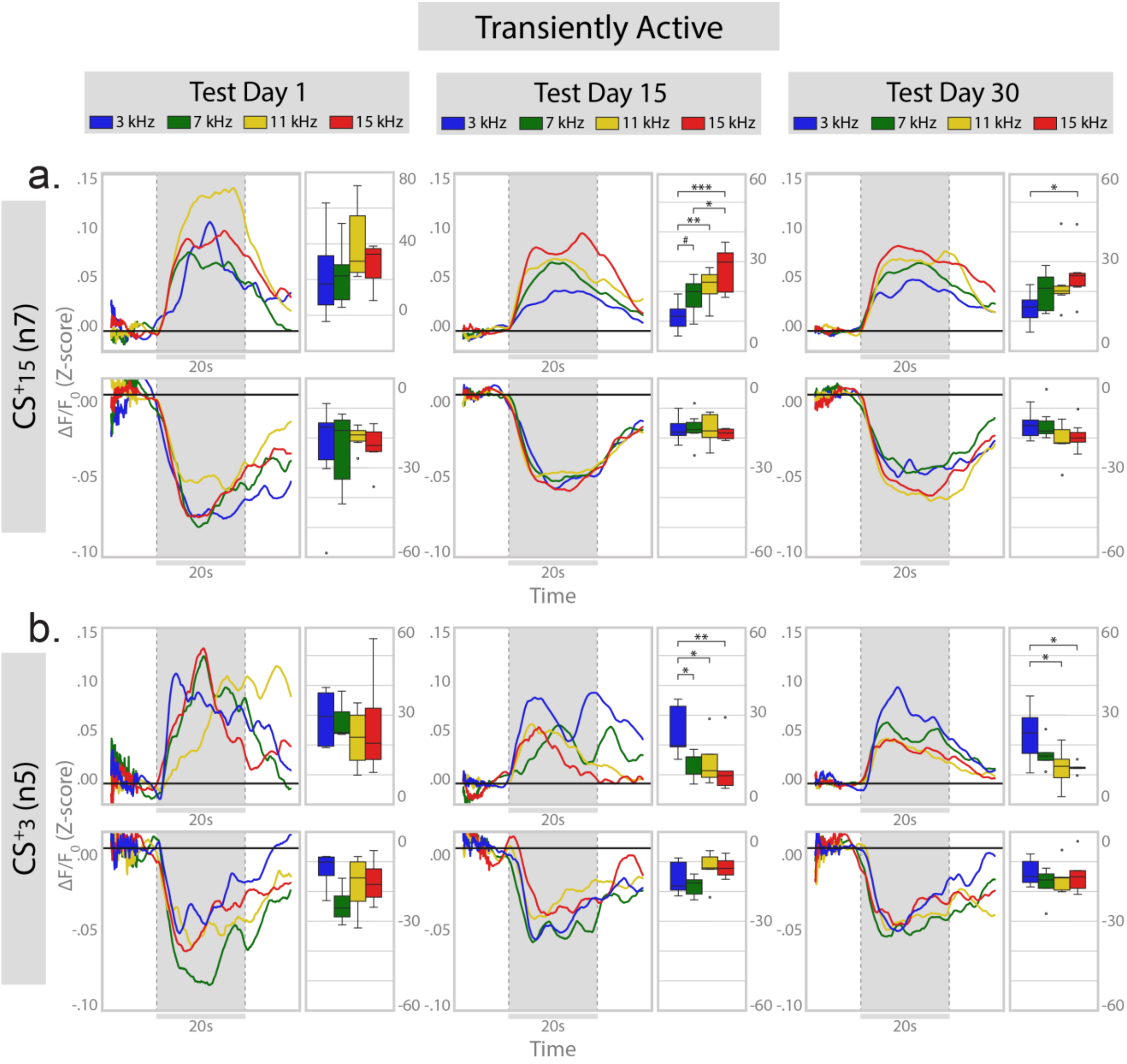


**Figure S6. Population activity of transiently active neurons.**
Transiently active neurons were active on only a single test day.
**(a-b)** Population activity of transiently active neurons from animals trained with a 15 kHz CS+ **(a)** or a 3 kHz CS+ **(b).** Upper panels show positively tone-responsive neurons and lower panels show negatively tone-responsive neurons. Adjacent boxplots display areas under the population response curves. Transiently active neurons did not show graded population responses on day 1; graded responses emerged by day 15 and were maintained through day 30 in positive sound responder neurons. $*p < 0.05$; $**p < 0.01$.

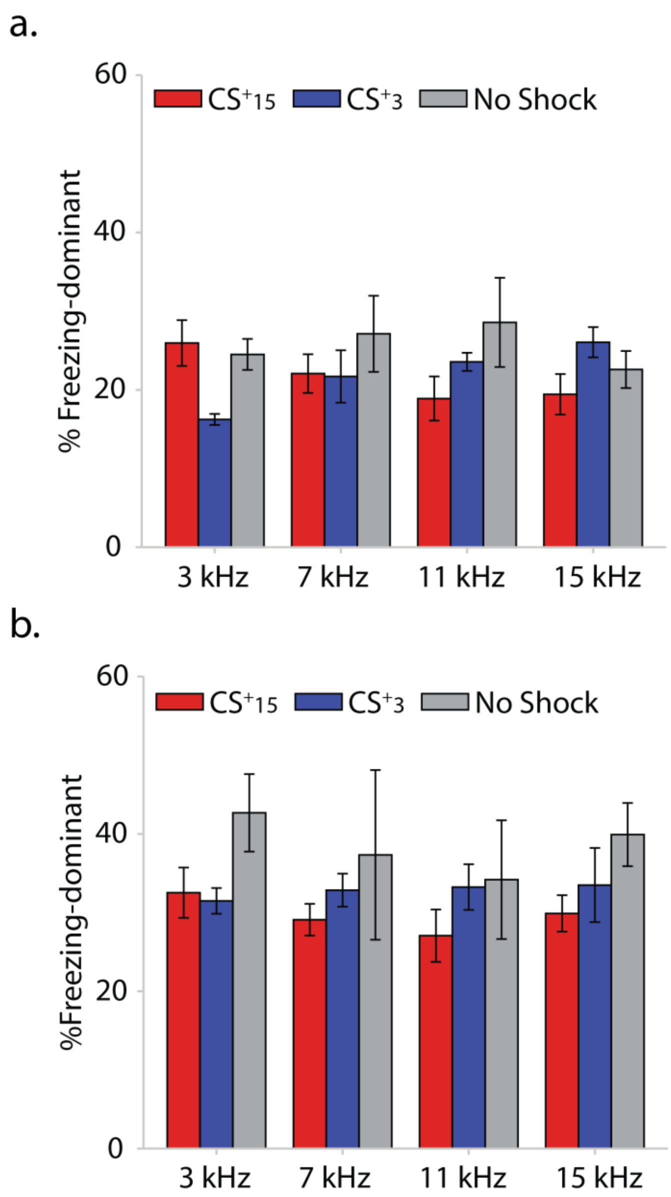


**Figure S7.** (a–b) Percentage of freezing-dominant neurons responding to each tone among positive (a) and negative (b) responders. A repeated-measures ANOVA, with group as the between-subjects factor and frequency as the within-subjects factor, revealed a significant group × frequency interaction [$F(6, 45) = 2.402$, $p < 0.05$]. No main effects of group or frequency were observed ($p > 0.05$). However, Tukey-corrected multiple comparisons revealed no significant pairwise differences, with only trends at 3 kHz between the CS+15 and CS+3 groups ($p = 0.087$) and at 11 kHz between the CS+15 and no-shock groups ($p = 0.069$).

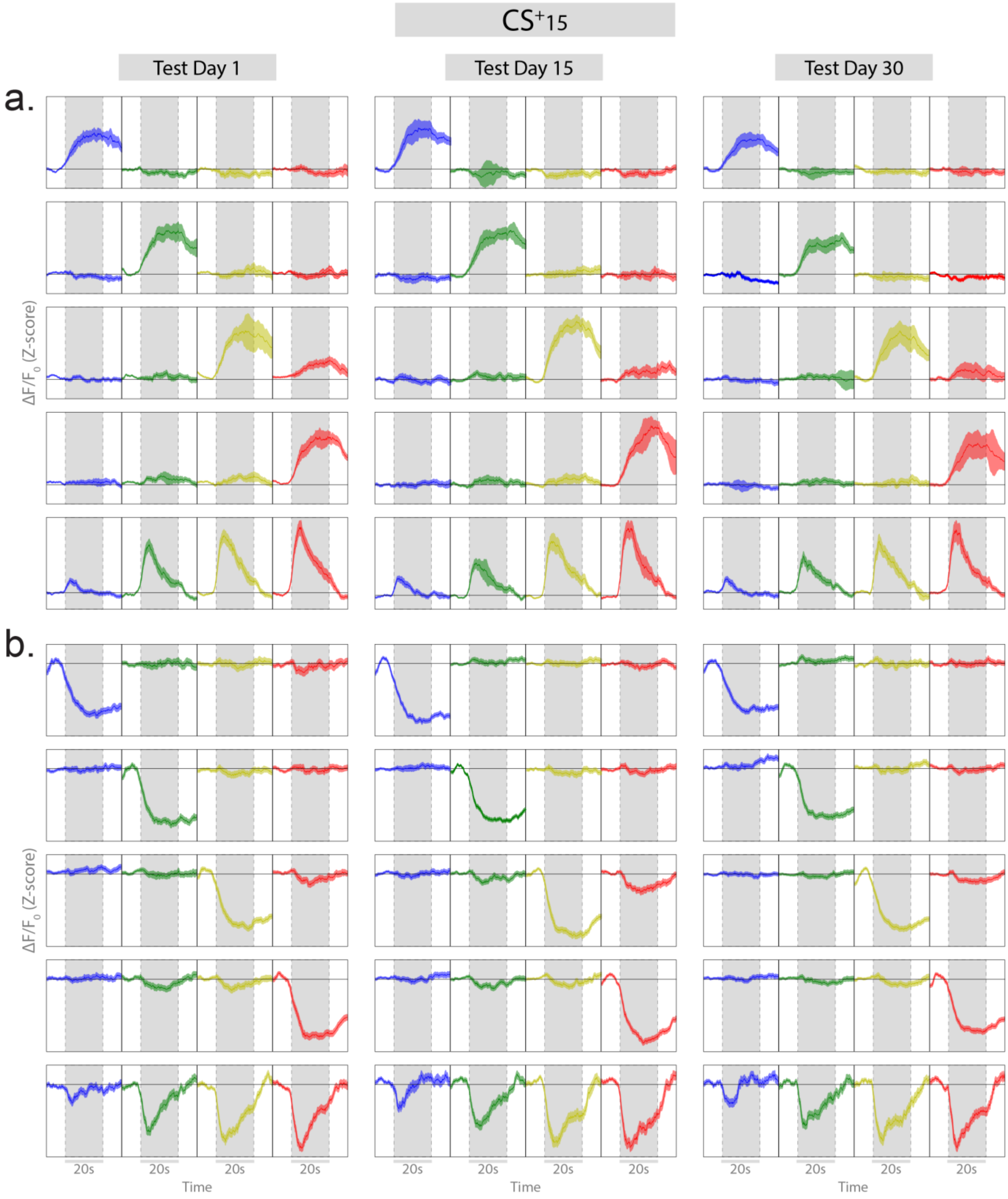


**Figure S8**. **Clustering of PL subnetworks based on signed mutual information for the CS+ 15 kHz group per testing session.**
**(a-b)** Average stimulus-aligned population responses for clusters showing positive **(a)** or negative **(b)** modulation to individual tones (3, 7, 11, or 15 kHz, upper panels) or graded emotional tuning **(a-b, bottom panels).**

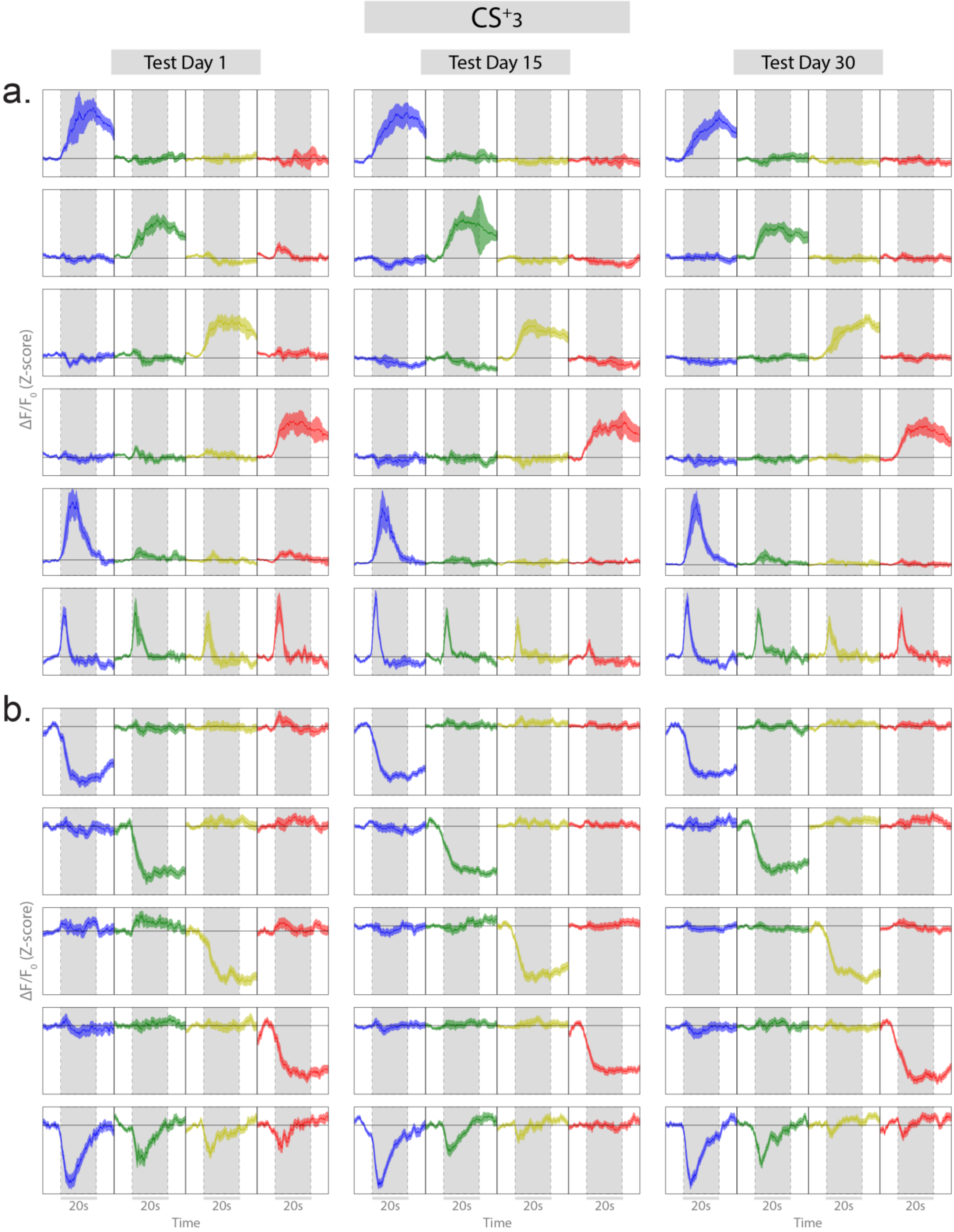


**Figure S9. Clustering of PL subnetworks based on signed mutual information for the CS+ 3 kHz group per testing session.**
**(a-b)** Average stimulus-aligned population responses for clusters showing positive **(a)** or negative **(b)** modulation to individual tones (3, 7, 11, or 15 kHz, upper panels) or graded emotional tuning **(a-b, bottom panels).**

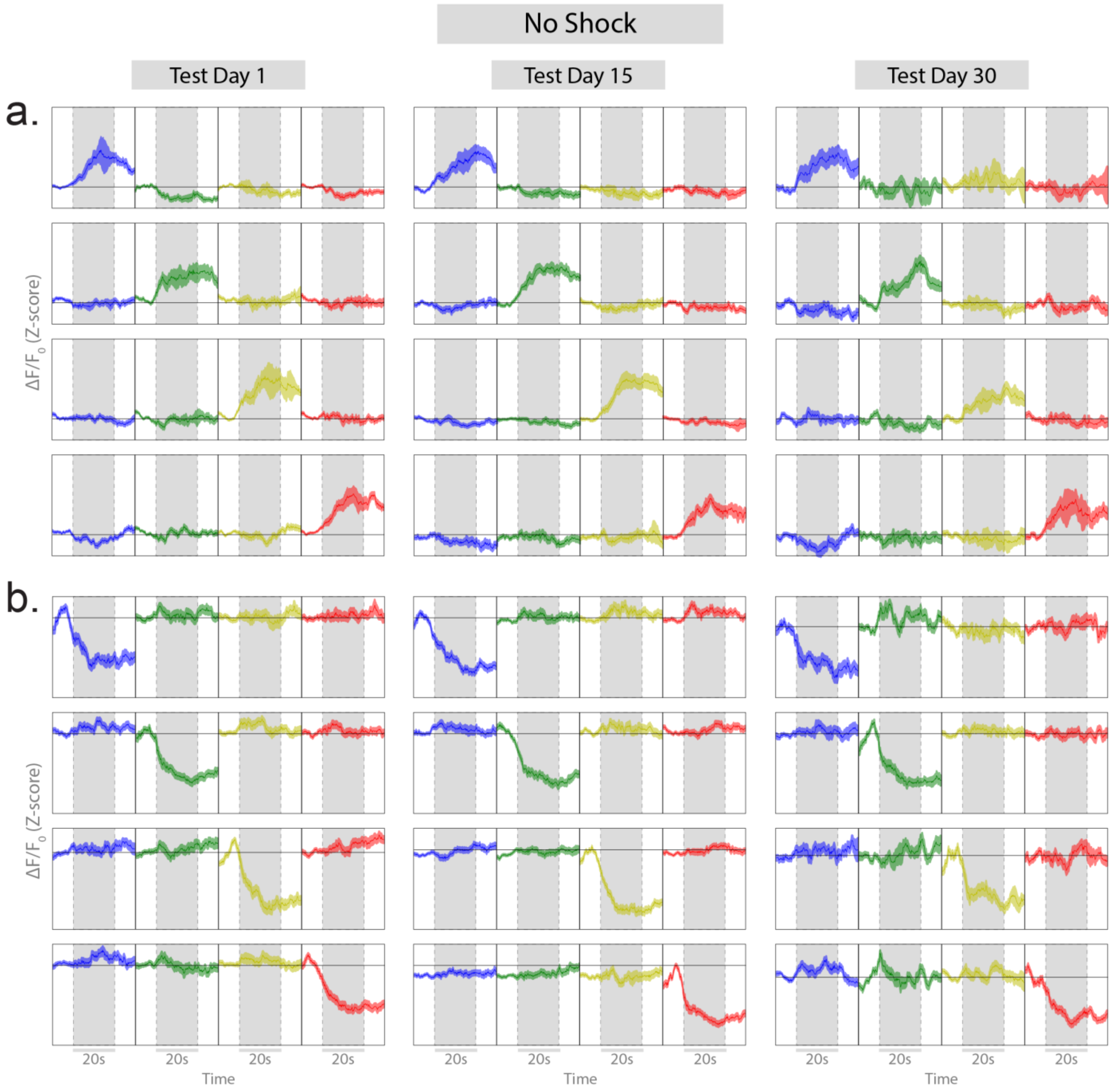


**Figure S10. Clustering of PL subnetworks based on signed mutual information for the no shock control per testing session.**
**(a-b)** Average stimulus-aligned population responses for clusters showing positive **(a)** or negative **(b)** modulation to individual tones (3, 7, 11, or 15 kHz).

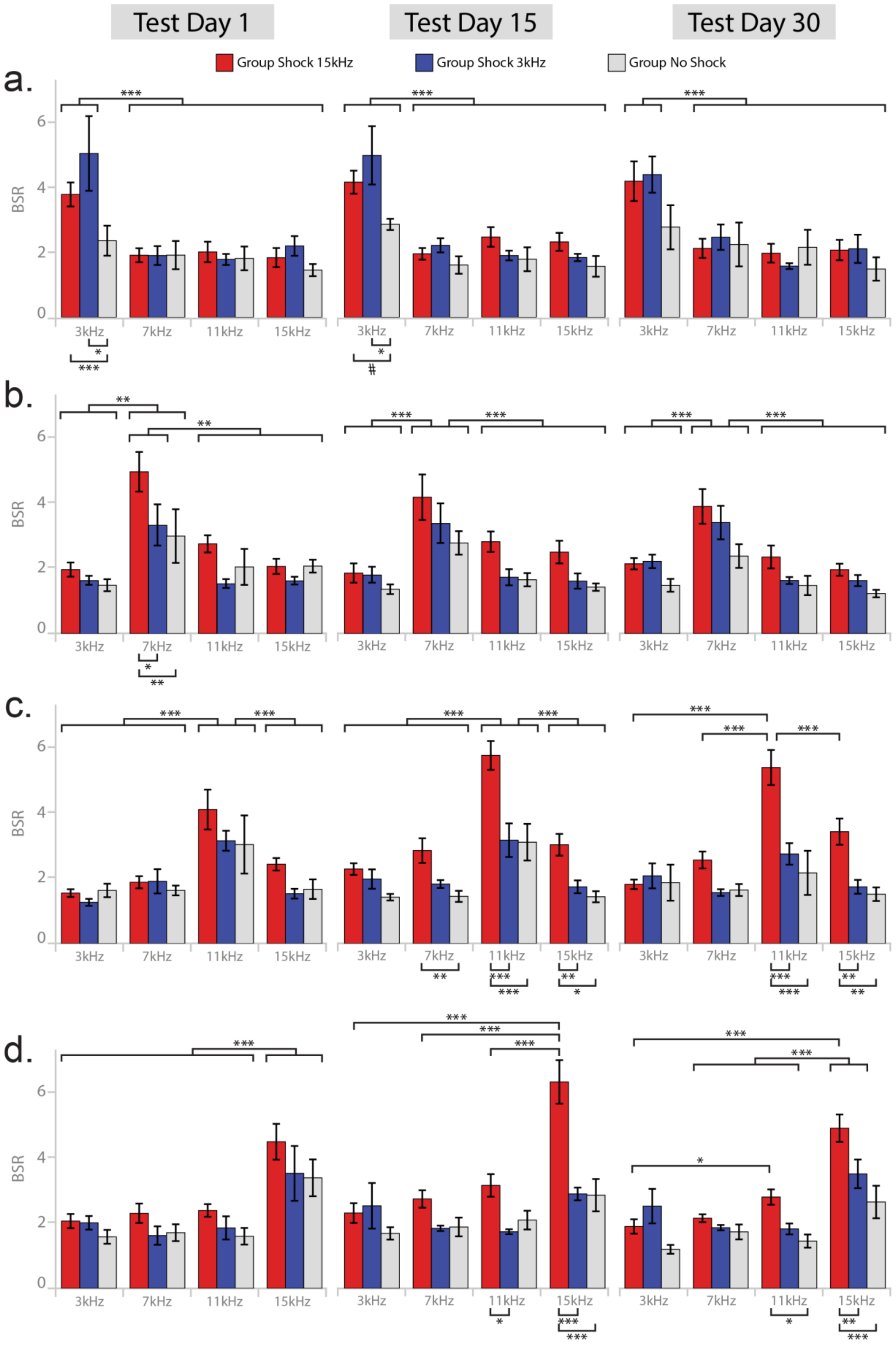


**Figure S11. a–d,** Baseline-to-stimulus firing rate ratio (BSR) illustrating changes in positively responding neurons within tone-specific clusters shown in Fig. 5, for the CS+15 (red), CS+3 (blue), and control groups.

**(a)** BSR of neurons in clusters primarily responsive to 3 kHz (Fig. 5c.1), **(b)** 7 kHz (Fig. 5c.2), **(c)** 11 kHz (Fig. 5c.3), and **(d)** 15 kHz (Fig. 5c.4). ANOVA results are reported in Table S3; Tukey's multiple-comparison tests indicate significance (*p* < 0.05; ***p*** < 0.01; *p* < 0.001).
