## Supplementary material for "Complementary stable and dynamic prelimbic ensembles encode learned threat value underlying generalization and discrimination": Tables S1-S7

**Table S1.**

**Fig. 2.**

| Analysis | Cell class/session | Variable | Sum Sq. | Mean Sq. | F | df | p | ηp² |
|---|---|---|---|---|---|---|---|---|
| One-way ANOVA | Consistently active neurons | Group | 0.0261 | 0.0130 | 2.704 | 2, 13 | 0.104 | 0.29 |
| One-way ANOVA | Emerged-retained neurons: Day 1 and 15 | Group | 0.0065 | 0.00325 | 1.445 | 2, 13 | 0.271 | 0.18 |
| One-way ANOVA | Emerged-retained neurons: Day 15 and 30 | Group | 0.00697 | 0.00349 | 1.018 | 2, 13 | 0.388 | 0.14 |
| One-way ANOVA | Emerged-retained neurons: Day 1 and 30 | Group | 0.0084 | 0.0042 | 0.332 | 2, 13 | 0.723 | 0.05 |
| One-way ANOVA | Transiently active neurons: Day 1 | Group | 0.000194 | 0.000097 | 0.0598 | 2, 13 | 0.942 | 0.01 |
| Kruskal-Wallis | Transiently active neurons: Day 15 | Group | NA | NA | 1.901 | 2 | 0.387 | $\varepsilon^2$=0.00 |
| Kruskal-Wallis | Transiently active neurons: Day 30 | Group | NA | NA | 7.515 | 2 | 0.023* | $\varepsilon^2$=0.42 |

**Pairwise multiple comparisons (Dunn's method)**

| Comparison | Diff. of ranks | Q | p |
|---|---|---|---|
| No shock vs. CS+15 | 8.179 | 2.741 | 0.018* |
| No shock vs. CS+3 | 5.350 | 1.675 | 0.282 |
| CS+15 vs. CS+3 | 2.829 | 1.015 | 0.931 |

**Table S1.** Statistical analyses corresponding to the Venn diagrams shown in Fig. 2. One-way ANOVAs were used when the assumptions of normality and homogeneity of variance were met; otherwise, Kruskal–Wallis tests were performed. Significant Kruskal–Wallis tests were followed by Dunn's multiple-comparisons tests. Effect sizes are reported as partial eta squared ($\eta p^2$) for ANOVAs and epsilon squared ($\varepsilon^2$) for Kruskal–Wallis tests.

**Table S2.**

### Fig. 3. 3-way mixed model ANOVA

#### a. Positive modulated

| Modulation | Variable | Sum Sq. | Mean Sq. | F | df | p | ηp² |
|---|---|---|---|---|---|---|---|
| Positive | Group | 41614 | 20807.1 | 16.803 | 2, 17.907 | < 0.001*** | 0.65 |
| | Time | 11118 | 5558.8 | 4.489 | 2, 191.314 | < 0.02* | 0.04 |
| | Frequency | 3480 | 1160.1 | 0.937 | 3, 189.815 | 0.424 | 0.01 |
| | Group x Time | 3617 | 904.3 | 0.730 | 4, 191.152 | 0.572 | 0.02 |
| | Group x Frequency | 189326 | 31554.4 | 25.482 | 6, 189.815 | < 0.001*** | 0.45 |
| | Time x Frequency | 9127 | 1521.1 | 1.228 | 6, 189.815 | 0.293 | 0.04 |
| | Group x Time x Frequency | 7936 | 661.3 | 0.534 | 12, 189.815 | 0.891 | 0.03 |

**Simple effects analysis conducted for Group x Frequency interaction**

**Group = CS+15**

| Contrast | Estimate | SE | df | t.ratio | p |
|---|---|---|---|---|---|
| 3 vs 7 | -45.634 | 11.9 | 230 | -3.820 | < 0.0005*** |
| 3 vs 11 | -87.231 | 11.9 | 230 | -7.302 | < 0.0001*** |
| 3 vs 15 | -103.651 | 11.9 | 230 | -8.677 | < 0.0001*** |
| 7 vs 11 | -41.598 | 11.9 | 230 | -3.482 | < 0.0012** |
| 7 vs 15 | -58.018 | 11.9 | 230 | -4.857 | < 0.0001*** |
| 11 vs 15 | -16.420 | 11.9 | 230 | -1.375 | 0.1706 |

**Group = CS+3**

| Contrast | Estimate | SE | df | t.ratio | p |
|---|---|---|---|---|---|
| 3 vs 7 | 59.286 | 14.1 | 230 | 4.194 | < 0.0002*** |
| 3 vs 11 | 81.162 | 14.1 | 230 | 5.742 | < 0.0001*** |
| 3 vs 15 | 80.288 | 14.1 | 230 | 5.680 | < 0.0001*** |
| 7 vs 11 | 21.876 | 14.1 | 230 | 1.548 | 0.3692 |
| 7 vs 15 | 21.002 | 14.1 | 230 | 1.486 | 0.3692 |
| 11 vs 15 | -0.874 | 14.1 | 230 | -0.062 | 0.9508 |

**Group = No shock control**

| Contrast | Estimate | SE | df | t.ratio | p |
|---|---|---|---|---|---|
| 3 vs 7 | -15.841 | 13.9 | 230 | -1.137 | 1.0000 |
| 3 vs 11 | -10.626 | 13.9 | 230 | -0.762 | 1.0000 |
| 3 vs 15 | -7.660 | 13.9 | 230 | -0.550 | 1.0000 |
| 7 vs 11 | 5.216 | 13.9 | 230 | 0.374 | 1.0000 |
| 7 vs 15 | 8.181 | 13.9 | 230 | 0.587 | 1.0000 |
| 11 vs 15 | 2.966 | 13.9 | 230 | 0.213 | 1.0000 |

#### b. Negative modulated

| Modulation | Variable | Sum Sq. | Mean Sq. | F | df | p | ηp² |
|---|---|---|---|---|---|---|---|
| Negative | Group | 5701.6 | 2850.8 | 2.2670 | 2, 18.142 | 0.1322 | 0.200 |
| | Time | 963.6 | 481.8 | 0.3831 | 2, 189.038 | 0.6822 | 0.004 |
| | Frequency | 6148.4 | 2049.5 | 1.6298 | 3, 186.216 | 0.1840 | 0.030 |
| | Group x Time | 1204.1 | 301.0 | 0.2394 | 4, 188.686 | 0.9158 | 0.005 |
| | Group x Frequency | 14708.2 | 2451.4 | 1.9494 | 6, 186.203 | 0.0750 | 0.059 |
| | Time x Frequency | 23070.6 | 3845.1 | 3.0577 | 6, 186.257 | < 0.0071** | 0.090 |
| | Group x Time x Frequency | 23011.2 | 1917.6 | 1.5249 | 12, 186.238 | 0.1182 | 0.089 |

**Simple effect analysis conducted for Time x Frequency interaction**

**Time: Day 1**

| Contrast | Estimate | SE | df | t.ratio | p |
|---|---|---|---|---|---|
| 3 vs 7 | 14.787 | 13.2 | 226 | -1.124 | 1.0000 |
| 3 vs 11 | -15.219 | 13.7 | 227 | -1.109 | 1.0000 |
| 3 vs 15 | -8.945 | 13.2 | 226 | -0.680 | 1.0000 |
| 7 vs 11 | -0.433 | 13.7 | 227 | -0.032 | 1.0000 |
| 7 vs 15 | 5.842 | 13.2 | 226 | 0.444 | 1.0000 |
| 11 vs 15 | -6.274 | 13.7 | 227 | 0.457 | 1.0000 |

**Time: Day 15**

| Contrast | Estimate | SE | df | t.ratio | p |
|---|---|---|---|---|---|
| 3 vs 7 | -14.825 | 13.7 | 227 | -1.080 | 1.0000 |
| 3 vs 11 | 9.416 | 13.2 | 226 | 0.716 | 1.0000 |
| 3 vs 15 | 3.458 | 13.2 | 226 | 0.263 | 1.0000 |
| 7 vs 11 | 24.241 | 13.7 | 227 | 1.766 | 0.4726 |
| 7 vs 15 | 18.283 | 13.7 | 227 | 1.332 | 0.9212 |
| 11 vs 15 | 5.958 | 13.2 | 226 | -0.453 | 1.0000 |

**Time: Day 30**

| Contrast | Estimate | SE | df | t.ratio | p |
|---|---|---|---|---|---|
| 3 vs 7 | -15.841 | 13.9 | 230 | -1.137 | 1.0000 |
| 3 vs 11 | -10.626 | 13.9 | 230 | -0.762 | 1.0000 |
| 3 vs 15 | -7.660 | 13.9 | 230 | -0.550 | 1.0000 |
| 7 vs 11 | 5.216 | 13.9 | 230 | 0.374 | 1.0000 |
| 7 vs 15 | 8.181 | 13.9 | 230 | 0.587 | 1.0000 |
| 11 vs 15 | 2.966 | 13.9 | 230 | 0.213 | 1.0000 |

**Table S2. Statistics comparing areas under the curve for the CS+15, CS+3, and No shock control groups across all frequencies and testing sessions.** (a-b) Three-way mixed model ANOVAs using Group (CS+15, CS+3, No shock), Frequency (3, 7, 11, 15 kHz), and Time (days 1, 15, 30) as variables for positive (a) and negative (b) cell responders. Simple effects were calculated using Holm correction.

**Table S3.**
**Fig. S4, S5, S6**

| Group | Type | Modulation | Test | Statistic | df | p | ηp² |
|---|---|---|---|---|---|---|---|
| CS+15 | Consistent | Positive | Test Day 1 | F = 6.11 | 3, 18 | 0.005 ** | 0.51 |
| | | | Test Day 15 | F = 6.29 | 3, 18 | 0.004** | 0.51 |
| | | | Test Day 30 | F = 4.43 | 3, 18 | 0.017* | 0.43 |
| | | Negative | Test Day 1 | F = 3.76 | 3, 18 | 0.030* | 0.39 |
| | | | Test Day 15 | F = 0.31 | 3, 18 | 0.817 | 0.05 |
| | | | Test Day 30 | F = 1.80 | 3, 18 | 0.183 | 0.23 |
| | Emerged Retained | Positive | Test Day 1 | F = 3.18 | 3, 13 | 0.060 | 0.42 |
| | | | Test Day 15 | F = 3.24 | 3, 18 | 0.046 * | 0.35 |
| | | | Test Day 30 | $\chi^2$ = 8.66 | 3 | 0.0.4* | NA |
| | | Negative | Test Day 1 | F = 1.89 | 3, 17 | 0.170 | 0.25 |
| | | | Test Day 15 | F = 1.45 | 3, 18 | 0.263 | 0.20 |
| | | | Test Day 30 | F = 2.63 | 3, 18 | 0.308 | 0.31 |
| | Transiently Active | Positive | Test Day 1 | F = 1.18 | 3, 17 | 0.332 | 0.17 |
| | | | Test Day 15 | F = 11.17 | 3, 18 | <0.001*** | 0.65 |
| | | | Test Day 30 | F = 3.87 | 3, 18 | 0.030 * | 0.39 |
| | | Negative | Test Day 1 | F = 0.39 | 3, 17 | 0.764 | 0.06 |
| | | | Test Day 15 | F = 0.30 | 3, 18 | 0.827 | 0.05 |
| | | | Test Day 30 | F = 2.08 | 3, 18 | 0.139 | 0.26 |
| CS+3 | Consistent | Positive | Test Day 1 | F = 9.32 | 3, 12 | 0.002 ** | 0.70 |
| | | | Test Day 15 | F = 3.18 | 3, 12 | 0.063 | 0.44 |
| | | | Test Day 30 | F = 5.39 | 3, 12 | 0.014 * | 0.57 |
| | | Negative | Test Day 1 | F = 1.28 | 3, 12 | 0.326 | 0.24 |
| | | | Test Day 15 | F = 0.53 | 3, 12 | 0.668 | 0.12 |
| | | | Test Day 30 | F = 0.47 | 3, 12 | 0.709 | 0.11 |
| | Emerged Retained | Positive | Test Day 1 | F = 1.49 | 3, 6 | 0.309 | 0.43 |
| | | | Test Day 15 | F = 6.18 | 3, 9 | 0.014 * | 0.67 |
| | | | Test Day 30 | F = 9.51 | 3, 9 | 0.004 ** | 0.76 |
| | | Negative | Test Day 1 | F = 4.75 | 3, 8 | 0.035 | 0.64 |
| | | | Test Day 15 | F = 0.65 | 3, 12 | 0.595 | 0.14 |
| | | | Test Day 30 | $\chi^2$ = 1.39 | 3 | 0.356 | NA |
| | Transiently Active | Positive | Test Day 1 | F = 0.28 | 3, 8 | 0.839 | 0.10 |
| | | | Test Day 15 | F = 6.45 | 3, 12 | 0.008 ** | 0.62 |
| | | | Test Day 30 | F = 5.49 | 3, 12 | 0.013 * | 0.58 |
| | | Negative | Test Day 1 | F = 1.12 | 3, 9 | 0.398 | 0.27 |
| | | | Test Day 15 | F = 1.57 | 3, 11 | 0.253 | 0.30 |
| | | | Test Day 30 | F = 0.426 | 3, 12 | 0.738 | 0.10 |

**Table S3. Statistics corresponding to consistently active, emerging-retained, and transiently active cells.** One-way ANOVAs with repeated measures were used when normality and equal variance tests were passed to test the effect of frequency. Friedman tests were used ($\chi^2$ with Kendall's W) when assumptions were violated, Friedman tests were used ($\chi^2$ with Kendall's W).

### Table S4.
### Fig. 4

#### Positive modulated cells

**Fig. 4a**

| Modulation | Variable | Sum Sq. | Mean Sq. | F | df | p | ηp² |
|---|---|---|---|---|---|---|---|
| Positive | Group | 8959 | 4.48 | 9.14 | 2, 15 | 0.003** | 0.55 |
| | Frequency | 7451 | 0.49 | 2.12 | 3, 45 | 0.111 | 0.12 |
| | Group x Frequency | 4859 | 0.81 | 8.69 | 6, 45 | < 0.001*** | 0.54 |

**Tukey post hoc comparisons**

| Tone | Comparison | Diff. of means | df | q | p |
|---|---|---|---|---|---|
| 3 kHz | CS+3 vs. No shock | 0.747 | 3 | 3.975 | 0.022* |
| | CS+3 vs. CS+15 | 0.491 | 3 | 2.706 | 0.151 |
| | CS+15 vs. No shock | 0.255 | 3 | 1.479 | 0.554 |
| 7 kHz | CS+15 vs. No shock | 0.684 | 3 | 3.963 | 0.022* |
| | CS+15 vs. CS+3 | 0.229 | 3 | 1.259 | 0.650 |
| | CS+3 vs. No shock | 0.455 | 3 | 2.423 | 0.215 |
| 11 kHz | CS+15 vs. No shock | 1.174 | 3 | 6.805 | < 0.001*** |
| | CS+15 vs. CS+3 | 0.737 | 3 | 4.058 | 0.019* |
| | CS+3 vs. No shock | 0.437 | 3 | 2.328 | 0.241 |
| 15 kHz | CS+15 vs. No shock | 1.215 | 3 | 7.040 | < 0.001*** |
| | CS+15 vs. CS+3 | 1.176 | 3 | 6.476 | < 0.001*** |
| | CS+3 vs. No shock | 0.0387 | 3 | 0.206 | 0.988 |

**Fig. 4b**

| Modulation | Variable | Sum Sq. | Mean Sq. | F | df | p | ηp² |
|---|---|---|---|---|---|---|---|
| positive | Group | 0.26 | 0.13 | 2.79 | 2, 15 | 0.094 | 0.27 |
| | Day | 0.02 | 0.007 | 0.806 | 3, 45 | 0.497 | 0.05 |
| | Group x Day | 0.154 | 0.025 | 2.84 | 6, 45 | 0.020* | 0.27 |

**Tukey post hoc comparisons**

| Tone | Comparison | Diff. of means | df | q | p |
|---|---|---|---|---|---|
| 3 kHz | No shock vs. CS+3 | 0.164 | 3 | 2.818 | 0.130 |
| | No shock vs. CS+15 | 0.0454 | 3 | 0.849 | 0.821 |
| | CS+15 vs. CS+3 | 0.119 | 3 | 2.107 | 0.308 |
| 7 kHz | No shock vs. CS+15 | 0.166 | 3 | 3.099 | 0.087 |
| | No shock vs. CS+3 | 0.105 | 3 | 1.800 | 0.420 |
| | CS+3 vs. CS+15 | 0.061 | 3 | 1.084 | 0.726 |
| 11 kHz | No shock vs. CS+15 | 0.185 | 3 | 3.455 | 0.051 |
| | No shock vs. CS+3 | 0.0418 | 3 | 0.718 | 0.868 |
| | CS+3 vs. CS+15 | 0.143 | 3 | 2.539 | 0.186 |
| 15 kHz | No shock vs. CS+15 | 0.172 | 3 | 3.207 | 0.074 |
| | No shock vs. CS+3 | 0.000427 | 3 | 0.00734 | 1.000 |
| | CS+3 vs. CS+15 | 0.171 | 3 | 3.040 | 0.095 |

**Fig. 4c**

| Modulation | Variable | Sum Sq. | Mean Sq. | F | df | p | ηp² |
|---|---|---|---|---|---|---|---|
| Positive | Group | 0.400 | 0.200 | 3.802 | 2, 15 | 0.046* | 0.336 |
| | Condition | 3.983 | 3.983 | 176.813 | 1, 15 | < 0.001*** | 0.922 |
| | Group x Condition | 0.267 | 0.133 | 5.921 | 2, 15 | 0.013* | 0.441 |

**Tukey post hoc comparisons**

| Factor | Comparison | Diff. of means | df | q | p |
|---|---|---|---|---|---|
| within CS+15 | Tone vs. freezing | 0.862 | 2 | 15.238 | < 0.001*** |
| within CS+3 | Tone vs. freezing | 0.696 | 2 | 10.361 | < 0.001*** |
| within Nonshock | Tone vs. freezing | 0.457 | 2 | 7.429 | < 0.001*** |
| within freezing | CS+15 vs. Nonshock | 0.045 | 3 | 0.684 | 0.880 |
| | CS+15 vs. CS+3 | 0.054 | 3 | 0.779 | 0.848 |

| | | | | | |
|---|---|---|---|---|---|
| | Nonshock vs. CS+3 | 0.009 | 3 | 0.125 | 0.996 |
| within tone | CS+15 vs. Nonshock | 0.451 | 3 | 5.271 | 0.005** |
| | CS+15 vs. CS+3 | 0.221 | 3 | 2.461 | 0.223 |
| | CS+3 vs. Nonshock | 0.230 | 3 | 2.483 | 0.218 |

**Fig. 4d**

| Modulation | Variable | Sum Sq. | Mean Sq. | F | df | p | ηp² |
|---|---|---|---|---|---|---|---|
| Positive | Group | 156.357 | 78.178 | 1.166 | 2, 15 | 0.336 | 0.13 |
| | Day | 17.999 | 68.24 | 0.174 | 2, 26 | 0.842 | 0.01 |
| | Group x Day | 106.236 | 26.559 | 0.512 | 4, 26 | 0.727 | 0.07 |

### Negative modulated cells

**Fig. 4e**

| Modulation | Variable | Sum Sq. | Mean Sq. | F | df | p | ηp² |
|---|---|---|---|---|---|---|---|
| Negative | Group | 0.135 | 0.0677 | 0.968 | 2, 15 | 0.402 | 0.11 |
| | Frequency | 0.101 | 0.0336 | 1.453 | 3, 44 | 0.240 | 0.09 |
| | Group x Frequency | 0.072 | 0.012 | 0.519 | 6, 44 | 0.791 | 0.07 |

**Fig. 4f**

| Modulation | Variable | Sum Sq. | Mean Sq. | F | df | p | ηp² |
|---|---|---|---|---|---|---|---|
| Negative | Group | 1.038 | 0.519 | 3.066 | 2, 15 | 0.076 | 0.29 |
| | Frequency | 0.141 | 0.0471 | 0.399 | 3, 44 | 0.754 | 0.03 |
| | Group x Frequency | 0.468 | 0.0779 | 0.661 | 6, 44 | 0.681 | 0.08 |

**Fig. 4g**

| Modulation | Variable | Sum Sq. | Mean Sq. | F | df | p | ηp² |
|---|---|---|---|---|---|---|---|
| Negative | Group | 0.004 | 0.002 | 0.061 | 2, 15 | 0.941 | 0.008 |
| | Condition | 0.308 | 0.308 | 32.496 | 1, 15 | < 0.001*** | 0.684 |
| | Group x Condition | 0.101 | 0.050 | 5.315 | 2, 15 | 0.018* | 0.415 |

**Tukey post hoc comparisons**

| Factor | Comparison | Diff. of means | df | q | p |
|---|---|---|---|---|---|
| within CS+15 | Tone vs. freezing | 0.302 | 2 | 8.213 | < 0.001*** |
| within CS+3 | Tone vs. freezing | 0.206 | 2 | 4.699 | 0.005** |
| within Nonshock | Tone vs. freezing | 0.053 | 2 | 1.338 | 0.359 |
| within freezing | Nonshock vs. CS+15 | 0.100 | 3 | 1.473 | 0.563 |
| | Nonshock vs. CS+3 | 0.070 | 3 | 0.943 | 0.786 |
| | CS+3 vs. CS+15 | 0.030 | 3 | 0.420 | 0.953 |
| within tone | CS+15 vs. Nonshock | 0.149 | 3 | 3.145 | 0.099 |
| | CS+15 vs. CS+3 | 0.066 | 3 | 1.315 | 0.631 |
| | CS+3 vs. Nonshock | 0.083 | 3 | 1.608 | 0.507 |

**Fig. 4h**

| Modulation | Variable | Sum Sq. | Mean Sq. | F | df | p | ηp² |
|---|---|---|---|---|---|---|---|
| Negative | Group | 886.884 | 443.442 | 2.756 | 2, 15 | 0.092 | 0.27 |
| | Day | 400.272 | 200.136 | 1.500 | 2, 26 | 0.242 | 0.10 |
| | Group x Day | 468.189 | 117.047 | 0.877 | 4, 26 | 0.491 | 0.12 |

**Table S4.** Statistical analysis of GLM-derived measures for cells used in the population curves shown in Fig. 4. Two-way ANOVAs with repeated measures were conducted to evaluate effects of group (CS+15, CS+3, or No shock), frequency (3, 7, 11, 15 kHz) or day (days 1, 15, 30), and their interactions. Tukey post hoc comparisons are shown where simple effects were tested.

**Table S5.**
**Fig. 6 and 7**

| Group | Cluster | Mod. | % Stable D1-D15 | P (D1-D15) | % Stable D15-D30 | P (D15-D30) | % Stable D1-D30 | P (D1-D30) |
|---|---|---|---|---|---|---|---|---|
| CS+15 Fig. 6 | Single tone c1 | + | 13.5 | 0.391 | 5.0 | 1.000 | 9.3 | 0.870 |
| | Single tone c2 | + | 3.4 | 1.000 | 7.1 | 1.000 | 8.2 | 1.000 |
| | Single tone c3 | + | 11.1 | 0.576 | 13.4 | 0.391 | 19.2 | 0.124 |
| | Single tone c4 | + | 15.5 | 0.391 | 15.1 | 0.391 | 20.9 | 0.121 |
| | Single tone d1 | - | 19.3 | 0.079 | 11.9 | 0.391 | 12.9 | 0.576 |
| | Single tone d2 | - | 10.4 | 0.765 | 10.0 | 0.661 | 10.3 | 0.576 |
| | Single tone d3 | - | 11.9 | 0.697 | 20.2 | 0.334 | 17.8 | 0.697 |
| | Single tone d4 | - | 20.2 | 0.011* | 11.8 | 1.000 | 10.3 | 1.000 |
| Fig. 7 | Graded a | | 28.2 | 0.000*** | 22.8 | 0.000*** | 21.3 | 0.001*** |
| | Graded b | | 11.1 | 0.034* | 5.5 | 0.576 | 5.9 | 0.576 |
| CS+3 Fig. 6 | Single tone c1 | + | 23.9 | 0.014* | 16.5 | 0.460 | 19.0 | 0.476 |
| | Single tone c2 | + | 13.4 | 0.708 | 6.5 | 0.982 | 11.9 | 0.792 |
| | Single tone c3 | + | 10.8 | 0.852 | 9.8 | 0.939 | 21.1 | 0.185 |
| | Single tone c4 | + | 10.4 | 0.756 | 7.7 | 0.982 | 19.6 | 0.521 |
| | Single tone d1 | - | 20.9 | 0.230 | 20.6 | 0.074 | 13.0 | 0.708 |
| | Single tone d2 | - | 11.5 | 0.714 | 8.0 | 0.579 | 8.6 | 0.592 |
| | Single tone d3 | - | 9.7 | 0.739 | 5.3 | 0.982 | 10.3 | 0.846 |
| | Single tone d4 | - | 8.9 | 0.872 | 7.6 | 0.972 | 12.8 | 0.592 |
| Fig. 7 | Graded c | | 21.2 | 0.008** | 17.9 | 0.026* | 25.0 | 0.008** |
| | Graded d | | 9.1 | 0.303 | 40.0 | 0.003** | 14.3 | 0.303 |
| | Graded e | | 12.9 | 0.047* | 15.4 | 0.047* | 8.3 | 0.589 |
| No Shock Fig. 6 | Single tone c1 | + | 7.8 | 1.000 | 8.3 | 1.000 | 22.2 | 1.000 |
| | Single tone c2 | + | 11.5 | 1.000 | 10.0 | 1.000 | 0.0 | 1.000 |
| | Single tone c3 | + | 21.6 | 1.000 | 16.7 | 1.000 | 40.0 | 1.000 |
| | Single tone c4 | + | 11.5 | 1.000 | 5.9 | 1.000 | 0.0 | 1.000 |
| | Single tone d1 | - | 10.0 | 1.000 | 7.1 | 1.000 | 0.0 | 1.000 |
| | Single tone d2 | - | 5.5 | 1.000 | 21.4 | 1.000 | 21.4 | 1.000 |
| | Single tone d3 | - | 12.0 | 1.000 | 4.0 | 1.000 | 33.3 | 1.000 |
| | Single tone d4 | - | 6.4 | 1.000 | 22.2 | 1.000 | 0.0 | 1.000 |

**Table S5.** Statistical analyses identifying neuronal clusters that remained stable over time, defined as neurons that retained the same functional response profile on days 15 and 30 as on day 1. These analyses correspond to the clusters shown in Figs. 6 and 7, as indicated in the table. For each cluster, the observed proportion of neurons that preserved their response profile was compared with a null distribution, with multiple comparisons controlled using the Benjamini–Hochberg procedure. Only the graded neuronal clusters in each experimental group (Fig. 7) exhibited significant stability across time. No graded clusters were identified in the no-shock control group.

**Table S6**
**Fig. S10.**

| Cluster | Groups | Day | Statistic | df | p | ηp² |
|---|---|---|---|---|---|---|
| **Statistics for clusters responding to individual frequencies (Fig. 6c)** | | | | | | |
| Fig. 6c, c.1<br>Fig. S7, S8, S9<br>Maximal response<br>at 3 kHz | CS+15<br>CS+3<br>No shock | Test Day 1 | Group: F =1.49 | 2, 15 | 0.258 | 0.17 |
| | | | Frequency: F=30.51 | 3, 6 | 0.001***. | 0.94 |
| | | | Interaction: 4.50 | 6, 45 | 0.001***. | 0.38 |
| | | Test Day 15 | Group: F =4.59 | 2, 15 | 0.03* | 0.38 |
| | | | Frequency: F=31.99 | 3, 6 | 0.001***. | 0.94 |
| | | | Interaction: 2.28 | 6,4 5 | 0.053. | 0.23 |
| | | Test Day 30 | Group: F =0.36 | 3, 13 | 0.705 | 0.08 |
| | | | Frequency: F=30.02 | 3, 6 | 0.001***. | 0.94 |
| | | | Interaction: 2.78 | 6, 39 | 0.024*. | 0.30 |
| Fig. 6c, c.2<br>Fig. S7, S8, S9<br>Maximal response<br>at 7kHz | CS+15<br>CS+3<br>No shock | Test Day 1 | Group: F =2.29 | 2,15 | 0.135 | 0.23 |
| | | | Frequency: F=30.18 | 3, 6 | 0.001***. | 0.94 |
| | | | Interaction: 2.62 | 6, 45 | 0.029*. | 0.26 |
| | | Test Day 15 | Group: F =3.50 | 2, 15 | 0.057 | 0.32 |
| | | | Frequency: F=27.46 | 3, 6 | 0.001***. | 0.93 |
| | | | Interaction: 0.99 | 6, 45 | 0.441. | 0.12 |
| | | Test Day 30 | Group: F =3.51 | 2,13 | 0.061 | 0.35 |
| | | | Frequency: F=24.64 | 3, 6 | 0.001***. | 0.93 |
| | | | Interaction: 1.057 | 6, 3 9 | 0.404. | 0.14 |
| Fig. 6c, c.3<br>Fig. S7, S8, S9<br>Maximal response<br>at 11 kHz | CS+15<br>CS+3<br>No shock | Test Day 1 | Group: F =1.45 | 2, 15 | 0.266 | 0.16 |
| | | | Frequency: F=20.44 | 3, 6 | 0.001***. | 0.91 |
| | | | Interaction: 0.912 | 6, 4 5 | 0.495. | 0.11 |
| | | Test Day 15 | Group: F =15.47 | 2, 15 | 0.001*** | 0.67 |
| | | | Frequency: F=38.53 | 3, 6 | 0.001***. | 0.95 |
| | | | Interaction: 3.40 | 6,4 5 | 0.007**. | 0.31 |
| | | Test Day 30 | Group: F =11.46 | 2, 13 | 0.001*** | 0.64 |
| | | | Frequency: F=14.20 | 3, 6 | 0.001***. | 0.88 |
| | | | Interaction: 6.12 | 6, 3 9 | 0.001***. | 0.49 |
| Fig. 6c, c.4<br>Fig. S7, S8, S9<br>Maximal response<br>at 15 kHz | CS+15<br>CS+3<br>No shock | Test Day 1 | Group: F =2.71 | 3, 15 | 0.099 | 0.35 |
| | | | Frequency: F=22.28 | 3, 6 | 0.001***. | 0.92 |
| | | | Interaction: 0.38 | 6,4 5 | 0.89. | 0.05 |
| | | Test Day 15 | Group: F =7.64 | 2,1 5 | 0.005** | 0.51 |
| | | | Frequency: F=33.86 | 3, 6 | 0.001***. | 0.94 |
| | | | Interaction: 10.58 | 6, 45 | 0.001***. | 0.59 |
| | | Test Day 30 | Group: F =8.01 | 2, 13 | 0.005** | 0.55 |
| | | | Frequency: F=30.05 | 3, 6 | 0.001***. | 0.94 |
| | | | Interaction: 4.15 | 6, 39 | 0.003**. | 0.39 |
| **Statistics for clusters showing graded valence (Fig.7)** | | | | | | |
| Graded cluster<br>Fig. 7a<br>Fig. S7a | Experimental<br>CS+15 | Test Day 1,<br>15, and 30 | Day: F =2.89 | 2, 12 | 0.10 | 0.33 |
| | | | Frequency: F=29.28 | 3, 12 | 0.001***. | 0.88 |
| | | | Interaction: 0.351 | 6,36 | 0.905. | 0.06 |
| Graded cluster<br>Fig. 7c<br>Fig. S8a | Experimental<br>CS+3 | Test Day 1,<br>15, and 30 | Day F =0.467 | 2, 8 | 0.643 | 0.11 |
| | | | Frequency: F=53.37 | 3,12 | 0.001***. | 0.93 |
| | | | Interaction: 2.63 | 6, 24 | 0.042*. | 0.40 |

**Table S6.** Statistical analyses corresponding to the normalized baseline-to-stimulus firing rate (BSR) for the neuronal clusters shown in Figs. S7–S9. BSRs for the clusters shown in Fig. S7-S9 are presented in Fig. S10, whereas BSRs for the graded clusters shown in Figs. S7-S8 are presented in Figs. 7h–i. For pure tone-selective clusters, two-way repeated-measures ANOVAs were used to evaluate the effects of group (CS+15, CS+3, or No Shock), frequency (3, 7, 11, and 15 kHz), and their interaction. For graded clusters, one-way repeated-measures ANOVAs were used to evaluate the effect of time. Tukey's multiple-comparisons tests are reported in Figs. S10 and 7h–i.

**Table 7.**
**Fig. 8**

### Positive modulated single-frequency neurons

#### Fig. 8a

| Modulation | Variable | Sum Sq. | Mean Sq. | F | df | p | ηp² |
|---|---|---|---|---|---|---|---|
| Positive | Group | 15.905 | 7.952 | 19.447 | 2, 15 | < 0.001*** | 0.72 |
| | Frequency | 0.832 | 0.277 | 1.108 | 3, 45 | 0.356 | 0.07 |
| | Group x Frequency | 6.319 | 1.053 | 4.21 | 6, 45 | 0.002** | 0.36 |

**Tukey post hoc comparisons**

| Tone | Comparison | Diff. of means | df | q | p |
|---|---|---|---|---|---|
| 3 kHz | CS+3 vs. No shock | 0.757 | 3 | 3.284 | 0.061 |
| | CS+3 vs. CS+15 | 0.374 | 3 | 1.677 | 0.467 |
| | CS+15 vs. No shock | 0.383 | 3 | 1.810 | 0.412 |
| 7 kHz | CS+15 vs. No shock | 0.969 | 3 | 4.574 | < 0.006** |
| | CS+15 vs. CS+3 | 0.341 | 3 | 1.530 | 0.529 |
| | CS+3 vs. No shock | 0.628 | 3 | 2.723 | 0.141 |
| 11 kHz | CS+15 vs. No shock | 1.609 | 3 | 7.597 | < 0.001*** |
| | CS+15 vs. CS+3 | 1.363 | 3 | 6.116 | < 0.001*** |
| | CS+3 vs. No shock | 0.246 | 3 | 1.065 | 0.733 |
| 15 kHz | CS+15 vs. No shock | 1.443 | 3 | 6.811 | < 0.001*** |
| | CS+15 vs. CS+3 | 1.218 | 3 | 5.464 | < 0.001*** |
| | CS+3 vs. No shock | 0.225 | 3 | 0.975 | 0.771 |

#### Fig. 8b

| Modulation | Variable | Sum Sq. | Mean Sq. | F | df | p | ηp² |
|---|---|---|---|---|---|---|---|
| Positive | Group | 0.207 | 0.104 | 6.62 | 2, 15 | 0.008** | 0.47 |
| | Day | 0.0178 | 0.0089 | 1.205 | 2, 26 | 0.316 | 0.08 |
| | Group x Day | 0.0379 | 0.00948 | 1.281 | 4, 26 | 0.303 | 0.16 |

**Tukey post hoc comparisons**

| Comparison | Diff. of means | df | q | p |
|---|---|---|---|---|
| No shock vs. CS+15 | 0.168 | 3 | 5.026 | < 0.008** |
| No shock vs. CS+3 | 0.117 | 3 | 3.316 | 0.080 |
| CS+3 vs. CS+15 | 0.0513 | 3 | 1.654 | 0.488 |

#### Fig. 8c

| Modulation | Variable | Sum Sq. | Mean Sq. | F | df | p | ηp² |
|---|---|---|---|---|---|---|---|
| Positive | Group | 0.49 | 0.245 | 5.277 | 2, 15 | 0.018* | 0.41 |
| | Condition | 3.962 | 3.962 | 254.641 | 1, 15 | < 0.001*** | 0.94 |
| | Group x Condition | 0.318 | 0.159 | 10.205 | 2, 15 | 0.002** | 0.58 |

**Tukey post hoc comparisons**

| Factor | Comparison | Diff. of means | df | q | p |
|---|---|---|---|---|---|
| within CS+15 | Tone vs. freezing | 0.899 | 2 | 19.078 | < 0.001*** |
| within CS+3 | Tone vs. freezing | 0.651 | 2 | 11.676 | < 0.001*** |

| | | | | | |
|---|---|---|---|---|---|
| within No shock | Tone vs. freezing | 0.459 | 2 | 9.005 | < 0.001*** |
| within freezing | CS+15 vs. No shock | 0.0531 | 3 | 0.767 | 0.851 |
| | CS+15 vs. CS+3 | 0.0317 | 3 | 0.435 | 0.949 |
| | CS+3 vs. No shock | 0.0214 | 3 | 0.284 | 0.978 |
| within tone | CS+15 vs. No shock | 0.494 | 3 | 7.134 | < 0.001*** |
| | CS+15 vs. CS+3 | 0.280 | 3 | 3.839 | 0.031* |
| | CS+3 vs. No shock | 0.214 | 3 | 2.843 | 0.131 |

#### Fig. 8d

| Modulation | Variable | Sum Sq. | Mean Sq. | F | df | p | ηp² |
|---|---|---|---|---|---|---|---|
| Positive | Group | 83.099 | 41.549 | 0.763 | 2, 15 | 0.482 | 0.09 |
| | Day | 62.468 | 31.234 | 0.649 | 2, 26 | 0.531 | 0.05 |
| | Group x Day | 130.193 | 32.548 | 0.677 | 4, 26 | 0.614 | 0.09 |

### Negative modulated single-frequency neurons

#### Fig. 8e

| Modulation | Variable | Sum Sq. | Mean Sq. | F | df | p | ηp² |
|---|---|---|---|---|---|---|---|
| Negative | Group | 1.765 | 0.882 | 11.246 | 2, 15 | 0.001*** | 0.60 |
| | Frequency | 0.121 | 0.0404 | 1.274 | 3, 45 | 0.295 | 0.08 |
| | Group x Frequency | 0.166 | 0.0277 | 0.872 | 6, 45 | 0.523 | 0.10 |

#### Tukey post hoc comparisons

| Comparison | Diff. of means | df | q | p |
|---|---|---|---|---|
| No shock vs. CS+15 | 0.365 | 3 | 6.624 | < 0.001*** |
| No shock vs. CS+3 | 0.142 | 3 | 2.361 | 0.249 |
| CS+3 vs. CS+15 | 0.223 | 3 | 3.852 | 0.039* |

#### Fig. 8f

| Modulation | Variable | Sum Sq. | Mean Sq. | F | df | p | ηp² |
|---|---|---|---|---|---|---|---|
| Negative | Group | 1.474 | 0.737 | 18.566 | 2, 15 | < 0.001*** | 0.71 |
| | Frequency | 0.131 | 0.0656 | 1.509 | 2, 26 | 0.240 | 0.10 |
| | Group x Frequency | 0.185 | 0.0462 | 1.062 | 4, 26 | 0.395 | 0.14 |

#### Tukey post hoc comparisons

| Comparison | Diff. of means | df | q | p |
|---|---|---|---|---|
| No shock vs. CS+15 | 0.451 | 3 | 8.650 | < 0.001*** |
| No shock vs. CS+3 | 0.284 | 3 | 5.176 | 0.006** |
| CS+3 vs. CS+15 | 0.167 | 3 | 3.450 | 0.067 |

#### Fig. 8g

| Modulation | Variable | Sum Sq. | Mean Sq. | F | df | p | ηp² |
|---|---|---|---|---|---|---|---|
| Negative | Group | 0.149 | 0.0746 | 4.47 | 2, 15 | 0.030* | 0.37 |
| | Day | 0.261 | 0.261 | 39.684 | 1, 15 | < 0.001*** | 0.73 |
| | Group x Day | 0.174 | 0.087 | 13.233 | 2, 15 | < 0.001*** | 0.64 |

#### Tukey post hoc comparisons

| Factor | Comparison | Diff. of means | df | q | p |
|---|---|---|---|---|---|

| | | | | | |
|---|---|---|---|---|---|
| within CS+15 | Tone vs. freezing | 0.343 | 2 | 11.207 | < 0.001*** |
| within CS+3 | Tone vs. freezing | 0.154 | 2 | 4.257 | < 0.009** |
| within No shock | Tone vs. freezing | 0.0178 | 2 | 0.538 | 0.709 |
| within freezing | No shock vs. CS+15 | 0.0125 | 3 | 0.294 | 0.977 |
| | No shock vs. CS+3 | 0.00849 | 3 | 0.184 | 0.991 |
| | CS+3 vs. CS+15 | 0.00397 | 3 | 0.0889 | 0.998 |
| within tone | CS+15 vs. No shock | 0.313 | 3 | 7.379 | < 0.001*** |
| | CS+15 vs. CS+3 | 0.185 | 3 | 4.144 | 0.019* |
| | CS+3 vs. No shock | 0.128 | 3 | 2.772 | 0.143 |

#### Fig. 8h

| Modulation | Variable | Sum Sq. | Mean Sq. | F | df | p | ηp² |
|---|---|---|---|---|---|---|---|
| Negative | Group | 1464.631 | 732.315 | 7.469 | 2, 15 | 0.005** | 0.50 |
| | Day | 177.73 | 88.865 | 1.756 | 2, 26 | 0.193 | 0.12 |
| | Group x Day | 1316.013 | 23.848 | 0.471 | 4, 26 | 0.756 | 0.07 |

#### Tukey post hoc comparisons

| Comparison | Diff. of means | df | q | p |
|---|---|---|---|---|
| No shock vs. CS+15 | 14.17 | 3 | 5.354 | < 0.005** |
| No shock vs. CS+3 | 8.058 | 3 | 2.890 | 0.136 |
| CS+3 vs. CS+15 | 6.112 | 3 | 2.492 | 0.216 |

### Positive modulated graded neurons

#### Fig. 8i

| Modulation | Variable | Sum Sq. | Mean Sq. | F | df | p | ηp² |
|---|---|---|---|---|---|---|---|
| Positive | Group | 0.449 | 0.449 | 29.231 | 1, 15 | < 0.001*** | 0.66 |
| | Frequency | 0.0709 | 0.977 | 0.977 | 3, 30 | 0.417 | 0.09 |
| | Group x Frequency | 3.933 | 1.311 | 54.221 | 3, 30 | < 0.001*** | 0.84 |

#### Tukey post hoc comparisons

| Tone | Comparison | Diff. of means | df | q | p |
|---|---|---|---|---|---|
| 3 kHz | CS+3 vs. CS+15 | 0.757 | 2 | 12.335 | < 0.001*** |
| 7 kHz | CS+15 vs. CS+3 | 0.215 | 2 | 3.495 | 0.018* |
| 11 kHz | CS+15 vs. CS+3 | 0.624 | 2 | 10.173 | < 0.001*** |
| 15 kHz | CS+15 vs. CS+3 | 0.702 | 2 | 11.446 | < 0.001*** |

#### Fig. 8j

| Modulation | Variable | Sum Sq. | Mean Sq. | F | df | p | ηp² |
|---|---|---|---|---|---|---|---|
| Positive | Group | 0.055 | 0.055 | 4.795 | 1, 10 | 0.053 | 0.32 |
| | Frequency | 0.0933 | 0.0467 | 6.548 | 2, 19 | 0.007** | 0.41 |
| | Group x Frequency | 0.0929 | 0.0464 | 6.518 | 2, 19 | 0.007** | 0.41 |

#### Tukey post hoc comparisons

| Factor | Comparison | Diff. of means | df | q | p |
|---|---|---|---|---|---|
| within CS+15 | 15 vs. 2 | 0.00771 | 3 | 0.228 | 0.986 |
| | 30 vs. 2 | 0.0194 | 3 | 0.575 | 0.914 |
| | 30 vs. 15 | 0.0117 | 3 | 0.368 | 0.964 |

| | | | | | |
|---|---|---|---|---|---|
| within CS+3 | 2 vs. 15 | 0.236 | 3 | 6.256 | < 0.001*** |
| | 2 vs. 30 | 0.0346 | 3 | 0.916 | 0.796 |
| | 30 vs. 15 | 0.202 | 3 | 5.340 | 0.004** |
| within 2 | CS+3 vs. CS+15 | 0.180 | 2 | 4.454 | 0.004** |
| within 15 | CS+15 vs. CS+3 | 0.0639 | 2 | 1.661 | 0.250 |
| within 30 | CS+3 vs. CS+15 | 0.126 | 2 | 3.275 | 0.028* |

#### Fig. 8k

| Modulation | Variable | Sum Sq. | Mean Sq. | F | df | p | ηp² |
|---|---|---|---|---|---|---|---|
| positive | Group | 0.779 | 0.779 | 4.761 | 1, 10 | 0.054 | 0.32 |
| | Condition | 17.748 | 17.748 | 221.136 | 1, 23 | < 0.001*** | 0.91 |
| | Group x Condition | 0.671 | 0.671 | 8.366 | 1, 23 | < 0.016* | 0.27 |

#### Tukey post hoc comparisons

| Factor | Comparison | Diff. of means | df | q | p |
|---|---|---|---|---|---|
| within CS+15 | Tone vs. freezing | 2.084 | 2 | 19.458 | < 0.001*** |
| within CS+3 | Tone vs. freezing | 1.405 | 2 | 11.090 | < 0.001*** |
| within freezing | CS+15 vs. CS+3 | 0.0261 | 2 | 0.181 | 0.900 |
| within tone | CS+15 vs. CS+3 | 0.705 | 2 | 4.874 | 0.003** |

#### Fig. 8l

| Modulation | Variable | Sum Sq. | Mean Sq. | F | df | p | ηp² |
|---|---|---|---|---|---|---|---|
| Positive | Group | 5.092 | 5.092 | 0.105 | 1, 10 | 0.752 | 0.01 |
| | Condition | 17.586 | 8.793 | 0.470 | 2, 19 | 0.632 | 0.05 |
| | Group x Condition | 20.386 | 10.193 | 0.545 | 2, 19 | 0.588 | 0.05 |

### Negative modulated graded neurons

#### Fig. 8m

| Modulation | Variable | Sum Sq. | Mean Sq. | F | df | p | ηp² |
|---|---|---|---|---|---|---|---|
| Negative | Group | 0.513 | 0.513 | 13.677 | 1, 10 | 0.004** | 0.58 |
| | Frequency | 0.00725 | 0.00242 | 0.0836 | 3, 30 | 0.968 | 0.01 |
| | Group x Frequency | 6.404 | 2.135 | 73.84 | 3, 30 | < 0.001*** | 0.88 |

#### Tukey post hoc comparisons

| Tone | Comparison | Diff. of means | df | q | p |
|---|---|---|---|---|---|
| 3 kHz | CS+15 vs. CS+3 | 1.006 | 2 | 13.781 | < 0.001*** |
| 7 kHz | CS+3 vs. CS+15 | 0.229 | 2 | 3.131 | 0.033* |
| 11 kHz | CS+3 vs. CS+15 | 0.776 | 2 | 10.628 | < 0.001*** |
| 15 kHz | CS+3 vs. CS+15 | 0.841 | 2 | 11.519 | < 0.001*** |

#### Fig. 8n

| Modulation | Variable | Sum Sq. | Mean Sq. | F | df | p | ηp² |
|---|---|---|---|---|---|---|---|
| Negative | Group | 0.00308 | 0.00308 | 0.122 | 1, 10 | 0.733 | 0.01 |
| | Day | 0.0437 | 0.0218 | 1.323 | 2, 15 | 0.296 | 0.15 |
| | Group x Day | 0.0299 | 0.015 | 0.907 | 2, 15 | 0.425 | 0.11 |

**Fig. 8o**

| Modulation | Variable | Sum Sq. | Mean Sq. | F | df | p | ηp² |
|---|---|---|---|---|---|---|---|
| Negative | Group | 0.0609 | 0.0609 | 1.421 | 1, 10 | 0.261 | 0.12 |
| | Day | 3.081 | 3.081 | 443.863 | 1, 10 | < 0.001*** | 0.98 |
| | Group x Day | 0.0436 | 0.0436 | 6.285 | 2, 15 | 0.031* | 0.46 |

**Tukey post hoc comparisons**

| Factor | Comparison | Diff. of means | df | q | p |
|---|---|---|---|---|---|
| within CS+15 | Tone vs. freezing | 0.813 | 2 | 25.825 | < 0.001*** |
| within CS+3 | Tone vs. freezing | 0.640 | 2 | 17.184 | < 0.001*** |
| within freezing | CS+15 vs. CS+3 | 0.0157 | 2 | 0.240 | 0.868 |
| within tone | CS+15 vs. CS+3 | 0.189 | 2 | 2.888 | 0.062 |

**Fig. 8p**

| Modulation | Variable | Sum Sq. | Mean Sq. | F | df | p | ηp² |
|---|---|---|---|---|---|---|---|
| Negative | Group | 174.242 | 174.242 | 2.001 | 1, 10 | 0.188 | 0.17 |
| | Day | 11.597 | 5.799 | 0.065 | 2, 19 | 0.937 | 0.01 |
| | Group x Day | 149.463 | 74.732 | 0.838 | 2, 19 | 0.448 | 0.08 |

**Table S7.** Statistical analysis of positive and negative modulated single-frequency and graded neurons shown in Fig. 8. Two-way ANOVAs with repeated measures were conducted to evaluate effects of group (CS+15, CS+3, or No shock), frequency (3, 7, 11, 15 kHz), day, condition (tone vs. freezing), and their interactions. Tukey post hoc comparisons are shown where simple effects were tested.
